# Evaluation of taxonomic classification and profiling methods for long-read shotgun metagenomic sequencing datasets

**DOI:** 10.1101/2022.01.31.478527

**Authors:** Daniel M. Portik, C. Titus Brown, N. Tessa Pierce-Ward

**Affiliations:** Pacific Biosciences, 1305 O’Brien Dr, Menlo Park, California 93025 USA; Department of Population Health and Reproduction, University of California Davis, Davis, California USA

**Keywords:** metagenomics, taxonomic classifier, taxonomic profiler, long reads, PacBio, Nanopore, mock community, benchmarking, sourmash

## Abstract

**Background:** Long-read shotgun metagenomic sequencing is gaining in popularity and offers many advantages over short-read sequencing. The higher information content in long reads is useful for a variety of metagenomics analyses, including taxonomic classification and profiling. The development of long-read specific tools for taxonomic classification is accelerating, yet there is a lack of information regarding their relative performance. Here, we perform a critical benchmarking study using 11 methods, including five methods designed specifically for long reads. We applied these tools to several mock community datasets generated using Pacific Biosciences (PacBio) HiFi or Oxford Nanopore Technology (ONT) sequencing, and evaluated their performance based on read utilization, detection metrics, and relative abundance estimates.

**Results:** Our results show that long-read classifiers generally performed best. Several short-read classification and profiling methods produced many false positives (particularly at lower abundances), required heavy filtering to achieve acceptable precision (at the cost of reduced recall), and produced inaccurate abundance estimates. By contrast, two long-read methods (BugSeq, MEGAN-LR & DIAMOND) and one generalized method (sourmash) displayed high precision and recall without any filtering required. Furthermore, in the PacBio HiFi datasets these methods detected all species down to the 0.1% abundance level with high precision. Some long-read methods, such as MetaMaps and MMseqs2, required moderate filtering to reduce false positives to resemble the precision and recall of the top-performing methods. We found read quality affected performance for methods relying on protein prediction or exact k-mer matching, and these methods performed better with PacBio HiFi datasets. We also found that long-read datasets with a large proportion of shorter reads (<2kb length) resulted in lower precision and worse abundance estimates, relative to length-filtered datasets. Finally, for classification methods, we found that the long-read datasets produced significantly better results than short-read datasets, demonstrating clear advantages for long-read metagenomic sequencing.

**Conclusions:** Our critical assessment of available methods provides best-practice recommendations for current research using long reads and establishes a baseline for future benchmarking studies.

## Introduction

The identification of microbial species in environmental communities is an essential task in microbiology. Shotgun metagenomic sequencing (or metagenomics) can provide relatively unbiased sampling of the species in such communities, which can include bacteria, archaea, viruses, and eukaryotes. Whereas selective amplification (e.g., 16S, ITS) targets specific gene regions, the goal of metagenomics is to sequence complete genomic DNA for all species in a sample. Consequently, the set of tools used to predict the identities and relative abundances of microbial species differs greatly between these approaches. In particular, the difficulty of performing this task for complex shotgun sequencing data has led to the development of many taxonomic profiling methods, particularly for second-generation/short-read technologies (reviewed in [1]). The rapid expansion of short-read taxonomic classification and profiling tools led to recognition of the importance of methods comparisons, benchmarking, and standardized test datasets [1–10]. These benchmarking studies have been critical for understanding the relative performance of taxonomic profiling methods for different use-cases, which can vary greatly among microbiologists.

Though much of metagenomics has focused on short-read sequencing, there is rising awareness of the new opportunities offered by third-generation sequencing technologies which produce longer sequencing reads. Whereas short reads typically contain a single gene fragment, long reads often span multiple genes and intergenic regions which can be used for alignment algorithms and sequence matching. Among the most popular long-read sequencing platforms are those produced by Pacific Biosciences (PacBio) and Oxford Nanopore Technologies (ONT). While long reads have historically been accompanied by higher error rates, continual improvements in library preparation, sequencing chemistries and post-processing have dramatically reduced the error rates associated with longer reads. For example, the most recent combination of ONT “Q20” chemistry and the Bonito basecaller (v0.3.5+) is reported to produce modal read accuracies of 99% (∼Q20), and the development of PacBio HiFi sequencing allows for highly accurate consensus reads (>Q20, median Q30) that are 10–20 kb in length [11]. As a result of these improvements, both PacBio HiFi and ONT long reads offer new potential for metagenomic analyses, including metagenome assembly, functional annotation, and taxonomic profiling.

Until recently, few studies have evaluated the performance of taxonomic classification and profiling methods for long reads, in part because few tailored methods were available. However, the rate of development for long-read taxonomic classification methods appears to be increasing. For example, MetaMaps [12] and MEGAN-LR [13] were among the first long-read methods, and they became available over the course of several years. By contrast, multiple methods have appeared in the beginning of 2021, including MMseqs2 taxonomy [14] and BugSeq [15]. Prior long-read benchmarking studies applied short-read methods to long reads [3, 16] or compared the potential of long reads to short reads [17], yet only one study has included a comparison of long-read methods [18]. Given the dramatic decreases in long read error rates and the proliferation of long read classification methods, there is a pressing need to assess the performance of taxonomic profiling using long reads.

Here, we perform a critical benchmarking study to evaluate the performance of taxonomic classification and profiling methods for long-read datasets. We evaluate 11 methods, including five methods designed for long reads. We include both taxonomic classifiers and taxonomic profilers in our study. Taxonomic sequence classifiers are used to classify all input reads by aligning or matching the information content in reads to databases consisting of comprehensive nucleotide, protein, or whole genome datasets. The resulting matches or alignments are interpreted to provide taxonomic annotations per reads. When aggregated, the per-read classifications can be used to produce a taxonomic profile with relative abundance estimates (often based on read counts). We note that classifiers can also be used with contigs (versus reads), and this approach is generally referred to as taxonomic binning. However, taxonomic binning precludes relative abundance estimation unless additional steps are included. By contrast, taxonomic profilers are not intended to classify all input reads. Rather, they are designed to output a taxonomic profile with relative abundance estimates. Several profilers rely on smaller marker-specific databases, with contents selected to represent the unique signatures of species. For these marker-based profiling methods, it is expected that only a subset of reads will map successfully. However, profiling methods are not inherently restricted to marker-specific databases, and some methods can use comprehensive databases (see Materials and Methods). We also note that some methods may not be easily categorized as a classifier or profiler. Finally, we distinguish long-read methods from short-read methods as those which utilize the long-range information contained across a long read (often using multiple genes for classification).

We propose the ideal taxonomic classifier and profiler should display high precision and recall (e.g., low numbers of false positives and false negatives), and accurately estimate the relative abundances of taxa [1, 3–4, 7–10]. Furthermore, taxonomic classifiers should ideally assign all assignable reads (e.g., those with database representation). Given the design of marker-based profiling methods, read assignment is not as relevant as a metric of performance. We evaluate the relative performance of methods based on these criteria, using publicly available datasets. These datasets are generated from mock communities of known compositions, which were sequenced using PacBio HiFi or ONT. Mock communities are considered simplistic relative to environmental samples, but they allow a clear assessment of detection metrics (such as precision, recall, and F-scores) and are therefore highly informative for benchmarking. In order to tease apart the impacts of error profile and read length on performance, we also include comparisons using Illumina short-read datasets for two of the mock communities. Our main goals are to 1) identify which methods perform best for long-read datasets, 2) understand if long reads provide more accurate taxonomic profiles or abundance estimates relative to short reads, and 3) identify if differences in long read quality have any effects on performance. Overall, we provide a baseline assessment of available methods using reproducible analyses, which can inform current research and establish a foundation for future benchmarking studies.

## Materials and Methods

### Mock Community Datasets

We obtained two PacBio HiFi datasets and two ONT datasets from publicly available sources. We chose empirical datasets versus simulated datasets because simulations do not capture true variation in error profiles, read length heterogeneity, and the effects of DNA extraction, library preparation, and sequencing. Furthermore, pseudo-mock communities (e.g., those created from multiple isolate sequencing datasets) may combine older and newer sequencing chemistries/platforms for a given technology, creating additional confounding effects.

The two PacBio datasets are available on NCBI (Table 1). The first PacBio HiFi dataset is for the ATCC MSA-1003 mock community (PRJNA546278: SRX6095783, released June 2019). The ATCC MSA-1003 mock community contains 20 bacteria species in staggered abundances (5 species at 18%, 1.8%, 0.18% and 0.02% abundance levels, respectively). The PacBio ATCC dataset was generated using the Sequel II System and contains 2.4 million HiFi reads with a median length of 8.3 kb, for a total of 20.54 Gb of data (Fig. 1, Table 1). We refer to this dataset as HiFi ATCC MSA-1003. The second PacBio HiFi dataset is for the ZymoBIOMICS Gut Microbiome Standard D6331 (PRJNA680590: SRX9569057, released November 2020). The Zymo D6331 mock community contains 17 species (including 14 bacteria, 1 archaea, and 2 yeasts) in staggered abundances. Five species occur at 14% abundance, four at 6%, four at 1.5%, and one species per 0.1%, 0.01%, 0.001%, and 0.0001% abundance level. There are five strains of *E. coli* contained in this community (each at 2.8% abundance), which we treat here as one species at 14% abundance. The PacBio Zymo D6331 dataset was generated using the Sequel II System and contains 1.9 million HiFi reads with a median length of 8.1 kb, for a total of 17.99 Gb of data (Fig. 1, Table 1). We refer to this dataset as HiFi Zymo D6331.

**Figure 1.**
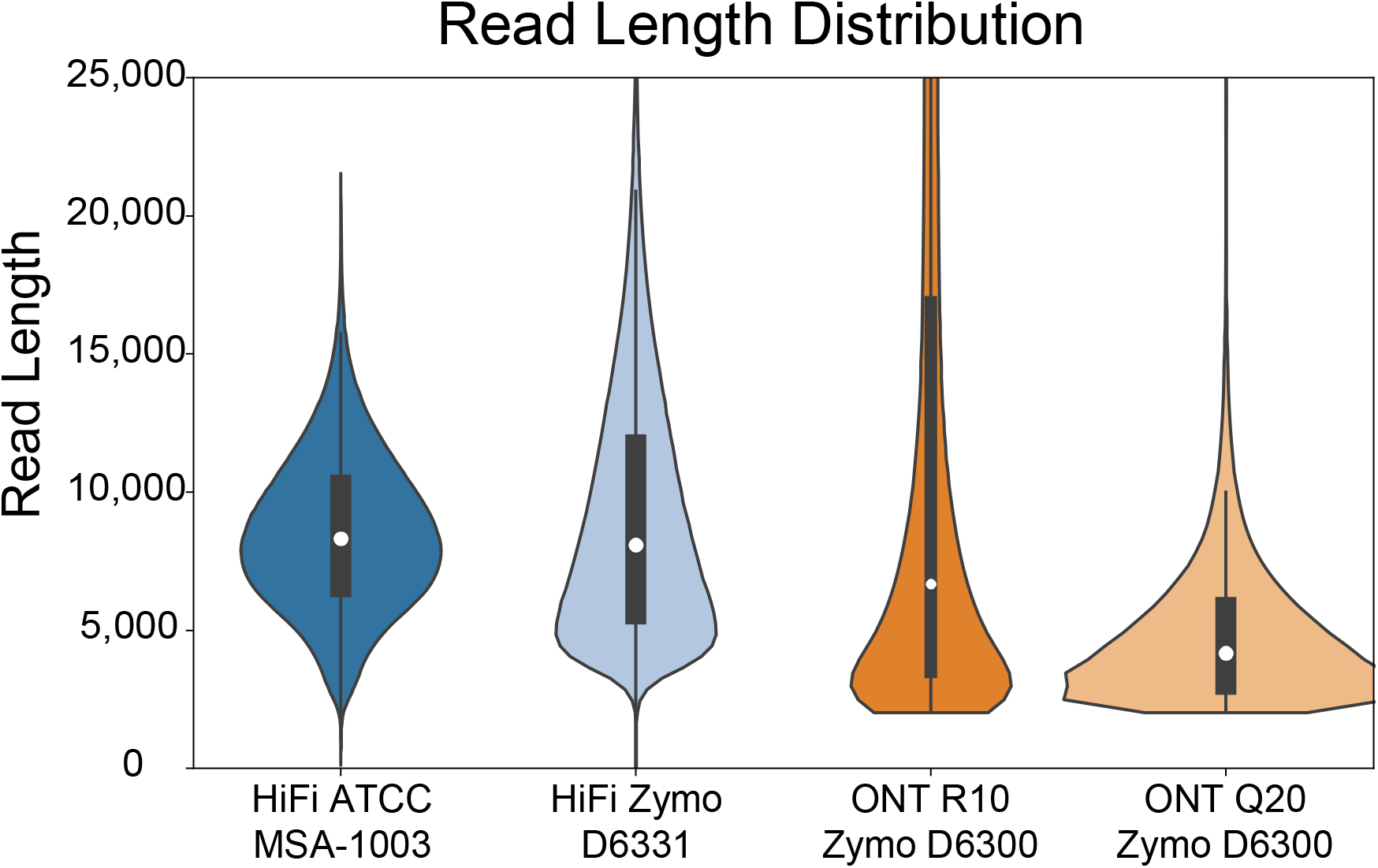
Violin plots showing the read length distributions for the four mock community datasets included in this study, after length-filtering was applied to remove shorter reads (see methods). Interiors of plots contain white dots representing median values, black bars represent interquartile values, and black lines represent minimum and maximum range values. Read sizes range up to 50,000 bp in length, but the plot is clipped at 25,000 bp to show the core size distributions.

**Table 1.**
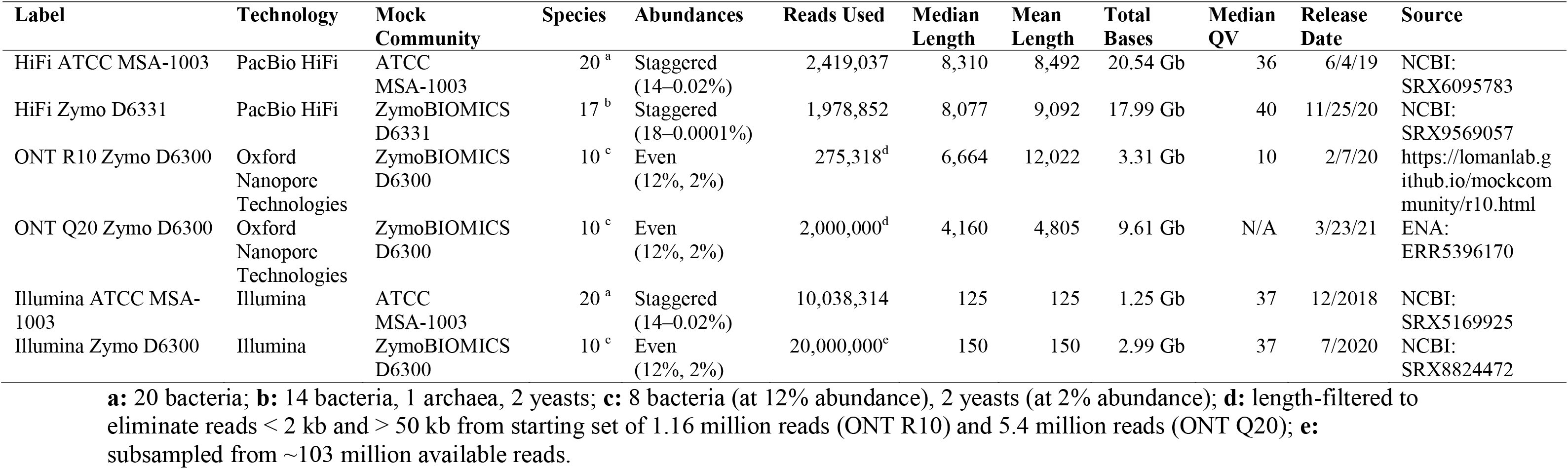
Description of the publicly available mock community datasets used for this experiment.

We obtained two ONT datasets for the ZymoBIOMICS D6300 microbial community standard. The Zymo D6300 standard is simpler in design and contains 10 species in even abundances, including 8 bacteria at 12% abundance and two yeasts at 2% abundance. The two ONT datasets contained a broader distribution of read lengths which included a large tail of shorter reads (<2kb in length). Our initial work indicated these shorter reads may have an adverse effect on taxonomic profiling, a result also supported by [19]. We therefore included two variations of each ONT dataset. The primary datasets are the focus of our methods comparison and resulted from length filtering to remove all short reads (<2kb) and ultra-long reads (>50kb). We found ultra-long reads caused compatibility issues with some taxonomic profiling programs (particularly the short-read methods). To investigate the potential effects of shorter reads, we created secondary datasets which contained a large proportion of shorter long reads. The first ONT dataset comes from a continually updated resource produced by [20]. We downloaded the R10.3 chemistry data release (February 2020) which was produced from two flowcells on an ONT GridION, resulting in 1.16 million reads (4.64 Gb data). We used NanoFilt [21] to remove all short (<2kb) and ultra-long reads (>50kb). Length-filtering resulted in the removal of 873,079 short reads and 12,129 ultra-long reads (1.33 Gb total; 75% and 0.01% of total reads, respectively), and the retention of 275,318 ONT reads (23% of total reads). The resulting length filtered ONT reads have a median length of 6.6 kb, for a total of 3.31 Gb of data (Fig. 1, Table 1). We refer to this primary dataset as ONT R10 Zymo D6300. The secondary version of this dataset uses all reads <50kb in length. It contains 3.86 Gb data (1,148,397 reads) with a median read length of 660 bp and mean read length of 3.3 kb, and is referred to as ONT R10 Short (Supplementary Figure S1). The second ONT dataset was obtained from the European Nucleotide Archive (PRJEB43406: ERR5396170, released March 2021) and represents the ‘Q20 chemistry’ release for the Zymo D6300 standard (described at: https://github.com/Kirk3gaard/2020-05-20_ZymoMock_Q20EA). It was generated using a PromethION, resulting in 5.4 million reads (17.95 Gb data). We again used NanoFilt to remove short reads (<2 kb) and ultra-long reads (>50 kb), which resulted in the elimination of 2.13 million (39%) and 819 (<0.001%) of the total reads, respectively. From the remaining ∼3.2 million reads, we subsampled to obtain 2 million reads (a number comparable to the HiFi datasets). This produced a length filtered ONT dataset of 2 million reads with a median length of 4.2 kb, for a total of 9.6 Gb of data (Fig. 1, Table 1). We refer to this primary dataset as ONT Q20 Zymo D6300. The secondary version of this dataset contains a comparable number of shorter long reads. We used NanoFilt to remove all reads >3kb in length and subsampled the remaining reads to obtain 2 million reads. We refer to this as ONT Q20 Short, and this dataset contains 2.72 Gb data with a median read length of 1.2 kb and mean read length of 1.3 kb (Supplementary Figure S1). The read names required to reconstitute the ONT R10 Zymo D6300 and ONT Q20 Zymo D6300 datasets are available on the Open Science Framework project page for this paper (https://osf.io/bqtdu/).

As a final comparison to the long-read datasets, we included short-read sequence data for two of the mock communities (Table 1). We downloaded Illumina sequence data for ATCC MSA-1003 (PRJNA510527: SRX5169925, released December 2018), which included a total of ∼10 million 150 bp paired-end reads produced by a HiSeq2500 (but available pre-trimmed to 125 bp). We also obtained Illumina sequence data for the Zymo D6300 community (PRJNA648136: SRX8824472, released July 2020). These data were produced using a NovaSeq 6000 and include ∼100 million 150bp PE reads. Given the large difference in read numbers between these datasets, we subsampled the Zymo Illumina data to obtain 20 million total reads. We refer to these datasets as Illumina ATCC MSA-1003 and Illumina Zymo D6300, respectively. A variety of factors, including different DNA extraction methods, can affect the final composition of DNA sequenced for metagenomic samples and potentially bias relative abundance estimates [21]. Additionally, variation in error profiles across sequencing technologies could also cause potential differences in results. To control for these potential confounding effects in the Illumina datasets, we also “simulated” short-read data from our long-read datasets. Each long read was divided into 150 bp non-overlapping segments, and 10 segments were randomly selected to create a “simulated” short-read dataset. We chose this subsampling strategy (versus retaining all available segments) to create a consistent number of short reads per long read, which varied in length. This strategy generated ∼21 million 150 bp “reads” from the HiFi ATCC MSA-1003 dataset, and 20 million 150 bp “reads” from the ONT Q20 Zymo D6300 dataset. We refer to these datasets as SR-Sim ATCC MSA-1003 and SR-Sim ZymoD6300, respectively.

### Taxonomic Classification and Profiling Methods

We evaluated the performance of 11 methods on the long-read mock community datasets. We included five methods developed specifically for long reads, five popular short-read methods, and one generalized method (Table 2), which we summarize here. We ran all methods for the primary long-read datasets and secondary ONT datasets, and used only short-read methods for the short-read datasets.

**Table 2.** Overview of taxonomic profiling methods used in this experiment.

| **Intended Use** | **Method** | **Variations** | **Reference Database** | **Strategy** |
| --- | --- | --- | --- | --- |
| Short reads | Kraken2 | - | “PlusPF” | K-mer-based |
|  | Bracken | - | “PlusPF” | Bayesian Refinement |
|  | Centrifuge | h22, h500 | Refseq ABVF | BW transform, FM-index |
|  | MetaPhlAn3 | - | mpa_v30_CHOCOPhlAn_201901 | Read mapping, coverage scores |
|  | mOTUs2 | - | V3.0.3 | Read mapping |
| General | sourmash | k31, k51 | GenBank | K-mer min-set-cov; LCA algorithm |
| Long reads | MetaMaps | - | MiniSeq+H | Approximate mapping |
|  | MMseqs2 | - | NCBI nr | Translation alignment, LCA algorithm |
|  | MEGAN-LR-prot | - | NCBI nr | Translation alignment, LCA algorithm |
|  | MEGAN-LR-nuc | HiFi, ONT | NCBI nt | Nucleotide alignment, LCA algorithm |
|  | BugSeq-V2 | - | NCBI nt | Nucleotide alignment, LCA algorithm |

The short-read methods include Kraken2 [23–24], Bracken [25], Centrifuge [26], MetaPhlAn3 [27], and mOTUs2 [28]. Among these methods, Kraken2 and Centrifuge are taxonomic sequence classifiers, Bracken is a type of taxonomic profiler, and MetaPhlAn3 and mOTUs2 are both marker-based taxonomic profilers. Kraken2 is a k-mer-based read classifier, which is often paired with Bracken for profiling. Following Kraken2 analyses, Bracken is used for Bayesian re-estimation of abundances. Centrifuge uses a Burrows-Wheeler transform and Ferragina-Manzini index for storing and mapping sequences. We include two variations of Centrifuge analyses, one using the default settings suitable for short reads (referred to as Centrifuge-h22), and another with settings for long reads (referred to as Centrifuge-h500; see details below). MetaPhlAn3 uses coverage scores to calculate the relative abundances of taxa, based on read mapping to a unique clade-specific marker database. Similarly, mOTUs2 maps reads to a unique marker specific database. Specifically, it uses a database composed of single copy phylogenetic marker genes for operational taxonomic units (mOTUs). Recently, a “long read” option was introduced for mOTUs2, which divides each long read into multiple short read segments (highly similar to our SR-Sim datasets) and uses these outputs to run the typical short read workflow. We used the “long read” option for our analyses as recommended by the authors, but note that it should not be considered a true long-read method. The resulting artificial short read datasets contained 25–35x more reads than the initial long read datasets.

The long-read methods include MetaMaps [12], MEGAN-LR [13, 29], MMseqs2 [14], and BugSeq [15]. All long-read methods described here are considered taxonomic sequence classifiers. MetaMaps was among the first methods designed specifically for long reads, and it uses approximate mapping with probabilistic scoring to estimate sample composition. MEGAN­LR was developed from MEGAN6 and was designed to interpret translation alignments of long nucleotide sequences to a protein reference database. These alignments can be made using any program capable of translation alignment (e.g., blastx mode), but here we specifically use DIAMOND [30] due to its favorable long-read options (e.g., range-culling and frameshift-aware alignment; [31]). MEGAN-LR assigns reads to taxa using a novel interval-union lowest common ancestor (LCA) algorithm, in combination with other relevant features (e.g., lcaCoveragePercent, minSupportPercent, minPercentReadCover). MEGAN-LR can likewise interpret alignments to nucleotide databases using similar options, such as those created with minimap2 [32]. For this experiment, we created alignments based on protein references (using DIAMOND) and nucleotide references (using minimap2), and subsequently used MEGAN-LR for taxonomic classification. To distinguish between these methods, we refer to them as MEGAN-LR-prot and MEGAN-LR-nuc. Furthermore, we tested settings in minimap2 that were specific to HiFi or ONT data (see below) and ran both settings on all mock communities. We refer to these analyses as MEGAN-LR-nuc-HiFi and MEGAN-LR-nuc-ONT. Thus, we include three analyses that involve MEGAN-LR: MEGAN-LR-prot, MEGAN-LR-nuc-HiFi, and MEGAN-LR-nuc-ONT. We note that MEGAN-LR-prot is unique from all other methods in that it also simultaneously assigns functional annotations to genes on reads, providing a taxonomic and functional profile for a sample. The MMseqs2 taxonomy tool extracts all possible protein fragments in six frames from the long reads, pre-filters the protein sequences, aligns the retained protein sequences to the reference protein database, and ultimately assigns reads to taxa using a novel LCA algorithm (“approximate 2bLCA”). The published BugSeq algorithm (V1) performs minimap2 alignments using a nucleotide database, followed by Bayesian reassignment and LCA identification [15]. Following initial development, a BugSeq V2 method was developed which includes minimap2 alignment of sequences to a nucleotide database followed by LCA identification and abundance calculation (S. Chorlton, personal communication). BugSeq V2 performs better for longer reads (>1kb), higher sequencing depth, and shotgun metagenomics (vs. cDNA sequencing experiments). An auto-detect feature selects the V1 or V2 version based on the dataset uploaded to the online platform, and in our experiment BugSeq V2 was selected for all long-read datasets.

In addition to methods which are generalized to short or long reads, we also ran sourmash [33, 34], which is a k-mer-based sequence analysis tool that can be used for taxonomic profiling. Sourmash uses a fractional scaling (‘FracMinHash’) approach to representatively subsample both metagenome and reference datasets in a way that supports accurate sequence similarity comparisons [35]; this allows rapid search of large databases. Sourmash can be used with any type of sequencing data, but its taxonomic profiling (sourmash gather + sourmash taxonomy) has thus far been primarily applied to short reads datasets. Sourmash profiling differs from the k-mer methods above in that it uses combinatorial observations of k-mers to find the minimum set of reference genomes that cover all information (k-mers) in the metagenome query, and then aggregates the taxonomic information from these genomes using an LCA approach [35]. Long nucleotide k-mer exact matching is more stringent than alignment-approaches, with stringency increasing as k-mer length increases. As a result, long k-mer searches may miss some reference matches if sufficient nucleotide divergence exists between the metagenome sequence and the strain available in the reference database [36]. Sourmash uses a k-mer length of 31 for species-level matching (default), and suggests 51 for strain-level resolution; we test both here. We use the default fractional scaling (1/1000) for all analyses.

A standardized output format was required to facilitate comparisons of the results across methods. We selected kraken-report (kreport) format because it contains cumulative counts and level counts across the complete hierarchical taxonomy for each taxon assigned. The level count is the number of reads specifically assigned to a taxon, whereas the cumulative count is the sum of the level counts for a taxon plus its descendants. For example, the cumulative count of a genus is the level count for that genus plus the level counts of all species and strains contained in that genus. This output format is readily available for Kraken2, Bracken, MMseqs2, and BugSeq. We created conversion tools for all other methods (MetaPhlAn3, MetaMaps, MEGAN-LR), which are available on github: https://github.com/PacificBiosciences/pb-metagenomics-tools. The kreport output format was recently added to sourmash and is available in sourmash v4.5.1.

### Comparative Analyses

We evaluated method performance using several criteria. We assessed read utilization, detection metrics at the species and genus level, and relative abundance estimates. We provide details for each of these categories below.

#### Read Utilization

We evaluated read utilization for each profiling method in two ways. First, we simply calculated the total percent of reads that received a taxonomic assignment. For sourmash, we use the total percent of the dataset with an assignment, as it does not assign taxonomy to specific reads. Second, we calculated the percentage of reads (dataset) that were assigned to specific taxonomic levels. We performed this for the following ranks: class, order, family, genus, species, and subspecies/strain. Values were obtained by summing the level counts of all taxa within a given rank. In general, we expected methods that utilize LCA algorithms to display read assignments across multiple taxonomic levels, relative to methods that do not. The exception is sourmash, which makes non-overlapping k-mer assignments to specific genomes (∼strain level) and only uses LCA to aggregate genome matches to higher taxonomic ranks. We expected marker-based profilers (MetaPhlAn3, mOTUs2) to display relatively low read assignments, and mainly used read utilization to evaluate performance among the remaining methods.

#### Detection Metrics

The species compositions of the mock communities are known, allowing for a complete evaluation of detection metrics. For each profiling method, we scored the presence/absence of a taxon based on whether or not the cumulative read count for that taxon exceeded a minimum percent threshold of the total reads. We used a minimum percent threshold (versus a fixed number of reads) because our datasets contained different numbers of total reads. We recognize that setting a minimum detection threshold in this way penalizes methods that assign a smaller proportion of the total reads available. However, setting a threshold based on the number of reads assigned in a given analysis could produce misleading results (for example, a method could assign only 10% of total reads but achieve perfect precision). We evaluated three minimum read thresholds, including 0.001% (mild filtering, mainly for removing singleton count taxa for short-read methods), 0.1% (moderate filtering), and 1% (heavy filtering) of the total number of reads per dataset (Table 3). The threshold filtering was mainly used to explore the effects on precision (particularly the impact on false positives) across the four primary datasets. However, we also used filtering to investigate the effects on the staggered abundance communities (ATCC MSA-1003 and Zymo D6331). These two mock communities contained several taxa in low abundances, and we explored how filtering might cause detection dropout for different abundance levels. We performed our evaluations at the species level and the genus level. We expected detection to be more difficult at the species level and easier at the genus level. This is because assignments to multiple non-target species within a genus would be considered incorrect at the species level, but correct at the genus level.

**Table 3.** Minimum detection thresholds used to score the presence/absence of mock community taxa at the species or genus level.

| **Dataset** | **Number of Reads** | **0.001% Threshold** | **0.1% Threshold** | **1% Threshold** |
| --- | --- | --- | --- | --- |
| HiFi ATCC MSA-1003 | 2,419,037 | 24 | 2,419 | 24,190 |
| HiFi Zymo D6331 | 1,978,852 | 19 | 1,978 | 19,788 |
| ONT R10 Zymo D6300 | 275,318 | 2 | 275 | 2,753 |
| ONT Q20 Zymo D6300 | 2,000,000 | 20 | 2,000 | 20,000 |
| Illumina ATCC MSA1003 | 10,038,314 | 100 | 10,038 | 100,383 |
| Illumina Zymo D6300 | 20,000,000 | 200 | 20,000 | 200,000 |

We calculated several detection metrics (precision, recall, F-scores) which are based on the number of true positives, false positives, and false negatives. In this context, we define a true positive as the detection of a mock community taxon (based on a read count exceeding the minimum read threshold). We define a false positive as the detection of taxon that is not present in the mock community. We define a false negative as the failure to detect a taxon in the mock community (based on a zero count or count below the minimum threshold). The formulas for precision, recall and F-scores are as follows:

> **Precision** = true positives / (true positives + false positives)
>
> **Recall** = true positives / (true positives + false negatives)
>
> **F1** = (2 * precision * recall) / (precision + recall)
>
> **F0.5** = ((1 + 0.5^2^) * precision * recall) / ((0.5^2^ * precision) + recall)

The values for the above metrics each range from 0 to 1. For precision, a score of 1 indicates only mock community taxa were detected, whereas lower scores indicate detection of additional taxa (e.g., false positives). For recall, a score of 1 indicates all taxa in the mock community were detected, whereas a lower score indicates some taxa were not detected. The F-scores provide a useful way to summarize the information from precision and recall. The F1 score is the harmonic mean of precision and recall (both measures are weighted equally), whereas the F0.5 score gives more weight to precision (placing more importance on minimizing false positives). A value of 1 for either F-score indicates perfect precision and recall.

We controlled for two issues that can negatively impact these metrics. First, we observed and accounted for differences in taxonomy, particularly as it relates to synonymies. In the case of a species synonomy, we used the sum of cumulative counts for the species and all synonyms as the read count for the taxon. This included two species in ATCC MSA-1003 (*Luteovulum sphaeroides* = *Rhodobacter sphaeroides*, *Cereibacter sphaeroides; Phocaeicola vulgatus* = *Bacteroides vulgatus*), one species in Zymo D6300 (*Limosilactobacillus fermentum* = *Lactobacillus fermentum*), and three species in Zymo D6331 (*Limosilactobacillus fermentum* = *Lactobacillus fermentum; Bacillus subtilis* = *Bacillus spizizenii; Faecalibacterium sp. AF28­13AC* = *Faecalibacterium prausnitzii*). Most of these synonomies are related to changes in taxonomy, but for *Faecalibacterium prausnitzii* we observed that *Faecalibacterium sp. AF28­13AC* contained a genome sequence identical to *F. prausnitzii* in the NCBI database. Second, we observed that sequences and/or taxonomy information was lacking for two species (Zymo D6331: *Veillonella rogosae, Prevotella corporis*) in multiple databases (“PlusPF”, Refseq ABVF, MiniSeq+H, NCBI nt). To remedy this issue, we excluded the two species from the set of taxa used to calculate detection metrics at the species-level for all methods. However, we observed that many reads were assigned to alternate species in the same genus, so we included the two genera in the genus-level analysis.

We calculated detection metrics for each dataset. To understand the performance of each method across all datasets, we took an average of precision, recall, F_1_ and F_0.5_. We also took the average of these values for the HiFi datasets and ONT datasets separately, to see if any methods performed differently across the technologies.

#### Relative abundance estimates

We attempted to obtain relative abundances for each method, but acknowledge several potential issues. First, there are clear differences in intended outputs among methods. For example, profiling methods provide taxonomic abundances whereas classifiers provide sequence abundances (which must be transformed into taxonomic abundances). Second, the read counts obtained from classifiers do not account for the length heterogeneity of reads in long read datasets, and counts are not weighted by total base pairs. Although some methods offer this type of correction (MEGAN-LR), it is not available across all methods and difficult to implement. Third, DNA extraction methods can affect the final composition of DNA sequenced for metagenomic samples [21], which could lead to systematically skewed abundance estimates. Despite these caveats, relative abundances are of interest to the research community and are therefore included here.

We used the read counts output directly from Kraken, Bracken, Centrifuge, mOTUs2, MetaMaps, MMSeqs2, all MEGAN-LR methods, and BugSeq. The output of sourmash is abundance-projected base pair estimates, which is a projection of the number of base pairs that the percent of matched k-mers represents. To estimate the “read counts” for this method, we obtained a total from the base pair estimates across species plus all unassigned base pairs, and divided the base pair estimates from all species by this total. For MetaPhlAn3, we multiplied the percent abundance of each taxon by the total number of mapped reads. We note that for mOTUs2, the read counts are based on the artificial short reads generated, and not the initial long reads. These numbers therefore represent an overestimate. However, given the low read counts recovered using this method (<1%; see Results), we did not attempt to transform these read counts.

Relative abundances were estimated for each profiling method at the species and genus level. We obtained cumulative counts for the mock community species or genera and the sum of cumulative counts for all false positives at the species or genus level (classified as “Other”). These data were normalized to obtain the percent abundance of each taxon. We corrected for the absence of two species from multiple databases (*Veillonella rogosae, Prevotella corporis*) in HiFi Zymo D6331. For methods affected by these databases, we observed many reads were assigned to other species in these two genera. Rather than scoring these as “Other”, we allowed all species-level assignments within these genera to contribute to the read counts for these two species. To be consistent, we allowed this for all methods for HiFi Zymo D6331. In other words, genus-level counts for *Veillonella rogosae* and *Prevotella corporis* were used for the species abundances, rather than exclude these two taxa.

For each method, we calculated an L1 distance (following [9]) and performed a chi-squared goodness of fit test to determine if the estimated abundances were significantly different from the theoretical abundances. The theoretical abundances were obtained from the manufacturer’s specifications, which are based on genomic DNA (versus cell counts). We calculated L1 distance by summing the absolute error between the theoretical and empirical estimate per species per community. We included the false positives lumped in the “Other” category in this calculation and compared them against a theoretical abundance of zero for this category. We compared the chi-squared statistic to the critical value obtained at the 95% significance level and obtained a corresponding P-value. For this test, larger chi-squared statistic values indicate greater differences between the observed and expected values. We applied a Bonferroni correction for multiple testing (n = 11) per dataset, for which α_altered_ = 0.05/11 = 0.0045. A P-value < 0.0045 allows rejection of the null hypothesis, and indicates the observed distribution is significantly different from the theoretical distribution.

### Reference Databases

The choice of reference database directly affects the outcome of taxonomic profiling. For example, the use of a complete reference database versus a subset of that database can result in drastically different assignments if the same profiling method is run with otherwise identical settings. Under ideal conditions, all profiling methods would use an identical reference database. This would control for differences in information content and taxonomy, allowing observed differences in assignment results to be attributed to the profiling methods. However, differences in method design and matching algorithms required the use of multiple reference databases. We therefore provide a brief description and comparison of these databases below.

The databases used for Kraken2, Bracken, and Centrifuge are highly similar. For Kraken2 and Bracken, we used a pre-built database that includes all RefSeq sequences for archaea, bacteria, viruses, plasmid, human, protozoa, and fungi (“PlusPF”, released 1/27/2021, available from: https://benlangmead.github.io/aws-indexes/k2). The Centrifuge database was built from RefSeq sequences for archaea, bacteria, viruses, and fungi (downloaded 4/2021). The Centrifuge database used can be considered a subset of the PlusPF database, but with complete overlap for several target groups (archaea, bacteria, fungi).

The marker-based profilers each used a specific database. MetaPhlAn3 uses a highly distinct reference database which is composed of ∼1.1 million unique clade-specific markers from ∼99,500 bacteria/archaea reference genomes and ∼500 eukaryotic reference genomes. We used the mpa_v30_CHOCOPhlAn_201901 database release. mOTUs2 also uses a highly distinct database, which is composed of single copy phylogenetic marker genes for operational taxonomic units (mOTUs). We used database version 3.0.3, which contains ∼12,000 reference based mOTUs, ∼2,300 mOTUs obtained from metagenomic samples, and ∼19,400 MAG-based mOTUs.

MetaMaps provides a pre-built database composed of 12,058 complete RefSeq genomes (215 archaeal, 5774 bacterial, 6059 viral/viroidal, 7 fungi, 1 human), which is referred to as MiniSeq+H. The option to create a custom database (such as NCBI nt) was initially developed for MetaMaps, but this feature is currently not functional. The MiniSeq+H database was therefore the only option available for running MetaMaps in our experiment, and it represents the smallest and most incomplete database across the methods used.

We used the NCBI non-redundant protein database (NCBI nr) for MMseqs2 and MEGAN-LR-prot, and the NCBI nucleotide database (NCBI nt) for MEGAN-LR-nuc and BugSeq v2 (both databases downloaded April 2021). We used a more recent version of the NCBI nucleotide database for sourmash (downloaded March 2022), which was added in our revision to this manuscript. These pre-built sourmash databases consist of 47952 viral, 8750 archaeal, 1193 protozoa, 10286 fungi, and 1148011 bacterial GenBank genomes and were constructed using FracMinHash 1/1000 fractional scaling (∼1.3million genomes, ∼40G size all together; available at https://sourmash.readthedocs.io/en/latest/databases.html). Sourmash provides a corresponding lineages file with taxonomic information for each database. The NCBI nt databases represent the most complete reference databases across the methods. We note that the RefSeq databases for Kraken2, Bracken, and Centrifuge are contained in NCBI nt.

### Profiling Method Commands

To facilitate reproducible results, we provide the general commands or instructions to run each method.

#### Kraken2

We ran Kraken version 2.1.1 for each sample. We used the pre-built PlusPF database described above, and used the following command:

~~~
kraken2 --db PlusPF --threads 24 –report SAMPLE.kreport.txt
SAMPLE.fasta > SAMPLE.kraken
~~~

#### Bracken

We ran Bracken version 2.6.0 for each sample, using the kreport outputs from Kraken2. We used the pre-built PlusPF database described above, and the following command to obtain abundances at the species level (-l S):

~~~
bracken -d PlusPF -i SAMPLE.kreport.txt -o SAMPLE.bracken -r 50
-l S -t 10
~~~

#### Centrifuge

We ran Centrifuge version 1.0.4. We were unable to use centrifuge-download to obtain the RefSeq sequences required to build the database. We instead used kraken2-build to obtain the relevant RefSeq sequences and taxonomy files. The kraken headers were removed from the fasta sequences, and the database was built using the following command:

~~~
centrifuge-build -p 24 --conversion-table centrifuge-
seqid2taxid.map --taxonomy-tree /taxonomy/nodes.dmp --name-table
/taxonomy/names.dmp arc-bac-vir-fungi.fna abvf
~~~

Centrifuge offers the option to specify the minimum length of partial hits required for classification (--min-hitlen). We used two values for this option. We used the default value of 22, which is suitable for short read analysis, and used a value of 500 which is suitable for long reads (labeled as Centrifuge-h22 and Centrifuge-h500, respectively).

We ran Centrifuge-h22 for each sample using the following command:

~~~
centrifuge -f --min-hitlen 22 -k 20 -t -p 24 -x abvf -U
SAMPLE.fasta -S SAMPLE-h22.txt --report-file SAMPLE-
h22.centrifuge_report.tsv
~~~

We ran Centrifuge-h500 for each sample using the following command:

~~~
centrifuge -f --min-hitlen 500 -k 20 -t -p 24 -x abvf -U
SAMPLE.fasta -S SAMPLE-h500.txt --report-file SAMPLE-
h500.centrifuge_report.tsv
~~~

Outputs were converted to kreport format using the centrifuge-kreport module.

#### MetaPhlAn3

Analyses were run using MetaPhlAn v3.0.7. The settings used in MetaPhlAn3 to run Bowtie2 will fail for long reads, so we first created alignments externally using Bowtie2:

~~~
bowtie2 -p 12 -f --local --no-head --no-sq --no-unal -S
SAMPLE.sam -x /metaphlan/mpa_v30_CHOCOPhlAn_201901 -U
SAMPLE.fasta
~~~

After alignments were created, we ran MetaPhlAn3 with the following settings (adjusting the number of reads per dataset, --neads):

~~~
metaphlan SAMPLE.sam --nproc 24 --input_type sam --nreads
READ_NUMBER -o SAMPLE.profiled_metagenome.txt --index
mpa_v30_CHOCOPhlAn_201901 --bowtie2db /metaphlan
~~~

#### mOTUs2

Analyses were run using mOTUs2 v3.0.3. Each long-read dataset was converted into a short read dataset and then run through the typical profiling algorithm using the following set of commands:

~~~
motus prep_long -i SAMPLE.fastq.gz -o SAMPLE_mOTUs.fastq -no_gz
gzip SAMPLE_mOTUs.fastq
motus profile -s SAMPLE_mOTUs.fastq.gz -o
SAMPLE_mOTUs.counts.txt -c -t 48
~~~

#### Sourmash

Analyses were run using sourmash version 4.5.1. A streamlined workflow for sourmash is available (Taxonomic-Profiling-Sourmash) at: https://github.com/PacificBiosciences/pb-metagenomics-tools. The pipeline is provided as a configurable snakemake workflow.

Read datasets were sketched in the same manner as sourmash pre-prepared databases, using a fractional scaling of 1/1000:

~~~
sourmash sketch dna SAMPLE.fna.gz -p k=31,k=51,scaled=1000,abund
-name SAMPLE -o SAMPLE.sig.zip
~~~

The database search was performed separately for each k-mer size using sourmash gather. This analysis took 3-7 hours on a single thread, requiring 40-100G of memory (depending on dataset):

~~~
sourmash gather SAMPLE.sig.zip genbank-2022.03-bacteria-k31.zip
genbank-2022.03-archaea-k31.zip genbank-2022.03-viral-k31.zip
genbank-2022.03-protozoa-k31.zip genbank-2022.03-fungi-k31.zip
-k 31 -o SAMPLE.gather.k31.csv
~~~

After searching with sourmash defaults, we also ran gather at its most sensitive, allowing detection of even a single shared hash in the database (by adding --`threshold-bp 0` to the command). For each dataset and ksize, taxonomic aggregation of genome-level matches was performed using the sourmash taxonomy module, with kreport output, e.g. k31:

~~~
sourmash tax metagenome -g SAMPLE.gather.k31.csv -t genbank-
2022.03-*.lineages.csv.gz -o SAMPLE.gather.k31 -F kreport
~~~

Note that sourmash gather outputs initial k-mer assignments to individual genomes, which is ∼strain-level profiling; we did not evaluate these in our results.

#### MetaMaps

We used MetaMaps v0.1 to run analyses with the following set of commands:

~~~
metamaps mapDirectly --all -r /databases/miniSeq-H/DB.fa -q
SAMPLE.fasta --maxmemory 35 -t 24 -o SAMPLE_results
metamaps classify -t 12 --mappings SAMPLE_results --DB
/databases/miniSeq-H
~~~

The conversion from MetaMaps output format to kreport format was performed at the species level, but we note that MetaMaps can produce a large number of strain assignments that are not represented in our results.

#### MMseqs2

We used MMseqs2 v13.45111 to run all analyses. We first built the database for NCBI nr using the following command:

~~~
mmseqs databases NR /mmseqs-database/NR_db /scratch --threads 24
~~~

We then used the easy-taxonomy module to run analyses for each sample, using the following general command:

~~~
mmseqs easy-taxonomy SAMPLE.fasta /mmseqs-database/NR_db SAMPLE
/scratch --threads 48 --split-memory-limit 120G
~~~

#### MEGAN-LR-prot

A streamlined workflow for MEGAN-LR-prot is available (Taxonomic-Profiling-Diamond-Megan) at: https://github.com/PacificBiosciences/pb-metagenomics-tools. The pipeline is provided as a configurable snakemake workflow. To use the workflows, we first downloaded the NCBI nr database and created a DIAMOND index using the following command:

~~~
diamond makedb --in nr.gz --db diamond_nr_db --threads 24
~~~

We downloaded MEGAN6 community edition to obtain the executable tools required for these workflows (sam2rma, rma2info), as well as the required MEGAN protein mapping file (megan­map-Jan2021.db). We then ran the Taxonomic-Functional-Profiling-Protein pipeline. The locations of the nr index, sam2rma, and the mapping file were specified in the main configuration file for the analysis (config.yaml), and we used all other default settings (see documentation). The information for the sample fasta files was added to the sample configuration file (Sample-Config.yaml), and the snakemake (Snakefile-taxprot) was executed. Details for the usage of each program are provided in the online documentation.

Analyses resulted in RMA output files, which were used as inputs for the MEGAN-RMA-Summary pipeline. The location of rma2info was specified in the main configuration file for the analysis (config.yaml), information for the sample fasta files was added to the sample configuration file (Sample-Config-protein.yaml), and we created the required sample-read­counts file. This snakemake (Snakefile-summarizeProteinRMA) was run using all other default settings, and kreport files were included in the outputs.

#### MEGAN-LR-nuc

A streamlined workflow for MEGAN-LR-nuc is available (Taxonomic­Profiling-Minimap-Megan) at: https://github.com/PacificBiosciences/pb-metagenomics-tools. The pipeline is provided as a configurable snakemake workflow. To use the workflow, we first downloaded the NCBI nt database and indexed it with minimap2 using the following command:

~~~
minimap2 -k 19 -w 10 -I 10G -d mm_nt_db.mmi nt.gz
~~~

We downloaded MEGAN6 community edition to obtain the executable tools required for these workflows (sam2rma, rma2info), as well as the required MEGAN nucleotide mapping file (megan-nucl-Jan201.db). We then ran the Taxonomic-Profiling-Nucleotide pipeline. The locations of the minimap2 nt index, sam2rma, and the mapping file were specified in the main configuration file for the analysis (config.yaml), and we also changed the maximum number of secondary alignments from 20 to 5. The information for the sample fasta files was added to the sample configuration file (Sample-Config.yaml), and the snakemake (Snakefile-taxnuc) was executed. Details for the usage of each program are provided in the online documentation.

Analyses resulted in RMA output files, which were used as inputs for the MEGAN-RMA-Summary pipeline. The location of rma2info was specified in the main configuration file for the analysis (config.yaml), information for the sample fasta files was added to the sample configuration file (Sample-Config-nucleotide.yaml), and we created the required sample-read­ counts file. This snakemake (Snakefile-summarizeNucleotideRMA) was run using all other default settings, and kreport files were included in the outputs.

The above instructions are for the MEGAN-LR-nuc-HiFi analysis. Running the MEGAN-LR­nuc-ONT analysis required some changes. Specifically, we indexed the database with minimap2 using the following command:

~~~
minimap2 -k 15 -w 10 -I 10G -d mm_nt_db_ONT.mmi nt.gz
~~~

We then edited the minimap2 command in the snakemake file to include the ONT recommended settings:

~~~
minimap2 -ax map-ont
~~~

#### BugSeq

We uploaded datasets to the BugSeq online platform: https://bugseq.com. For each dataset, we selected the NCBI nt reference database option, and submitted the analysis. After successful completion all results were available for download.

## Results

The kreport files produced from all taxonomic classification and profiling methods, and the Jupyter notebooks used to generate the following results, are freely available on the Open Science Framework project page for this paper (https://osf.io/bqtdu/). These files can be used to replicate all results reported below.

### Comparative Analyses

*Read Utilization.* Total read assignment differed drastically across methods (Fig. 2). In terms of short-read methods, Kraken, Bracken, and Centrifuge-h22 assigned the greatest number of reads (93–100% for HiFi, 81–99% for ONT). Centrifuge-h500, which required a minimum total length of 500 for partial hits, assigned far fewer reads across datasets (1–53%), with the exception of HiFi ATCC MSA-1003 (which had 98% read assignment). Read assignment was exceptionally low for Centrifuge-h500 in ONT R10 Zymo D6300 (∼1%; Fig. 2). As expected, both marker-based profilers assigned the fewest reads (MetaPhlAn3: 23–39%; mOTUs2: 0.2–1%; Fig. 2). Slightly more of the dataset was assigned by sourmash-k51 versus k31 (4–15% difference; Fig. 2). However, the greatest difference in sourmash assignment occurred between HiFi and ONT datasets, with far more of the dataset assigned in HiFi (81–90%) versus ONT (26–41% for ONT R10.3, 59–68% for ONT Q20).

**Figure 2.**
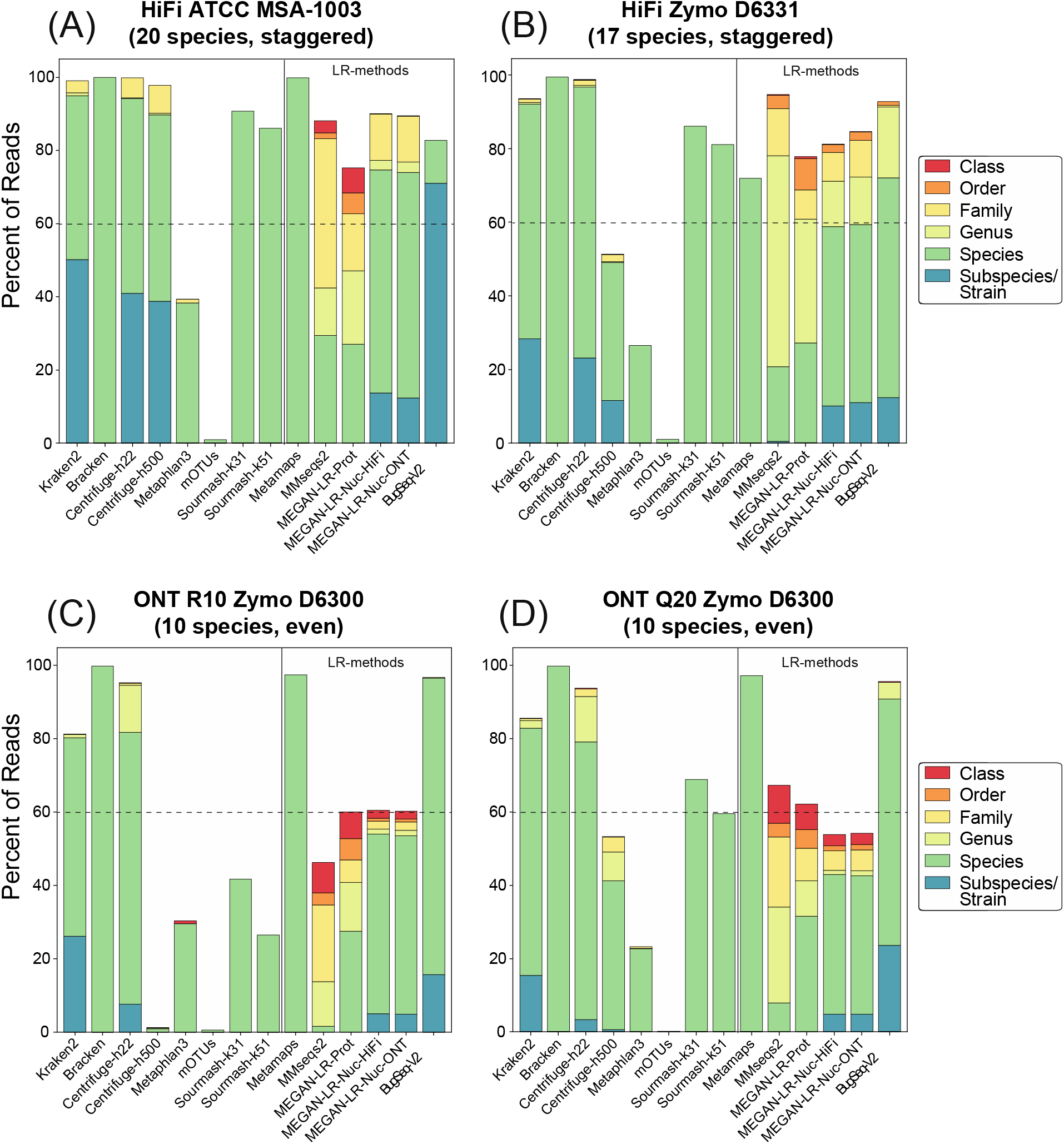
Read utilization for (A) HiFi ATCC MSA-1003, (B) HiFi Zymo D6331, (C) ONT R10 Zymo D6300, and (D) ONT Q20 Zymo D6300. The stacked barplots show the total percent of reads that were assigned to taxonomy. Different colors show the percentage of reads assigned to specific taxonomic ranks.

There was considerable variation in read assignments across the long-read methods and across different sequencing technologies (Fig. 2). Total read assignment in the HiFi datasets ranged from 71–99% (average = 85%) across all long-read methods, and for ONT ranged from 46–97% (average = 71%). For the ONT datasets, MetaMaps and BugSeq-V2 assigned the greatest number of reads (95–97%), with all other methods assigning fewer reads (46–67%). Methods that rely on translation alignments to protein references assigned more reads in the HiFi datasets versus ONT datasets, including MMseqs2 (HiFi: 94–99%; ONT: 46–67%) and MEGAN-LR-prot (HiFi: 71–74%; ONT: 60–62%) (Fig. 2). There were no clear differences in total read assignment for MEGAN-LR-nuc-HiFi and MEGAN-LR-nuc-ONT within the ONT datasets or the HiFi datasets, suggesting read assignment was not sensitive to different minimap2 settings. The MEGAN-LR-nuc methods resulted in a higher number of reads assigned in HiFi datasets (81–90%) versus ONT datasets (54–60%). BugSeq-V2 assigned more reads in the ONT datasets (95–96%) versus HiFi datasets (82–93%). As expected, methods using LCA algorithms during assignment (MMseq2, all three MEGAN-LR workflows, BugSeq-V2) displayed a significant proportion of annotations to taxonomic ranks above the strain and species level (Fig. 2). However, the MEGAN-LR-nuc methods showed a smaller proportion of reads assigned to higher ranks, relative to the protein-alignment methods.

#### Detection Metrics

The complete set of read counts per dataset used in the species and genus-level analyses are provided in Supplementary Tables S1–S8. Detection at different thresholds follows the minimum read counts in Table 3. Species and genus level results are provided for each dataset in Figures 3 and 5 and Table 4. Averaged results per method across all datasets are shown in Figures 4 and 6, and technology specific results are shown in Supplementary Figures S2 and S3.

**Figure 3.**
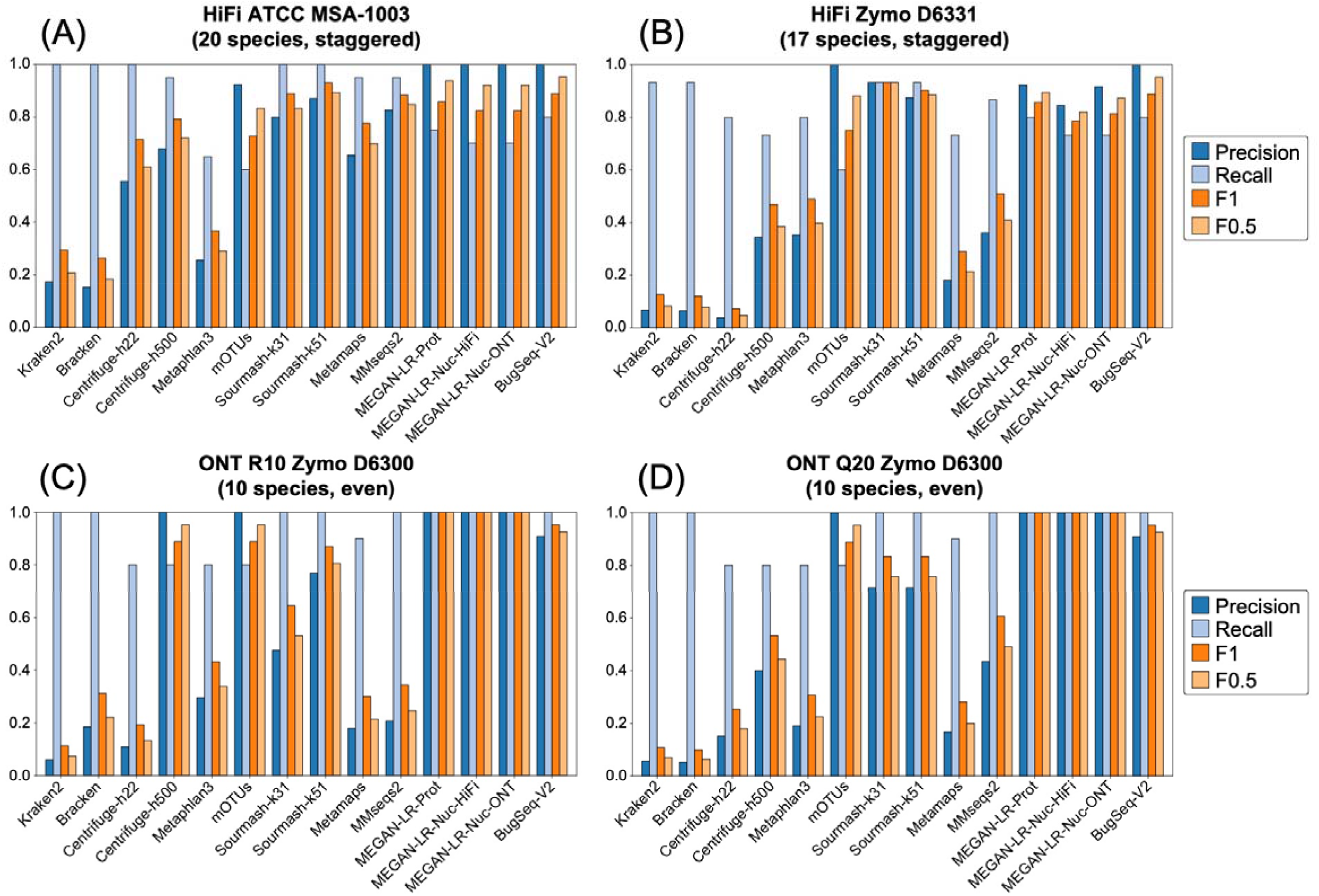
Precision, recall and F-scores for the species-level analysis based on a minimum threshold of 0.001% of the total reads for (A) HiFi ATCC MSA-1003, (B) HiFi Zymo D6331, (C) ONT R10 Zymo D6300, and (D) ONT Q20 Zymo D6300.

**Figure 4.**
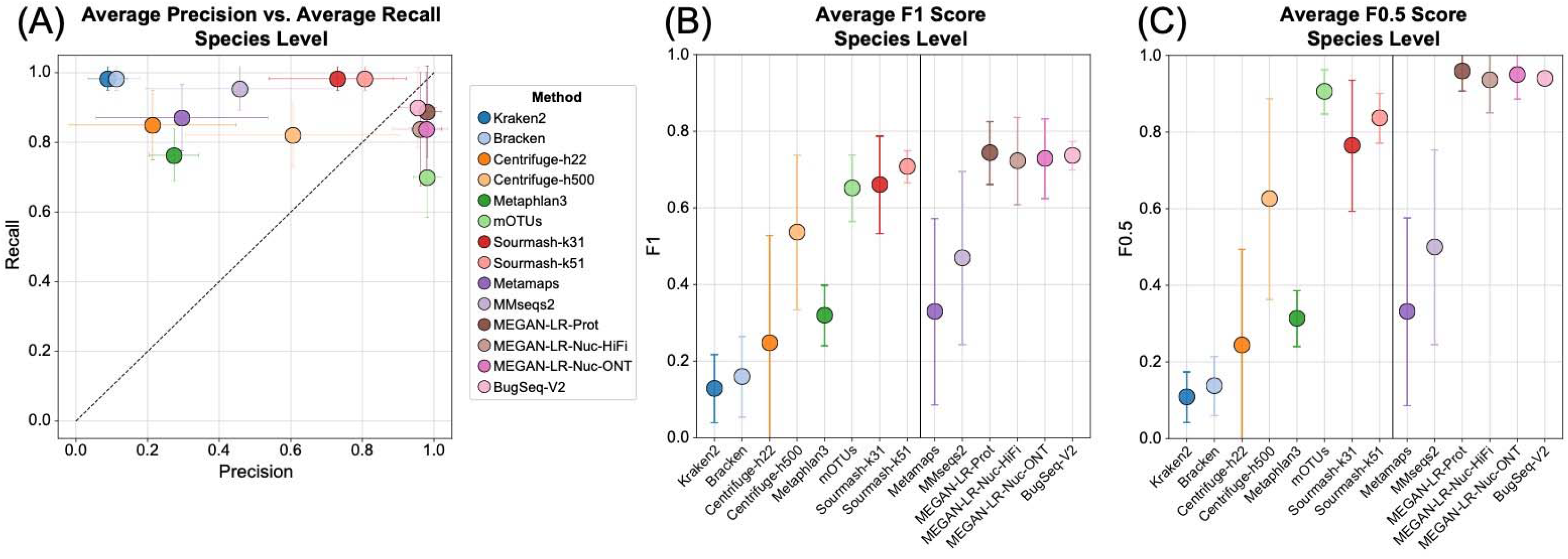
The average values for (A) precision and recall, (B) F1 scores, and (C) F0.5 scores for the species-level analysis based on a minimum threshold of 0.001% of the total reads. Methods to the right of the vertical line in (B) and (C) are the long-read classifiers.

**Figure 5.**
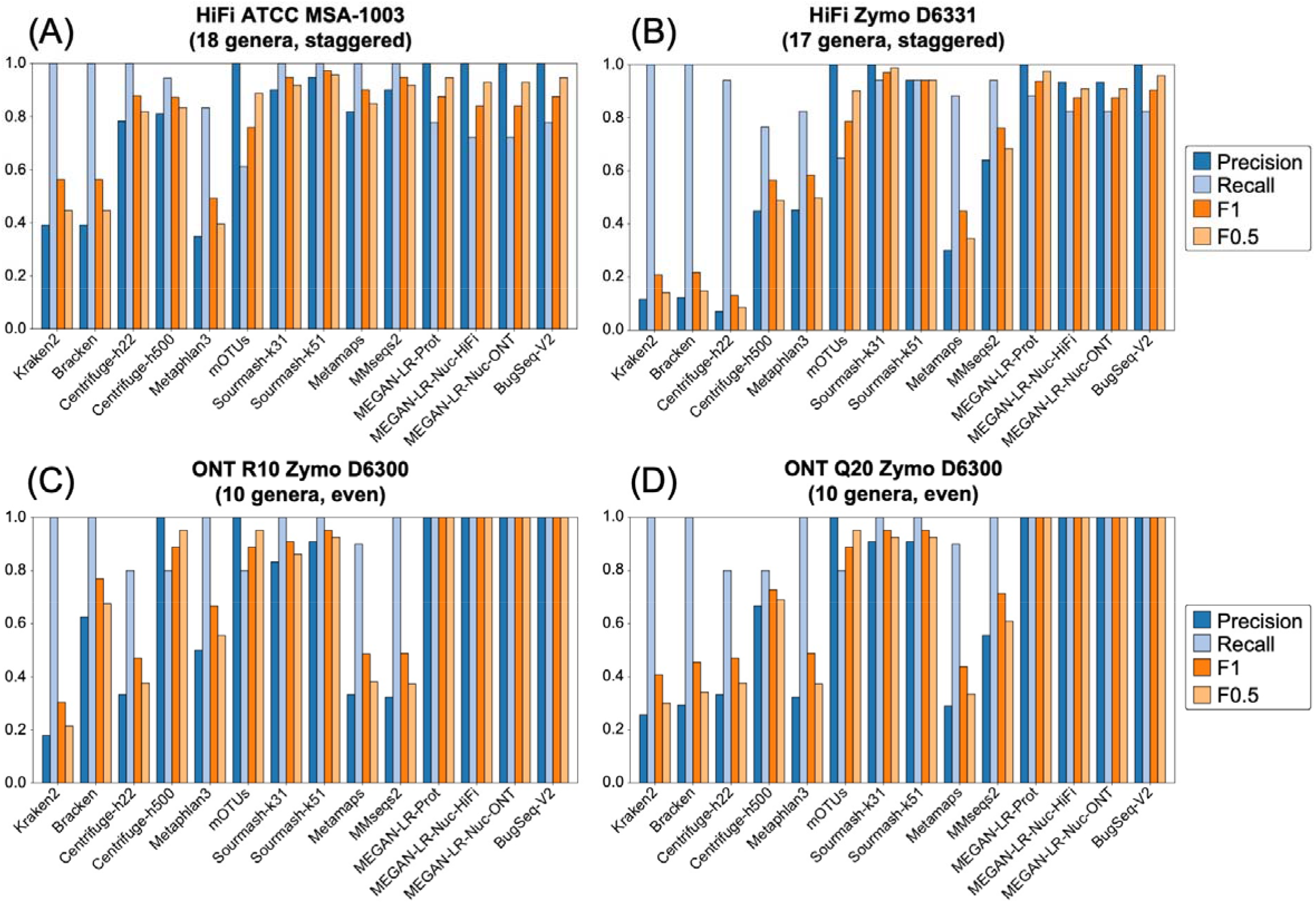
Precision, recall and F-scores for the genus-level analysis based on a minimum threshold of 0.001% of the total reads for (A) HiFi ATCC MSA-1003, (B) HiFi Zymo D6331, (C) ONT R10 Zymo D6300, and (D) ONT Q20 Zymo D6300.

**Figure 6.**
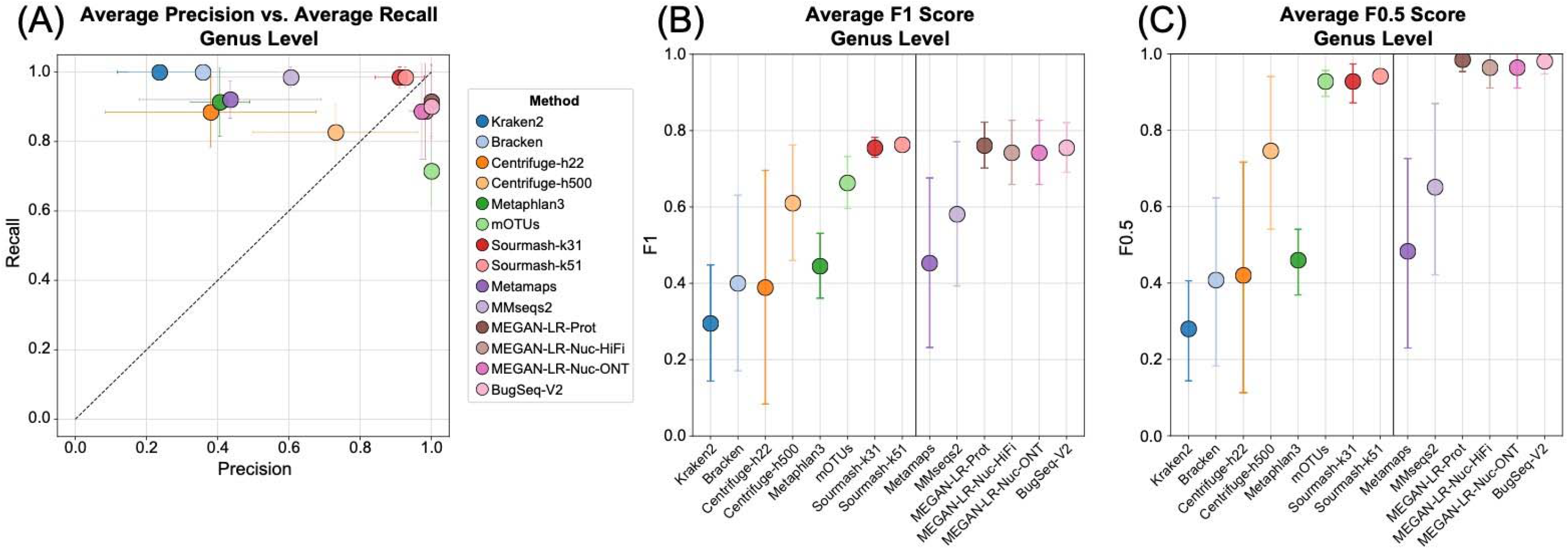
The average values for (A) precision and recall, (B) F1 scores, and (C) F0.5 scores for the genus-level analysis based on a minimum threshold of 0.001% of the total reads. Methods to the right of the vertical line in (B) and (C) are the long-read classifiers.

**Table 4.**
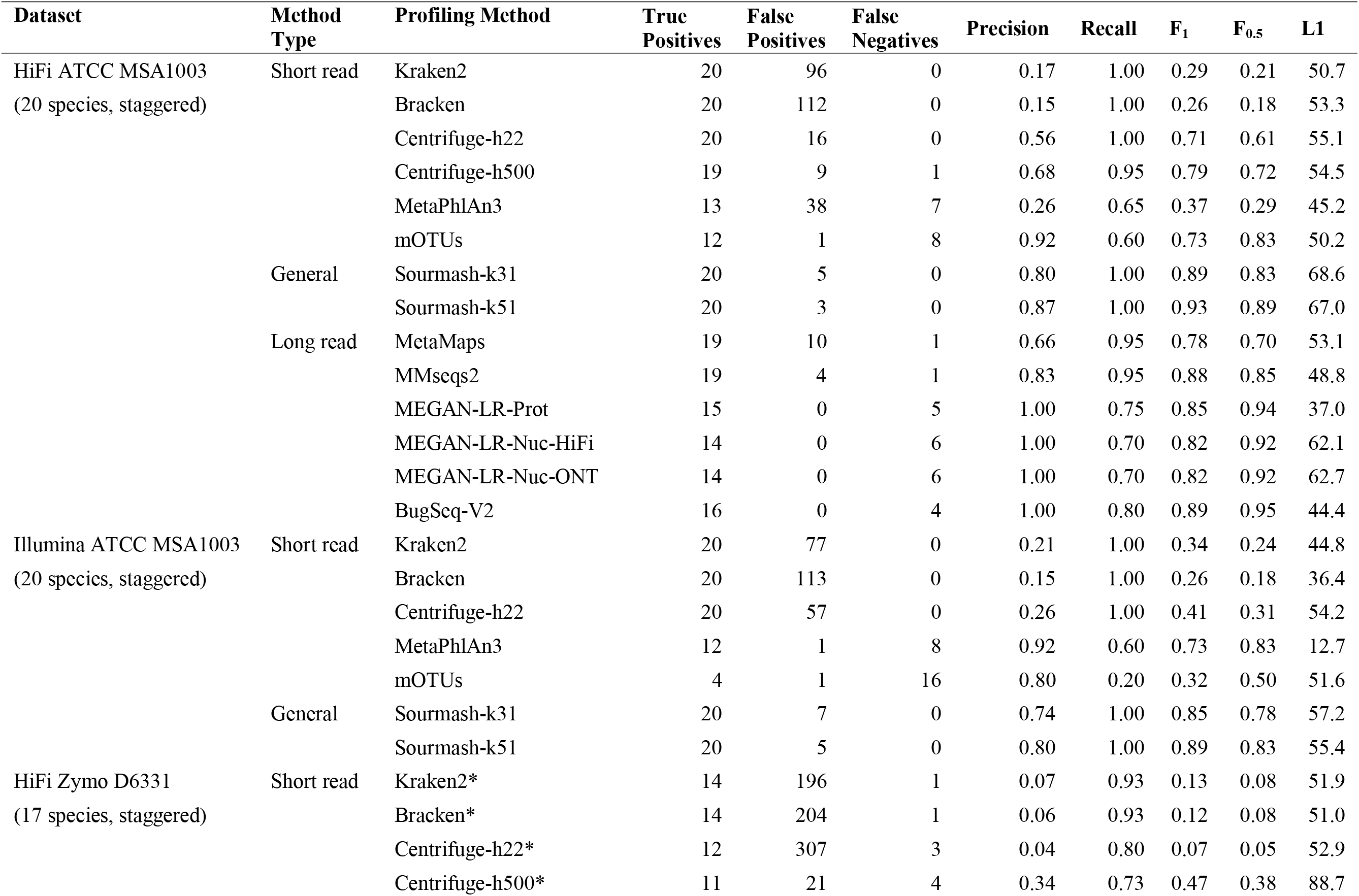
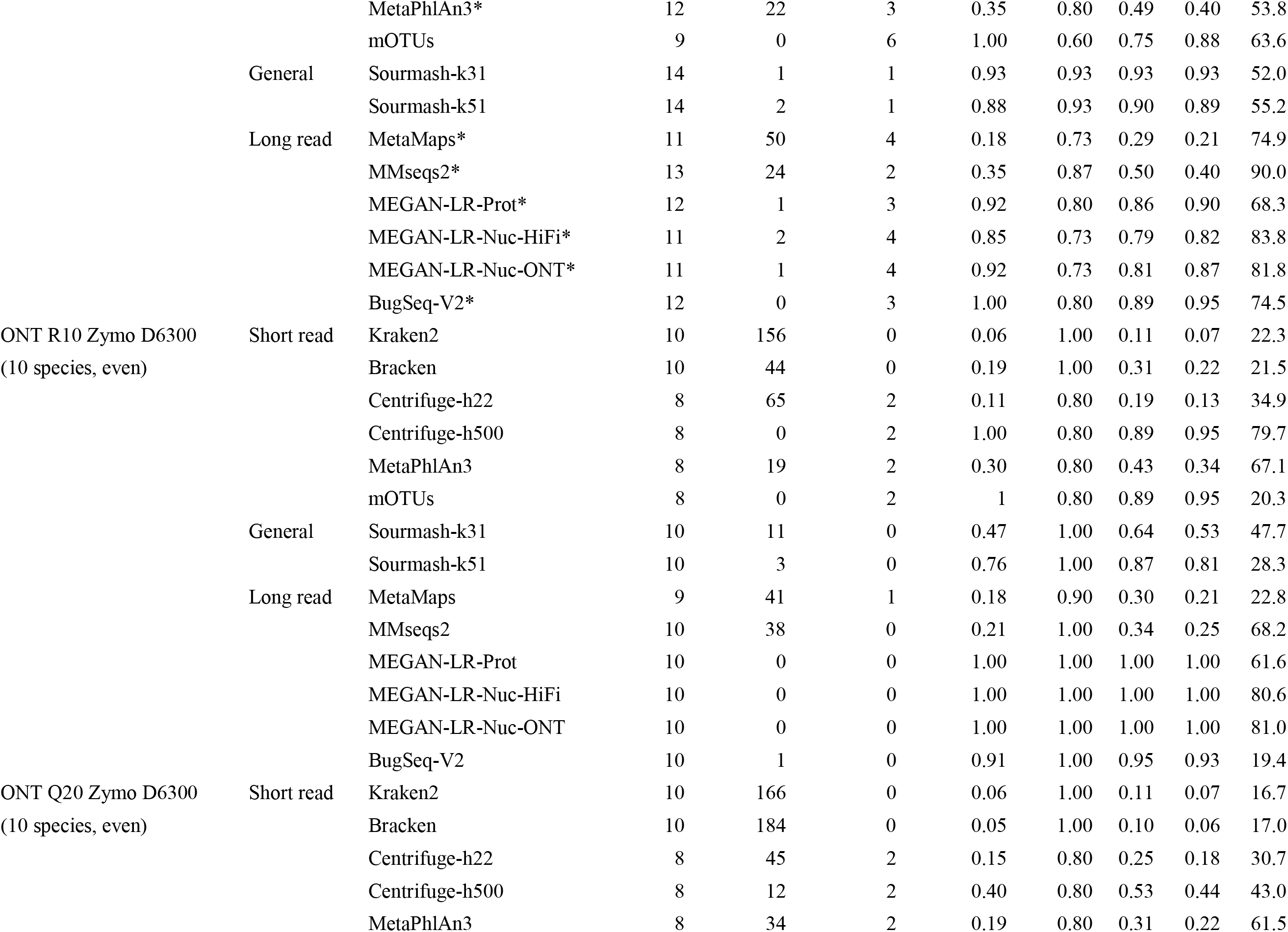
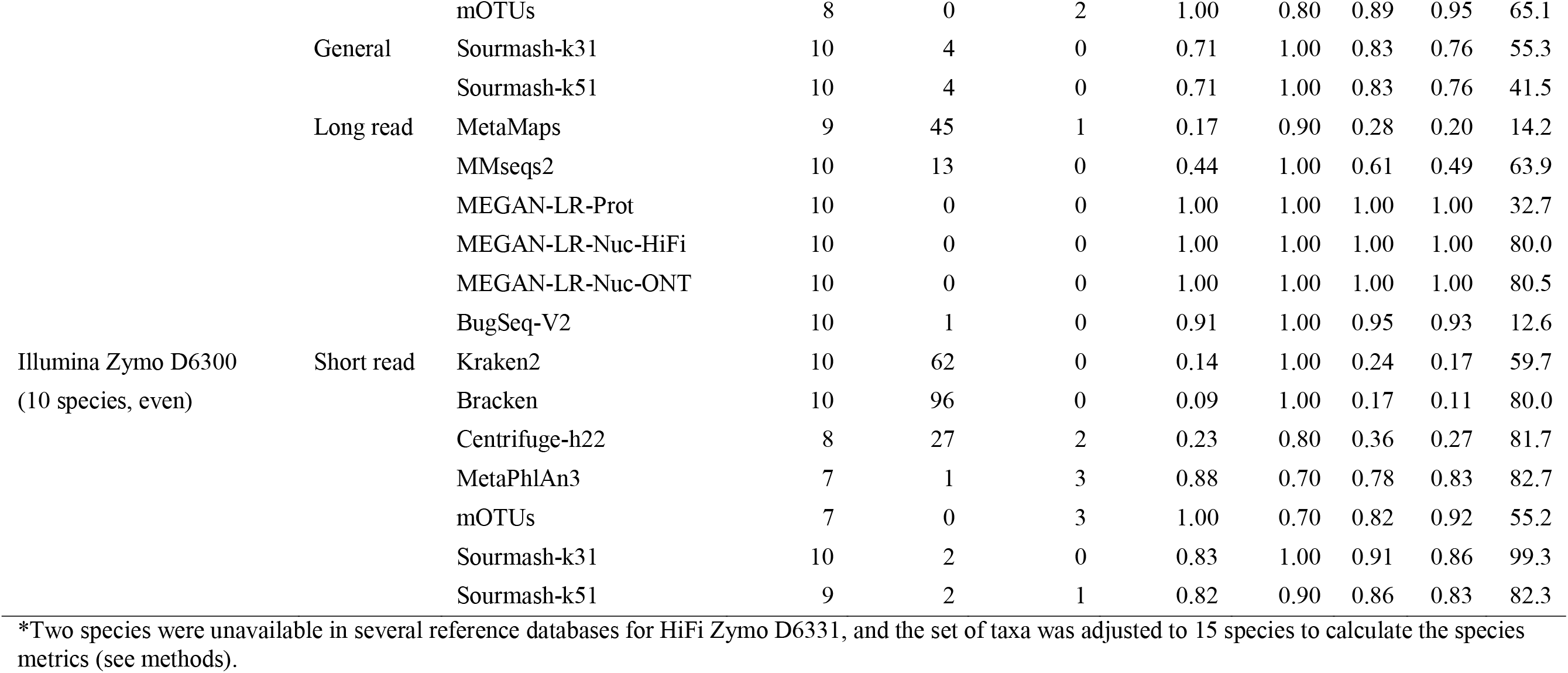
Species-level detection results based on the minimum 0.001% of total reads threshold.

The species-level detection results based on the minimum threshold of 0.001% of the total reads are summarized in Figures 3 and 4 and Table 4. The clearest difference in performance occurs between short-read and long-read/generalized methods (including sourmash). The short-read methods display very low precision and relatively high recall, and consequently very low F-scores (Figs. 3, 4). These results for precision and F-scores are driven by the large number of false positives detected (40–300) despite the presence of few false negatives (Table 4). We note that Bracken did not significantly improve the results of Kraken2, based on these measures (Figs. 3, 4). The Centrifuge-h500 analysis, which required longer matches, resulted in a lower number of false positives and consequently higher precision (Fig. 3, Table 4), though this improvement varied considerably across datasets (Fig. 4). MetaPhlAn3 displayed values that were intermediate between Centrifuge-h500 and the other short-read methods. An exception to this rule occurs with mOTUs2, which displays high precision and moderate recall (Figs. 3, 4). By precision and F-scores, mOTUs2 outperforms all other short read methods by a considerable margin.

The long-read methods and sourmash outperformed the short-read methods in terms of precision, recall, and F-scores (Fig. 3, Table 4), but they also displayed variation in performance. Some methods did not show consistent results and performed better for a particular dataset. For example, MetaMaps and MMseqs2 performed quite well for HiFi ATCC MSA-1003. However, these two methods performed worse for the other three datasets and more closely resembled the results for the short-read methods (e.g., very low precision, higher recall; Fig. 3, Table 4). Interestingly, sourmash displayed high precision and recall for HiFi datasets (highest in k51), outperforming most long-read methods (Fig. 3, Supplementary Fig. S2). However, its performance decreased for the ONT datasets; this is particularly noticeable for ONT R10 (Fig. 3, Supplementary Fig. S3). Across all four datasets, MEGAN-LR-prot, MEGAN-LR-nuc-HiFi, MEGAN-LR-nuc-ONT, and BugSeq-V2 consistently displayed the best performance (Figs. 3, 4). These four methods detected most species in the communities (e.g., low false negatives) and rarely called any false positives (0–2). Consequently, they display high precision, moderate to high recall, and the highest F-scores (Fig. 3). The moderate recall scores for the HiFi datasets resulted from the failure to detect species at lower abundances, particularly for the 0.02% to 0.0001% abundance levels (Supplementary Table S9). Sourmash (k31 and k51) displayed exceptional recall for these challenging HiFi datasets, detecting all species at 0.02% and 0.001% relative abundance (Supplementary Table S9). For the ONT datasets, the species in Zymo D6300 had comparatively high abundances (12% and 2%), and this was reflected in perfect recall for nearly all long-read methods as well as sourmash (Fig. 3, Table 4). We did not observe any difference in performance between MEGAN-LR-nuc-HiFi and MEGAN-LR-nuc-ONT for the ONT datasets or HiFi datasets, suggesting the profiling analyses are not sensitive to minimap2 alignment settings.

The genus-level analysis based on the minimum threshold of 0.001% of the total reads largely mirrored the species-level results, but with expected improvements in precision, recall, and F-scores (Figs. 5, 6, Supplementary Table S10). Improvements were nearly guaranteed because reads assigned to multiple species within a genus are all considered correct at the genus level, and consequently the number of false positives (and potentially false negatives) decreased. Despite improvements in precision, recall, and F-scores across all methods at the genus level, the long-read methods still outperformed most short-read methods by a considerable margin (Fig. 4, Supplementary Table S10). We observed perfect precision in mOTUs2, but it displayed lower recall relative to long-read methods (Fig. 6). Sourmash (k31 and k51) displayed perfect recall and precision was comparable to the long-read methods (particularly for HiFi datasets, Figs. 5, 6, Supplementary Figure S2).

Requiring a moderate minimum threshold for detection (0.1% of total reads) for the species-level analysis had an overall positive effect on precision, but negative effect on recall (Supplementary Fig. S4, Supplementary Table S11). These changes were most dramatic for the short-read methods, in which the number of false positives was reduced from several hundred to ∼10 or fewer, thereby increasing precision considerably (Supplementary Table S11). However, despite this improvement the long-read methods still performed better in terms of precision and F-scores (Supplementary Fig. S4). Precision increased for some long-read methods (MetaMaps, MMseqs2), but others were unaffected as they were already high at the lower detection threshold. As expected, this increase in minimum detection threshold most strongly impacted recall in the communities with staggered abundances (HiFi datasets) versus communities with even abundances (ONT datasets). In the HiFi datasets, the long-read methods displayed more false negatives which resulted in lower recall (Supplementary Fig. S8). At the 0.1% total reads detection threshold, all methods (long and short) failed to detect species with <0.02% abundance and missed several species with 0.1–1.8% abundance (Supplementary Table S12). Surprisingly, this detection threshold also reduced the recall of some methods for the ONT datasets, with a more noticeable reduction in recall values for ONT R10 Zymo D6300 (Supplementary Fig. S4, Supplementary Table S11). The patterns for the genus-level analysis using the 0.1% total reads detection threshold mirrored the species-level results (Supplementary Fig. S5). Precision increased in the short-read methods across all datasets, and recall was lowered in the staggered abundance communities (Supplementary Table S13).

The highest minimum threshold for detection used in our experiment (1% of total reads) exacerbated the effects described for the 0.1% detection threshold. The most noticeable effects were for the communities with staggered abundances: all methods displayed perfect precision (with one exception), but recall was drastically lowered (<0.6; Supplementary Fig. S6, Supplementary Table S14). In other words, false positives were completely eliminated, but at the cost of vastly increased false negatives. Using 1% of total reads as the minimum detection threshold for HiFi ATCC MSA-1003 and Zymo D6331, all methods (long and short) failed to detect species with <1.8% relative abundance, and some species were not detected in the 1.5% and 6% abundance levels (Supplementary Table S15). This higher threshold for detection also impacted results for the even abundance communities (ONT R10 and Q20 for Zymo D6300). Precision increased primarily for the short-read methods, yet perfect precision was not achieved by all methods (Supplementary Fig. S6, Supplementary Table S14). This higher detection threshold also caused recall to drop (<0.8) in these datasets for all methods except Kraken2, Bracken, and one instance of BugSeq V2, each of which maintained perfect recall (Supplementary Fig. S6). This indicates that multiple methods failed to detect several species at the 2% and 12% abundance levels in Zymo D6300. These effects were mirrored in the genus-level analysis with the 0.1% detection threshold (Supplementary Fig. S7, Supplementary Table S16).

#### Relative Abundance Estimates

The species-level and genus-level relative abundances are shown in Figures 7 and 8, respectively. The results of the chi-squared goodness of fit tests (GOF) are reported in Supplementary Tables S17 and S18 and highlighted in Figures 7 and 8. The L1 scores are reported in Table 4 and Supplementary Tables S10, S19, S23, and S27. At the species level, abundance estimates by the long-read methods and sourmash were more accurate than those produced by short-read methods across all datasets (based on L1 distances and chi-squared test statistic values). For HiFi ATCC MSA-1003, MetaMaps, MMseqs2, MEGAN-LR-prot, and BugSeq-V2 all passed the GOF, and BugSeq-V2 had the lowest error. All methods failed the GOF for HiFi Zymo D6331 at the species level (which had two species missing from most databases, see methods), but MEGAN-LR-prot and BugSeq-V2 resulted in the lowest error. For ONT R10 Zymo D6300, mOTUs2, sourmash-k51, and BugSeq-V2 passed the GOF. Both BugSeq-V2 and MEGAN-LR-prot passed the GOF for ONT Q20 Zymo D6300. At the genus level we generally found more methods passed GOF for each dataset, except for HiFi Zymo D6331 for which only sourmash (k31 and k51) and BugSeq-V2 passed (Supplementary Table S18). All methods that accurately estimated abundances at the species level also passed the GOF at the genus level (Figs. 7, 8). We additionally found Centrifuge (h22 and/or h500) and MetaMaps passed GOF at the genus level in some datasets in which they failed at the species level (Figs. 7, 8). Across all datasets and levels, we generally found that BugSeq-V2 had the lowest abundance error, followed closely by MEGAN-LR-prot (Supplementary Tables S17, S18). Across datasets, the proportion of reads assigned to false positives (‘Other’, Figs. 7, 8) was generally highest for MetaPhlAn3, followed by Kraken2 and Bracken.

**Figure 7.**
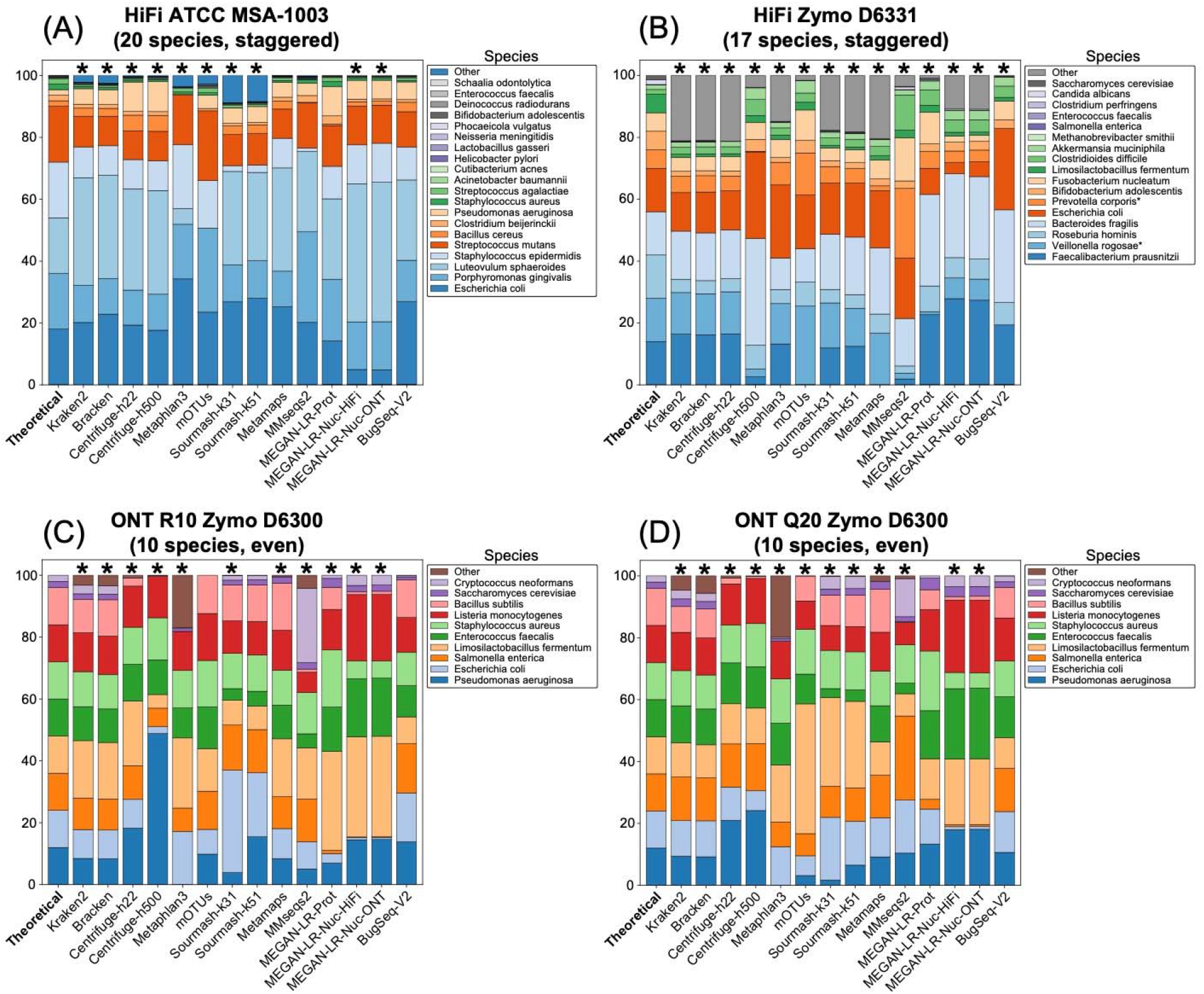
Species-level relative abundance estimates for (A) HiFi ATCC MSA-1003, (B) HiFi Zymo D6331, (C) ONT R10 Zymo D6300, and (D) ONT Q20 Zymo D6300. The theoretical distributions are shown on the left and are based on the manufacturer’s specifications. The read counts for all species-level false positives were grouped in a category labeled ‘Other’. For HiFi Zymo D6331, all species assignments within the genera *Prevotella* and *Veillonella* were counted towards *Prevotella corporis* and *Veillonella rogosae*, due to the absence of these species from several databases (see methods). Asterisks signify methods that failed the chi-squared goodness of fit test (e.g., the abundance estimates were significantly different from the theoretical values).

**Figure 8.**
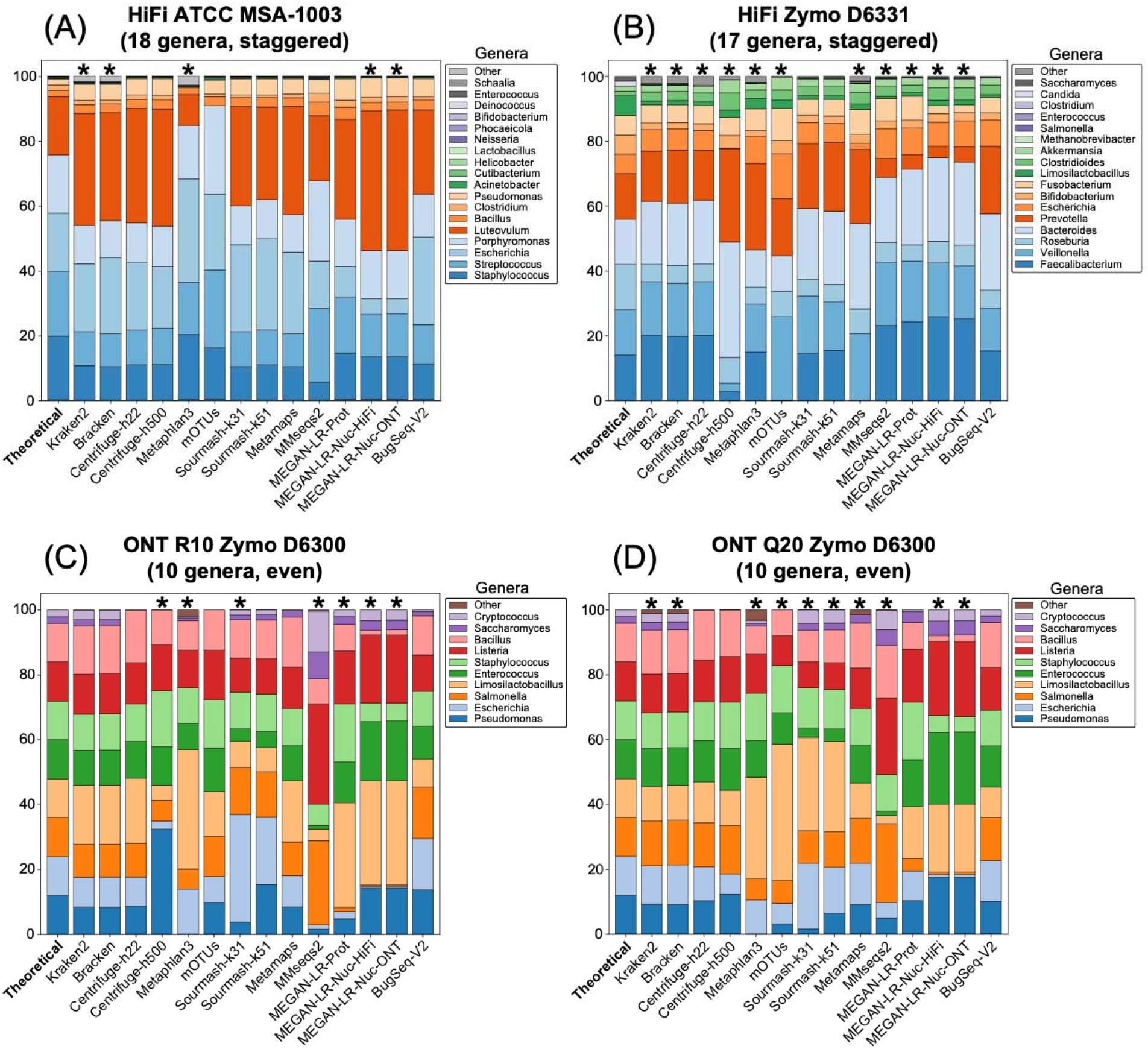
Genus-level relative abundance estimates for (A) HiFi ATCC MSA-1003, (B) HiFi Zymo D6331, (C) ONT R10 Zymo D6300, and (D) ONT Q20 Zymo D6300. The theoretical distributions are shown on the left and are based on the manufacturer’s specifications. The read counts for all genus-level false positives were grouped in a category labeled ‘Other’. Asterisks signify methods that failed the chi-squared goodness of fit test (e.g., the abundance estimates were significantly different from the theoretical values).

#### Analyses of Shorter ONT Reads

Comparisons of the length-filtered variations of each ONT dataset revealed that shorter reads (< 2kb) negatively impacted taxonomic profiling analyses. For each ONT dataset, we created a primary dataset which contained only longer reads (> 2kb) and a secondary dataset which had a large proportion of shorter reads (< 2kb; see methods). In the primary datasets, precision and F-scores were very high for long-read methods and low for short-read methods at the 0.001% reads detection threshold. In the secondary datasets, precision and F-scores were comparatively lower for the long-read methods and were similarly low for the short-read methods (Supplementary Fig. S8, Tables S19, S20). Based on Wilcoxon Signed-Rank tests, the observed differences in precision and F-scores between the primary and secondary datasets were not statistically significant. However, at the 0.1% reads detection threshold we found precision and F-scores were substantially lower in the secondary datasets at both the species and genus level, across all methods (Supplementary Fig. S8, Tables S19, S20). These differences in precision and F-scores were statistically significant (p < 0.01 for all comparisons). In contrast to most methods, BugSeq produced relatively consistent results in precision and F-scores between the primary and secondary datasets across the different filtering thresholds.

Relative abundance estimates appeared heavily skewed in the secondary datasets, and most methods greatly overestimated the abundance of *Limosilactobacillus fermentum* in the community (Supplementary Fig. S9). Interestingly, in the secondary datasets the abundance error at the species level decreased for the short-read methods but increased in the long-read methods. At the genus level, abundance error appeared to increase across all methods in the secondary datasets. Based on Wilcoxon Signed-Rank tests, we did not find a significant difference in abundance error between the primary and secondary datasets at the species level, but at the genus level abundance error was significantly higher in the secondary datasets (p < 0.05 for the R10 and Q20 comparison). In the secondary datasets, nearly every method failed the chi-squared goodness of fit test at the species level (21 of 22) and genus level (20 of 22; Supplementary Tables S21, S22). We found BugSeq and Centrifuge-h22 passed the GOF for the species level of ONT R10 Short, and BugSeq passed the GOF for ONT R10 Short at the genus level (Supplementary Tables S21, S22). No methods passed the GOF for ONT Q20 Short at the species or genus level.

#### Analyses of Illumina and Artificial Short Reads

We evaluated the performance of Kraken2, Bracken, Centrifuge-h22, MetaPhlAn3, mOTUs2, and sourmash (k31 and k51) for two types of short-read datasets for the ATCC MSA-1003 and Zymo D6300 mock communities. We found detection and abundance results were highly similar between the Illumina short-read datasets and the “simulated” short-read datasets (SR-Sim; which were derived from the long reads). This indicates that for short-read methods, the differences in results between the long-read datasets and the Illumina short-read datasets are unlikely to be driven by platform-specific or confounding effects (such as DNA extraction methods or error profiles). However, the fraction of dataset assigned using sourmash was quite different between the Illumina (94–96%) and the SR-Sim ONT dataset (62.9–72.6%) for Zymo D6300. The SR-Sim ONT was created from the ONT Q20 long reads, and we note sourmash also assigned a comparable fraction of reads in the full length ONT Q20 dataset (59–68%). These results suggest that error profile impacts sourmash profiling performance.

The precision, recall, and F-score values obtained from the short-read datasets strongly resembled those obtained from long reads for both communities (Figs. 9, 10, Table 4, Supplementary Figure S10, Supplementary Tables S23–24, S27–28). This overall pattern included low precision and high recall for Kraken2, Bracken, and Centrifuge-h22. MetaPhlAn3 improved in performance, with high precision and moderate recall, comparable to mOTUs2. Sourmash was the top performer in the short-reads datasets with perfect recall and high precision (Figs. 9, 10). More stringent filtering (0.1% or 1% of total reads) dramatically reduced false positives for Kraken2, Bracken, and Centrifuge-h22, but also negatively impacted recall (Supplementary Table S23), and in many cases produced scores that were worse than the long-read scores for these method and filtering combinations (Supplementary Table S11, S14). The same patterns were present for the genus-level analyses of the short-read datasets of ATCC MSA-1003 (Supplementary Table S24) and the less complex ZymoD6300 community (10 species).

**Figure 9.**
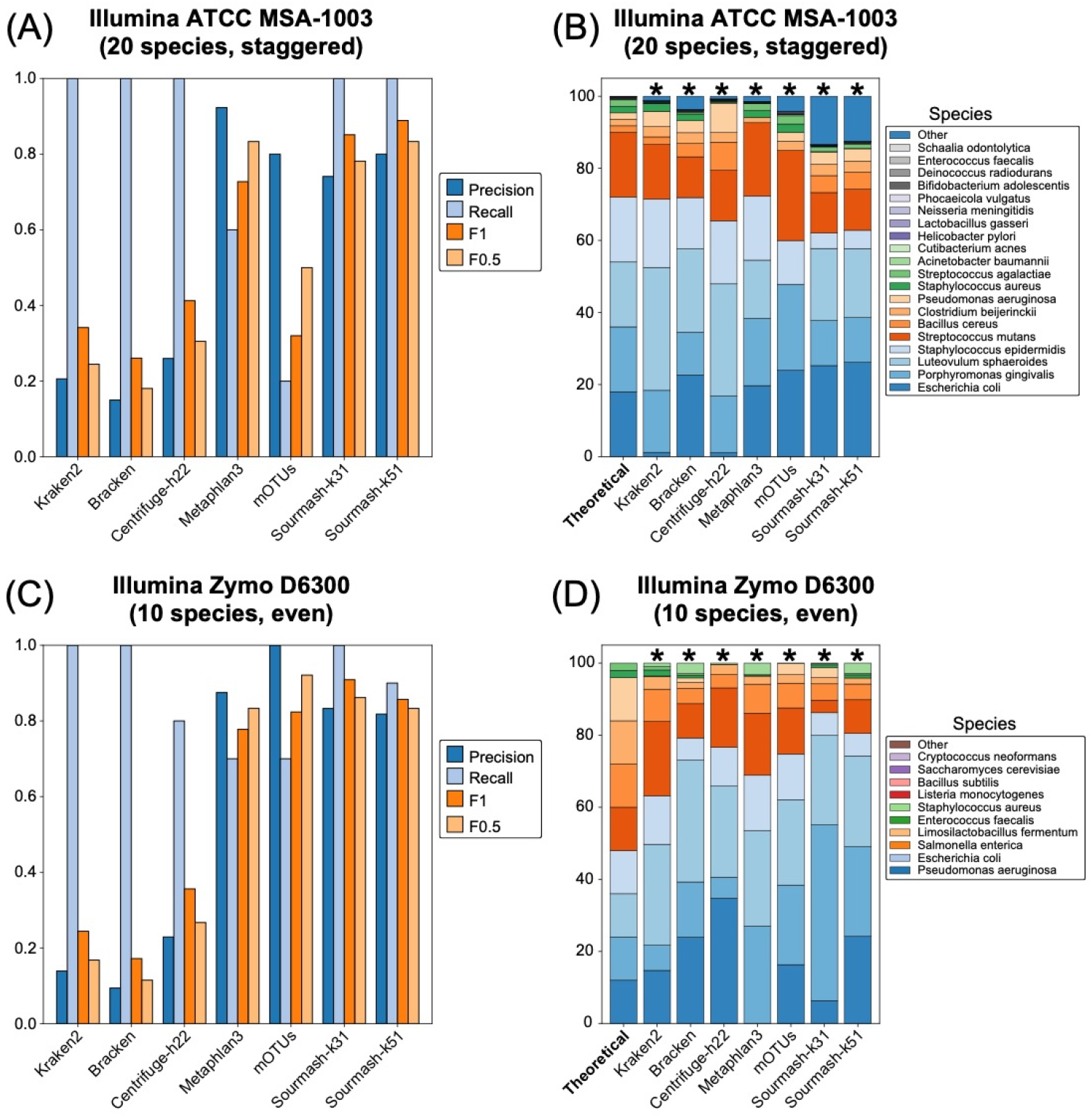
Results for the two Illumina short-read datasets. Precision, recall and F-scores for the species-level analysis based on a minimum threshold of 0.001% of the total reads for (A) Illumina ATCC MSA-1003 and (B) Illumina Zymo D6300. Species-level relative abundance estimates for (C) Illumina ATCC MSA-1003 and (D) Illumina Zymo D6300. The theoretical distributions are shown on the left and are based on the manufacturer’s specifications. The read counts for all species-level false positives were grouped in a category labeled ‘Other’. Asterisks signify methods that failed the chi-squared goodness of fit test.

**Figure 10.**
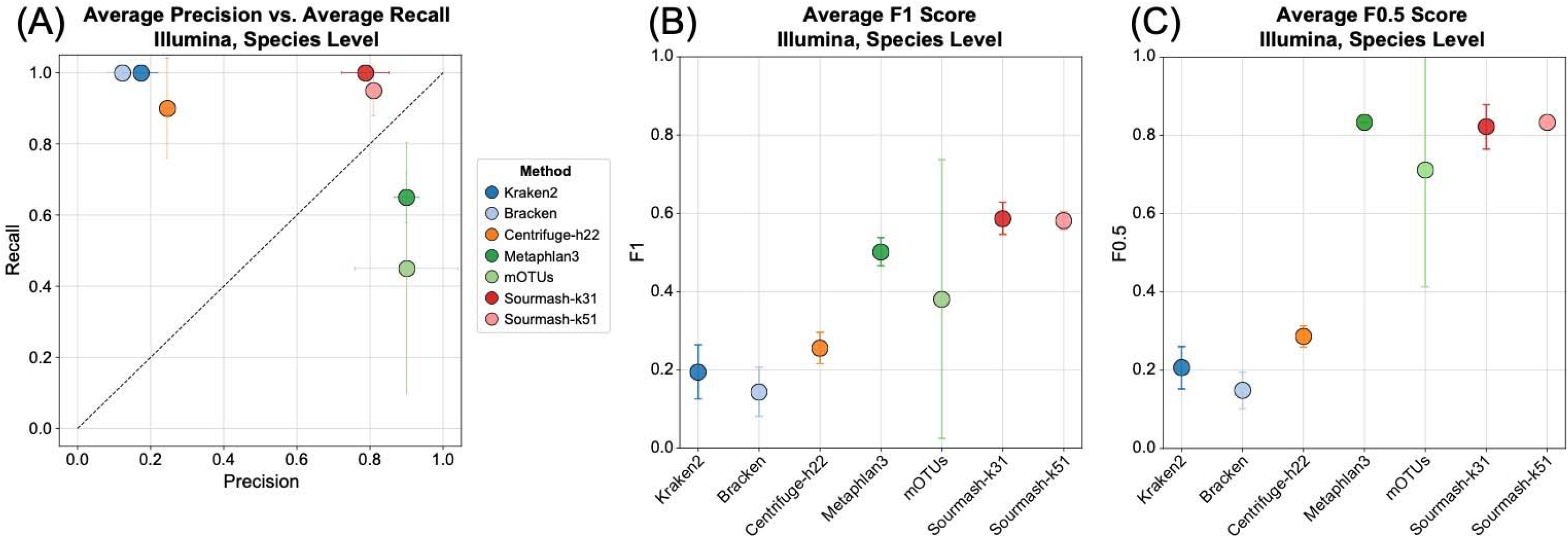
Results for the two Illumina short-read datasets. The average values for (A) precision and recall, (B) F1 scores, and (C) F0.5 scores for the species-level analysis based on a minimum threshold of 0.001% of the total reads.

The short-read datasets failed to produce accurate relative abundance estimates (Fig. 9, Supplementary Figures S11–12, Supplementary Tables S25–26, S29–30). All short-read methods failed the chi-squared goodness of fit test at the species level in both communities, and at the genus level only sourmash-k51 passed the goodness of fit test across multiple datasets (Supplementary Figure S12).

## Discussion

With decreasing error rates in long reads and the recent introduction of new long-read read profiling methods, long reads are increasingly utilized for metagenomic applications. We used publicly available mock community datasets to perform a critical assessment of taxonomic profiling methods for long-read datasets, including five long-read methods, five short-read methods, and one generalized method. While all methods displayed some trade-offs between precision and recall, our results suggest that generalized methods (e.g., sourmash) and methods designed for long reads performed best.

In our study, we included a mix of short-read classifiers (Kraken2, Centrifuge), short-read profilers (Bracken, MetaPhlAn3, mOTUs2), a generalized profiler (sourmash), and several long-read classifiers (MetaMaps, MMSeqs2, BugSeq, MEGAN-LR-prot, MEGAN-LR-nuc-HiFi, and MEGAN-LR-nuc-ONT). The ideal taxonomic classifier or profiler should display high precision and recall. We found that the methods examined here tended to fall into three broad categories: 1) high precision and moderate recall, 2) moderate precision and high recall, and 3) low precision and high recall (Fig. 3A). The first two categories provide the best tradeoffs, with the third category displaying undesirable properties. Overall, we find that BugSeq, MEGAN-LR­prot, and MEGAN-LR-nuc provide the best tradeoffs for all long-read metagenomics data. In addition to these three, sourmash was also a top-performing method for HiFi datasets. Below, we discuss our findings for short-read, long-read, and generalized methods, including tradeoffs, best practices, and the impact of shorter reads. Finally, we briefly summarize the effects of read accuracy on method performance.

### Short-read methods

A majority of short-read methods (Kraken2, Bracken, Centrifuge-h22) assigned a high proportion of reads and displayed high recall, but they produced poor abundance estimates. They also recovered a very high number of false positives (15–300 species) and consequently had very low precision and F-scores (Figs. 2–4). False positives were not a trivial proportion of assigned reads; they comprised up to 25% of the reads assigned at the species level (Fig. 7). We attempted to apply long-read settings to Centrifuge (Centrifuge-h500) to improve detection results. Unfortunately, this setting reduced total read assignment and had unpredictable outcomes on precision, recall, and F-scores across the datasets (Figs. 2–4). The marker-based profilers had variable performance. MetaPhlAn3 displayed low precision and moderate recall, whereas mOTUs2 displayed high precision with comparable recall (Fig. 4). Both methods assigned a low percentage of reads, which is typical for marker-based mapping methods. Previous studies have shown similar results for these methods with short-read datasets [3, 8, 9], but here we demonstrate the use of long reads does not significantly change these trade-offs.

We attempted to improve the results from short-read methods using various levels of filtering. Specifically, we applied different minimum thresholds for detection (0.001%, 0.1%, and 1% of the total reads) in an effort to reduce false positives and improve precision. A moderate detection threshold (0.1% total reads) successfully reduced the false positive count of species from hundreds to fewer than 15, and without significantly reducing recall. However, precision in these methods was still below scores produced by the long-read methods without any filtering. A stringent detection threshold (1% total reads) greatly improved precision for many short-read methods, but severely impacted recall by eliminating detection of many species at lower abundance levels (<2% abundance). Overall, we found that filtering was necessary to reduce false positives and improve precision in the short-read methods. However, none of the filtering strategies successfully balanced precision and recall to produce results similar to the long-read methods.

We analyzed short read Illumina datasets for two of the mock communities to evaluate if any short-read methods performed differently. We found consistent results across short and long-read datasets for Kraken2, Bracken, and Centrifuge (high false positives, low precision). For these methods, the outcomes appear to be driven by characteristics of the methods themselves, rather than read type. However, we observed an improvement in MetaPhlAn3 (higher precision), indicating this method is potentially sensitive to the read type. We could not appropriately evaluate differences mOTUs2 because the “long read” analyses consisted of short reads derived from the long reads, meaning the inputs for both the short and long-read analyses were highly similar.

### Long-read and generalized methods

Several long-read profiling methods showed consistent and favorable characteristics across all datasets. These include MEGAN-LR-prot, MEGAN-LR-nuc (both mapping settings), and BugSeq, which displayed medium to high read assignment and very high precision (Figs. 2, 5, Table 4). Recall values from these methods differed between the staggered abundance and even abundance communities (0.7–0.8 and 1, respectively). This difference is explained by the failure to detect species with <0.02% abundance in the staggered community. In contrast to the short-read methods, several long-read methods estimated accurate species abundances for the complex communities (particularly ATCC MSA-1003; Fig. 7). Across all communities, we generally found BugSeq displayed the lowest abundance error of any method, followed by MEGAN-LR-prot. Though abundance error was higher for Metamaps, MMseqs2, and MEGAN-LR-nuc, these methods still performed better than most short-read methods in most cases. We found that MetaMaps and MMseqs2 showed high read assignment and precision for one dataset (HiFi ATCC MSA-1003), but for all other datasets showed unfavorable qualities which resembled many short-read methods (e.g., high false positives and low precision, high recall). This contrasts with a recent study by Marić et al. (2020), who found MetaMaps performed better than MEGAN-LR. However, Marić et al. (2020) produced alignments for MEGAN-LR using a different method (LAST) and a reduced database, which may explain these differences. Several long-read methods displayed high or perfect precision (MEGAN-LR-prot, MEGAN-LR-nuc, BugSeq), and this did not change after applying a moderate detection threshold (0.1% of total reads). However, we observed a dramatic improvement in precision for MMseqs2 and MetaMaps (Supplementary Fig. S6). This was accompanied by a slight reduction in recall, suggesting this filtering strategy is beneficial for these methods. A more stringent detection threshold (1% total reads) resulted in perfect precision but severely reduced recall for all long-read methods, and is not advised. Overall, we found that filtering was not required for many long-read methods (MEGAN-LR-prot, MEGAN-LR-nuc, BugSeq), and that moderate filtering could be used to balance precision and recall for methods with higher false positive rates (MetaMaps, MMseqs2).

The generalized method, sourmash, also performed consistently well on most datasets, with nearly perfect recall and precision similar to the top performing long-read classifiers. Sourmash k31 only had one false negative in any dataset: *Clostridium perfringens*, which had a theoretical abundance of 0.0001% in Zymo D6331. When sourmash gather was run with default fractional scaling (1/1000 k-mers) but without a detection threshold (any k-mer match is reported), matches were found to 651 *Clostridium perfringens* genomes, with the most k-mer matches to GCA_902166105.1 (Clostridium perfringens strain=4928STDY7387913; 220 k­mers, representing approximately 22,000 bp sequence). This finding suggests that the fractional scaling was sufficient for detection, but the match was eliminated during the greedy minimum­set-cover assignment to best-match genomes. Disambiguating extremely low-abundance genomes from similar genomes truly present in the community represents a challenge for sourmash’s greedy assignment algorithm: most k-mer matches to genomes in the genus *Clostridium* were shared with the *Clostridioides difficile* genome match (1.5% of Zymo D6331), leaving < 10kb of detected sequence that uniquely matched *Clostridium perfringens* genomes, far below the default threshold for sourmash gather (50kb). While zero-threshold gather is too sensitive (yielding many false positives), setting a moderately lowered detection threshold may improve recall of very low-abundance genomes in long-read datasets, particularly as sequencing depth tends to be lower than typical short-read datasets, which sourmash has primarily been tested on.

Sourmash displayed high precision, comparable to long-read classification methods. The majority of species-level false positives results represented different species in the same genus. As k-mer matching is less tolerant of sequence mismatch than alignment methods, these FP matches may represent genomic sequence shared across these species, but with sequence mismatches in the sequenced metagenome compared with the reference species in GenBank.

In terms of dataset utilization, sourmash performed less well for ONT data compared with datasets from other platforms, regardless of read length. This, with the observed improved performance on ONT Q20 compared with R10.3, suggests that the error profile may reduce exact matching of k31 and k51 k-mers to reference genomes. However, sourmash still performed well on ONT community composition and relative abundance, suggesting that ONT datasets provide sufficient non-erroneous k-mers for assignment via the minimum-set-cover approach, and that the error profile does not result in profiling bias across taxa.

### Best Practices and Detection limits

Our findings demonstrate the important trade-offs between precision, recall, and detection limits. Taxonomic profiling methods which have high recall (e.g., they find all the species present in a community) also tend to have low precision (e.g., they recover many false positives). In our experiment, methods with these characteristics include many short-read methods (Kraken2, Bracken, Centrifuge-h22, MetaPhlAn3), and several long-read methods (MetaMaps, MMseqs2). There is one clear exception to this rule – sourmash displays near perfect recall and high precision, particularly in the HiFi datasets (Fig. 3, Supplementary Fig. S2). Sourmash is k-mer-based, similar to Kraken2, Bracken, and Centrifuge, but uses k-mers from across the entire dataset, rather than individual reads, to find best-match genomes. In this way, it is able to leverage longer-range information present in a dataset, though not across reads themselves. By contrast, most other methods which have high precision (e.g., no false positives) tend to have lower recall (e.g., not all species are detected). In our experiment, this was represented by several long-read methods, including MEGAN-LR-prot, MEGAN-LR­nuc, and BugSeq. These three methods involve mapping reads to whole-reference databases, and subsequently interpreting alignments across the entire length of reads. This strongly suggests that top-performing methods are those that can utilize long-range information available in long reads. Although mOTUs2 displays high precision, its current implementation breaks long reads into artificial short reads and eliminates all long-range information, making it less desirable for long-read metagenomics.

If precision is the most important aspect of a long-read metagenomics experiment, we suggest using MEGAN-LR-prot, MEGAN-LR-nuc, or BugSeq, which do not require any additional post-processing or filtering. The choice among them could depend on which references will be used (proteins: MEGAN-LR-prot; nucleotide sequences: BugSeq, MEGAN­LR-nuc), computational skills/resource availability (BugSeq is an online service platform; the MEGAN-LR workflows require high resources and bioinformatics experience), and abundance estimation (BugSeq and MEGAN-LR-prot are considerably more accurate than MEGAN-LR­nuc). One additional advantage of MEGAN-LR-prot is that it simultaneously assigns functional annotations to genes on reads, providing both taxonomic and functional profiles.

There may also be cases where recall is more important for an experiment. For these use-cases we recommend using sourmash, which had the highest recall without reduced precision. With sourmash, we detected all species down to 0.001% relative abundance in the HiFi datasets, with only 2–3 false positives (Table 4, Supplementary Table S9). While this method appears to have reduced precision with ONT data (Supplementary Fig. S3), the genome-level assignments produced during rapid sourmash profiling could be used as candidate genomes for detailed, alignment-based analysis to confirm results and reduce false positives [35]. Other long-read methods with high precision (MEGAN-LR-prot, MEGAN-LR-nuc, BugSeq) had excellent recall for species with higher abundances. These three methods confidently detected species with 0.1% and greater abundance in all the mock communities, with no false positives detected at these higher abundance levels. However, the lower detection limit for these three methods appears to be somewhere between 0.1% and 0.02% relative abundance. An important caveat is that these detection limits are based on results from the PacBio HiFi staggered communities, which consist of 2–2.5 million reads and a minimum detection count of 20–25 reads (Table 3).

Finally, it is important to consider the impact of novel sequences on performance. All species in our study have suitable representation in the databases used (but see caveats for Zymo D6331), and we therefore did not investigate this topic explicitly. However, we propose three features may be important for working with novel diversity in empirical samples. First, the LCA algorithm provides beneficial behavior in ambiguous cases, preventing mis-assignments at the species level by making assignments to higher taxa. Second, protein-based alignments may be more advantageous than nucleotide alignments or k-mer matches for highly distant sequences. Finally, methods which utilize large, comprehensive databases should provide advantages over smaller or marker-specific databases. For example, utilizing NCBI nt or nr allows for the inclusion of new sequences that are continuously deposited in public databases. We propose the effects of novel sequences would be a useful topic for future study, particularly for long-read datasets.

### Effects of Shorter Reads

Our comparisons of length-filtered datasets strongly suggest that including shorter long reads (< 2kb) can have an adverse effect on taxonomic profiling. We found that datasets with many shorter reads had significantly lower precision and F-scores compared to datasets containing only longer reads. We also found that the inclusion of shorter reads heavily skewed relative abundance estimates, which are based on read counts in our experiment. We acknowledge that calculating abundance estimates from the total number of aligned bases could potentially mitigate this effect. More importantly, we found that precision, F-scores, and relative abundances were affected across all methods, suggesting these shorter read lengths may be a “gray” zone for both classes of methods. For example, some long-read methods require the presence of multiple genes for the LCA algorithm to function well (MMSeqs2, MEGAN-LR-prot). Reads that are <2kb are unlikely to satisfy this criterion. Therefore, we strongly recommend filtering these shorter long reads before attempting taxonomic classification. This can be achieved bioinformatically after sequencing, but performing size selection during library preparation can also greatly reduce the number of shorter fragments that are sequenced.

### Effects of Read Accuracy

We included mock community datasets sequenced with PacBio HiFi and ONT, allowing for limited comparisons of methods across sequencing technologies. One noticeable difference occurs in read utilization for methods that perform translation alignments to protein references and exact k-mer matching. For example, more reads were assigned in HiFi versus ONT datasets for MMseqs2 (94–99% vs. 46–67%) and to a lesser extent MEGAN-LR­prot (71–74% vs. 60–62%). This result could be related to differences in the mock communities sequenced, however the species in all three mock communities are expected to have adequate representation in the databases (except two species in HiFi Zymo D6331). It is more likely that differences in error profiles explain these results, as even slightly higher error rates are expected to negatively impact translation alignment (broken reading frames, premature stop codons). This is idea is supported by two observations. First, this effect was more pronounced for MMseqs2, which uses Prodigal for translation rather than a frameshift-aware method such as DIAMOND. Second, the ONT data include an R10.3 dataset with Guppy basecalling (mean = Q10.5; reported at data source) and the newest “Q20” chemistry release with Bonito v0.3.5 basecalling (expected modal quality ∼Q20), and we found fewer reads were assigned in the R10.3 dataset versus the Q20 dataset for MMSeqs2 (46% vs. 67%, respectively). We note the same pattern was present for Centrifuge-500, which requires 500 matched k-mer bases to the reference; read assignment improved dramatically from ONT R10.3 to Q20 (1% vs. 53%, respectively). This result also occurred for sourmash, another k-mer-based method. Here, read assignment improved from ONT R10.3 to Q20 (41% vs. 68% for sourmash-k31; 26% vs. 59% for sourmash-k51). However, despite the improvement in accuracy for the ONT Q20 dataset, it still had lower read assignment for protein alignment methods and sourmash as compared to both HiFi datasets (Fig. 2). The HiFi ATCC and Zymo datasets are more accurate; all reads are >Q20 and the median scores are Q36 and Q40. Together, these results suggest that read quality remains critical for high-quality taxonomic profiling with long-read methods.

Different mock communities were available for PacBio HiFi (ATCC MSA-1003, Zymo D6331) and ONT (Zymo D6300), which prevents a direct comparison of detection metrics (precision, recall, and F-scores) and detection limits across sequencing technologies. The mock community sequenced with ONT is simpler than the HiFi mock communities in terms of the total number of species (10 vs. 17/20) and relative abundances (even vs. staggered). The simpler mock community design also prevented us from estimating recall and detection limits for lower abundance species with ONT data; our conclusions about detection power at low abundances are based exclusively on PacBio HiFi data. In their study, Marić et al. [17] found that ONT pseudo-mock datasets displayed lower classification accuracy, higher false positives, and higher relative abundance error relative to PacBio pseudo-mock datasets. However, the pseudo-mock datasets for ONT and PacBio included in their study contained different numbers of species and abundance designs, meaning they were not direct comparisons. We caution against this type of approach, and instead propose that an objective comparison of detection metrics should be performed by sequencing the same mock community standard using both technologies. We also propose that a mock standard with high species diversity and staggered abundances will provide the most meaningful information for future benchmarking studies.

## Conclusion

With increasing quality and prevalence of long-read datasets, it is critical to assess the utility of these data for taxonomic profiling of metagenomic samples. Here, we evaluated several profiling and classification methods for mock communities sequenced with PacBio HiFi and ONT. We also included Illumina short read data for these communities as a comparison. Our results demonstrate there are clear precision and recall trade-offs associated with each method. We found that several popular short-read methods (Kraken2, Bracken, Centrifuge) resulted in many false positives, particularly at lower abundance levels. Filtering can increase precision for these methods, but comes at the cost of severely reducing recall. Importantly, we determined this pattern of low precision and high recall occurred for these methods using both long-read and short-read datasets. This suggests the methods themselves, rather than differences in read lengths or platform, are driving these outcomes. By contrast, we found sourmash and several long-read classifiers displayed high precision and recall without any filtering necessary. These long-read classifiers are alignment-based, and include BugSeq (nucleotide alignments), and MEGAN-LR using translation alignments (DIAMOND to NCBI nr) or nucleotide alignments (minimap2 to NCBI nt). Sourmash has the highest detection power, finding all species down to 0.001% relative abundance with minimal false positives. Our comparisons between long-read sequencing technologies indicate that read quality remains critical for taxonomic profiling performance. We found that read accuracy impacts the success of methods relying on protein predictions or exact k-mer matches. We also found a high proportion of shorter long reads (<2kb) can result in lower precision and inaccurate abundance estimates, relative to length-filtered datasets. However, we emphasize that for any given mock community, the long-read dataset (analyzed with sourmash or any long-read method) produced significantly better results than the short-read datasets. Methods which utilize long-range information present in long-read datasets provide clear improvements in taxonomic profiling and abundance estimation, and demonstrate a clear advantage over short-read methods. To continue studying these effects, we propose that cross-platform sequencing of more complex standardized mock communities would be useful for future benchmarking studies.

## Declarations

### Ethics approval and consent to participate

Not applicable.

### Consent for Publication

Not applicable.

### Availability of Data and Materials

The mock community datasets are publicly available from the National Center for Biotechnology Information (NCBI), European Nucleotide Archive (ENA), or public lab websites: HiFi ATCC MSA-1003 (NCBI: PRJNA546278: SRX6095783), HiFi Zymo D6331 (NCBI: PRJNA680590: SRX9569057), Illumina ATCC MSA-1003 (NCBI: PRJNA510527: SRX5169925), Illumina Zymo D6300 (NCBI: PRJNA648136: SRX8824472), ONT Q20 Zymo D6300 (ENA: PRJEB43406: ERR5396170), and ONT R10 Zymo D6300 (https://lomanlab.github.io/mockcommunity/r10.html). The kreport output files for all methods and datasets, along with Jupyter notebooks and results files, are freely available on the Open Science Framework: https://osf.io/bqtdu/.

## Funding

Not applicable.

### Authors’ Contributions

Daniel M. Portik, N. Tessa Pierce-Ward and C. Titus Brown conceptualized the experiment, Daniel M. Portik and N. Tessa Pierce-Ward performed data analysis, Daniel M. Portik and N. Tessa Pierce-Ward wrote the manuscript, and all authors reviewed the manuscript.

## Acknowledgments

We thank R. Hall, W. Rowell, and K.P. Chua for helpful feedback on an early version of this manuscript.

## LITERATURE CITED

1. Breitwieser, F.P., Lu, J., and S.L. Salzberg. (2019). A review of methods and databases for metagenomic classification and assembly. Briefings in Bioinformatics, 20: 1125–1139.

2. Lindgreen, S., Adair, K.L., and P.P. Gardner. (2016) An evaluation of the accuracy and speed of metagenome analysis tools. Scientific Reports, 6: 19233.

3. McIntyre, A.B.R., Ounit, R., Afshinnekoo, E., Prill, R.J., Hénaff, E., Alexander, N., Minot, S.S., Danko, D., Foox, J., Ahsanuddin, S., Tighe, S., Hasan, N.A., Subramanian, P., Moffat, K., Levy, S., Lonardi, S., Greenfield, N., Colwell, R.R., Rosen, G.L., and C.E. Mason. (2017). Comprehensive benchmarking and ensemble approaches for metagenomic classifiers. Genome Biology, 18: 182.

4. Sczyrba, A., Hofmann, P., Belmann, P., Koslicki, D., Janssen, S., Dröge, J., Gregor, I., Majda, S., Fiedler, J., Dahms, E., Bremges, A., Fritz, A., Garrido-Oter, R., Jørgensen, T.S.S., Shapiro, N., Blood, P.D., Gurevich, A., Bai, Y., Turaev, D., DeMaere, M.Z., Chikhi, R., Nagarajan, N., Quince, C., Meyer, F., Balvočiūtė, M., Hansen, L.H.H., Sørensen, S.J., Chia, B.K.H., Denis, B., Froula, J.L., Wang, Z., Egan, R., Don Kang, D., Cook, J.J., Deltel, C., Beckstette, M., Lemaitre, C., Peterlongo, P., Rizk, G., Lavenier, D., Wu, Y.-W.W., Singer, S.W., Jain, C., Strous, M., Klingenberg, H., Meinicke, P., Barton, M.D., Lingner, T., Lin, H.­H.H., Liao, Y.-C.C., Silva, G.G.G.Z., Cuevas, D.A., Edwards, R.A., Saha, S., Piro, V.C., Renard, B.Y., Pop, M., Klenk, H.-P.P., Göker, M., Kyrpides, N.C., Woyke, T., Vorholt, J.A., Schulze-Lefert, P., Rubin, E.M., Darling, A.E., Rattei, T., and A.C. McHardy. (2017) Critical assessment of metagenome interpretation -a benchmark of metagenomics software. Nature Methods, 14: 1063–1071.

5. Escobar-Zepeda, A., Godoy-Lozano, E.E., Raggi, L., Segovia, L., Merino, E., Gutiérrez-Rios, R.M., Juarez, K., Licea-Navarro, A.F., Pardo-Lopez, L., and A. Sanchez-Flores. (2018). Analysis of sequencing strategies and tools for taxonomic annotation: defining standards for progressive metagenomics. Scientific Reports, 8: 12034.

6. Meyer, F., Bremges, A., Belmann, P., Janssen, S., McHardy, A.C., and D. Koslicki. (2019). Assessing taxonomic metagenome profiles with OPAL. Genome Biology, 20: 51.

7. Tamames, J., Cobo-Simón, M., and F. Puente-Sánchez. (2019). Assessing the performance of different approaches for functional and taxonomic annotation of metagenomes. BMC Genomics, 20: 960.

8. Ye, S.H., Siddle, K.J., Park, D.J., and P.C. Sabeti. (2019). Benchmarking metagenomics tools for taxonomic classification. Cell. 178: 779–794.

9. Parks, D.H., Rigato, F., Vera-Wolf, P., Krause, L., Hugenholtz, P., Tyson, G.W., and D.L.A. Wood. (2021). Evaluation of the microba community profiler for taxonomic profiling of metagenomic datasets from the human gut microbiome. Frontiers in Microbiology, 12: 643682.

10. Meyer, F., Fritz, A., Deng, Z.-L., Koslicki, D., Lesker, T.L., Gurevich, A., Robertson, G., Alser, M., Antipov, D., Beghini, F., Bertrand, D., Brito, J.J., Brown, C.T., Buchmann, J., Buluç, A., Chen, B., Chikhi, R., Clausen, P.T.L.C., Cristian, A., Dabrowski, P.W., Darling, A.E., Egan, R., Eskin, E., Georganas, E., Goltsman, E., Gray, M.A., Hansen, L.H., Hofmeyr, S., Huang, P., Irber, L., Jia, H., Jørgensen, T.S., Kieser, S.D., Klemetsen, T., Kola, A., Kolmogorov, M., Korobeynikov, A., Kwan, J., LaPierre, N., Lemaitre, C., Li, C., Limasset, A., Malcher-Miranda, F., Mangul, S., Marcelino, V.R., Marchet, C., Marijon, P., Meleshko, D., Mende, D.R., Milanese, A., Nagarajan, N., Nissen, J., Nurk, S., Oliker, L., Paoli, L., Peterlongo, P., Piro, V.C., Porter, J.S., Rasmussen, S., Rees, E.R., Reinert, K., Renard, B., Robertsen, E.M., Rosen, G.L., Ruscheweyh, H.-J., Sarwal, V., Segata, N., Seiler, E., Shi, L., Sun, F., Sunagawa, S., Sørensen, S.J., Thomas, A., Tong, C., Trajkovski, M., Tremblay, J., Uritskiy, G., Vicedomini, R., Wang, Z., Wang, Z., Wang, Z., Warren, A., Willassen, N.P., Yelick, K., You, R., Zeller, G., Zhao, Z., Zhu, S., Zhu, J., Garrido-Oter, R., Gastmeier, P., Hacquard, S., Häußler, S., Khaledi, S., Maechler, F., Mesny, F., Radutoiu, S., Schulze-Lefert, P., Smit, N., Strowig, T., Bremges, A., Sczyrba, A., and A.C. McHardy. (2022). Critical assessment of metagenome interpretation: the second round of challenges. Nature Methods, 19: 420–440.

11. Wenger, A.M., Peluso, P., Rowell, W.J., Chang, P.-C., Hall, R.J., Concepcion, G.T., Ebler, J., Fungtammasan, A., Kolesnikov, A., Olson, N.D., Töpfer, A., Alonge, M., Mahmoud, M., Qian, Y., Chin, C.-S., Phillippy, A.M., Schatz, M.C., Myers, G., DePristo, M.A., Ruan, J., Marschall, T., Sedlazeck, F.J., Zook, J.M., Li, H., Koren, S., Carroll, A., Rank, D.A., and M.W. Hunkapiller. (2019). Accurate circular consensus long-read sequencing improves variant detection and assembly of a human genome. Nature Biotechnology, 37: 1155–1162.

12. Dilthey, A.T., Jain, C., Koren, S., and A.M. Phillippy. (2019). Strain-level metagenomic assignment and compositional estimation for long reads with MetaMaps. Nature Communications, 10: 3066.

13. Huson, D.H., Beier, S., Flade, I., Górska, A., El-Hadidi, M., Mitra, S., Ruscheweyh, H.-J., and R. Tappu. (2016). MEGAN Community Edition – interactive exploration and analysis of large-scale microbiome sequencing data. PLOS Computational Biology, 12: e1004957.

14. Mirdita, M., Steinegger, M., Breitwieser, F., Söding, J., and E.L. Karin. (2021). Fast and sensitive taxonomic assignment to metagenomic contigs. Bioinformatics, 2021: 1–3.

15. Fan, J., Huang, S., and S.D. Chorlton. (2021). BugSeq: a highly accurate cloud platform for long-read metagenomic analyses. BMC Bioinformatics, 22: 160.

16. Leidenfrost, R.M., Pöther, D.-C., Jäckel, U., and R. Wünschiers. (2020). Benchmarking the MinION: evaluating long reads for microbial profiling. Scientific Reports, 10: 5125.

17. Pearman, W.S., Freed, N.E., and O.K. Silander. (2020). Testing the advantage and disadvantages of short- and long-read eukaryotic metagenomics using simulated reads. BMC Bioinformatics, 21: 220.

18. Marić, J., Križanović, K., Riondet, S., Nagarajan, N., and M. Šikić. (2020). Benchmarking metagenomic classification tools for long-read sequencing data. bioRxiv, https://doi.org/10.1101/2020.11.25.397729.

19. Govender, K.N., and D.W. Eyre. (2022). Benchmarking taxonomic classifiers with Illumina and Nanopore sequence data for clinical metagenomic diagnostic applications. Microbial Genomics, 8: 000886.

20. Nicholls, S.M., Quick, J.C., Tang, S., and N.J. Loman. (2019). Ultra-deep, long-read nanopore sequencing of mock microbial community standards. GigaScience, 8: 1–9.

21. De Coster, W., D’Hert, S., Schultz, D.T., Cruts, M., and C. Van Broeckhoven. (2018). NanoPack: visualizing and processing long-read sequencing data. Bioinformatics, 34: 2666– 2669.

22. Sui, H.-Y., Weil, A.A., Nuwagira, E., Qadri, F., Ryan, E.T., Mezzari, M.P., Phipatanakul, W., and P.S. Lai. (2020). Impact of DNA extraction method on variation in human and built environment microbial community and functional profiles assessed by shotgun metagenomics sequencing. Frontiers in Microbiology, 11: 953.

23. Wood, D.E., and S.L. Salzberg. (2014). Kraken: ultrafast metagenomic sequence classification using exact alignments. Genome Biology, 15: R46.

24. Wood, D.E., Lu, J., and B. Langmead. (2019). Improved metagenomic analysis with Kraken 2. Genome Biology, 20: 257.

25. Lu, J., Breitwieser, F.P., Thielen, P., and S.L. Salzberg. (2017). Bracken: estimating species abundance in metagenomics data. PeerJ Computer Science, 3: e104.

26. Kim, D., Song, L., Breitwieser, F.P., and S.L. Salzberg. (2016). Centrifuge: rapid and sensitive classification of metagenomic sequences. Genome Research, 26: 1721–1729.

27. Beghini, F., McIver, L.J., Blanco-Miguez, A., Dubois, L., Asnicar, F., Maharjan, S., Mailyan, A., Thomas, A.M., Manghi, P., Valles-Colomer, M., Weingart, G., Zhang, Y., Zolfo, M., Huttenhower, C., Franzosa, E.A., and N. Segata. (2021). Integrating taxonomic, functional, and strain-level profiling of diverse microbial communities with bioBakery 3. eLife, 10:e65088.

28. Milanese, A., Mende, D.R., Paoli, L., Salazar, G., Ruscheweyh, H.-J., Cuenca, M., Hingamp, P., Alves, R., Costea, P.I., Coelho, L.P., Schmidt, T.S.B., Almeida, A., Mitchell, A.L., Finn, R.D., Huerta-Cepas, J., Bork, P., Zeller, G., and S. Sunagawa. (2019). Microbial abundance, activity and population genomic profiling with mOTUs2. Nature Communications, 10: 1014.

29. Huson, D.H., Albrecht, B., Bağci, C., Bessarab, I., Górska, A., Jolic, D., and R.B.H. Williams. (2018). MEGAN-LR: new algorithms allow accurate binning and easy interactive exploration of metagenomic long reads and contigs. Biology Direct, 13: 6.

30. Buchfink, B., Xie, C., and D.H. Huson. (2015) Fast and sensitive protein alignment using DIAMOND. Nature Methods, 12: 59–60.

31. Arumugam, K., Bağci, C., Bessarab, I., Beier, S., Buchfink, B., Górska, A., Qiu, G., Huson, D.H., and R.B.H. Williams. (2019). Annotated bacterial chromosomes from frame-shift­corrected long-read metagenomic data. Microbiome, 7: 61.

32. Li, H. (2018). Minimap2: pairwise alignment for nucleotide sequences. Bioinformatics, 34: 3094–3100.

33. Brown, C.T., and L. Irber. (2016). sourmash: a library for MinHash sketching of DNA. Journal of Open Source Software, 1: 27.

34. Pierce, N.T., Irber, L., Reiter, T., Brooks, P., and C.T. Brown. (2019). Large-scale sequence comparisons with *sourmash*. F1000Research, 8: 1006.

35. Irber, L., Brooks, P.T., Reiter, T., Pierce-Ward, N.T., Hera, M.R., Koslicki, D., and C.T. Brown. (2022). Lightweight compositional analysis of metagenomes with FracMinHash and minimum metagenome covers. bioRxiv, https://doi.org/10.1101/2022.01.11.475838

36. Koslicki, D., and D. Falush. (2016). MetaPalette: a k-mer painting approach for metagenomic profiling and quantification of novel strain variation. mSystems, 1: e00020–16.

