## Supplementary Material for "Evaluation of taxonomic classification and profiling methods for long-read shotgun metagenomic sequencing datasets"

**for**

**Table of Contents**

| **Item** | **Description** | **Page** |
| --- | --- | --- |
| Figure S1 | Read length distributions for secondary ONT datasets. | 3 |
| Figure S2 | Average precision, recall, and F1 scores for HiFi datasets. | 4 |
| Figure S3 | Average precision, recall, and F1 scores for ONT datasets. | 5 |
| Figure S4 | Precision, recall and F-scores, 0.1% threshold, species-level analysis. | 6 |
| Figure S5 | Precision, recall and F-scores, 0.1% threshold, genus-level analysis. | 7 |
| Figure S6 | Precision, recall and F-scores, 1% threshold, species-level analysis. | 8 |
| Figure S7 | Precision, recall and F-scores, 1% threshold, genus-level analysis. | 9 |
| Figure S8 | Precision, recall and F-scores for the ONT Short datasets. | 10 |
| Figure S9 | Species and genus-level relative abundances for the ONT Short datasets. | 11 |
| Figure S10 | Precision, recall and F-scores for short-read datasets. | 12 |
| Figure S11 | Species-level relative abundances for short-read datasets. | 13 |
| Figure S12 | Genus-level relative abundances for short-read datasets. | 14 |
| Table S1 | Species-level read counts for HiFi ATCC MSA-1003. | 15 |
| Table S2 | Genus-level read counts for HiFi ATCC MSA-1003. | 16 |
| Table S3 | Species-level read counts for HiFi Zymo D6331. | 17 |
| Table S4 | Genus-level read counts for HiFi Zymo D6331. | 18 |
| Table S5 | Species-level read counts for ONT R10 Zymo D6300. | 19 |
| Table S6 | Genus-level read counts for ONT R10 Zymo D6300. | 20 |
| Table S7 | Species-level read counts for ONT Q20 Zymo D6300. | 21 |
| Table S8 | Genus-level read counts for ONT Q20 Zymo D6300. | 22 |
| Table S9 | Abundance level-specific detection for staggered, 0.001% threshold. | 23 |
| Table S10 | Genus-level detection results, 0.001% threshold. | 25 |
| Table S11 | Species-level detection results, 0.1% threshold. | 28 |
| Table S12 | Abundance level-specific detection for staggered, 0.1% threshold. | 31 |
| Table S13 | Genus-level detection results, 0.1% threshold. | 33 |
| Table S14 | Species-level detection results, 1% threshold. | 36 |
| Table S15 | Abundance level-specific detection for staggered, 1% threshold. | 39 |
| Table S16 | Genus-level detection results, 1% threshold. | 41 |
| Table S17 | Chi-squared tests for relative abundances at the species level. | 44 |
| Table S18 | Chi-squared tests for relative abundances at the genus level. | 45 |
| Table S19 | Species-level detection results for ONT Short datasets. | 46 |
| Table S20 | Genus-level detection results for ONT Short datasets. | 49 |
| Table S21 | Chi-Squared tests for species level abundances, ONT Short datasets. | 52 |
| Table S22 | Chi-Squared tests for genus level abundances, ONT Short datasets. | 53 |
| Table S23 | Species-level detection results for ATCC short-read datasets. | 54 |
| Table S24 | Genus-level detection results for ATCC short-read datasets. | 56 |
| Table S25 | Chi-Squared tests for species abundances, ATCC short-read datasets. | 58 |
| Table S26 | Chi-Squared tests for genus abundances, ATCC short-read datasets. | 58 |
| Table S27 | Species-level detection results for Zymo D6300 short-read datasets. | 59 |
| Table S28 | Genus-level detection results for Zymo D6300short-read datasets. | 61 |
| Table S29 | Chi-Squared tests for species abundances, Zymo D6300 short-reads. | 63 |
| Table S30 | Chi-Squared tests for genus abundances, Zymo D6300 short-reads. | 63 |

**Supplementary Figure S1.** Violin plots showing the read length distributions of the primary and secondary versions of the ONT datasets included in this study. Interiors of plots contain white dots representing median values, black bars represent interquartile values, and black lines represent minimum and maximum range values. Read sizes range up to 50,000 bp in length, but the plot is clipped at 14,000 bp to show the core distributions. The ONT R10 Short dataset contains all the reads present in ONT R10 Zymo D6300, plus all reads < 2kb. The ONT Q20 Short dataset was obtained by subsampling 2 million reads that were < 3kb in length.

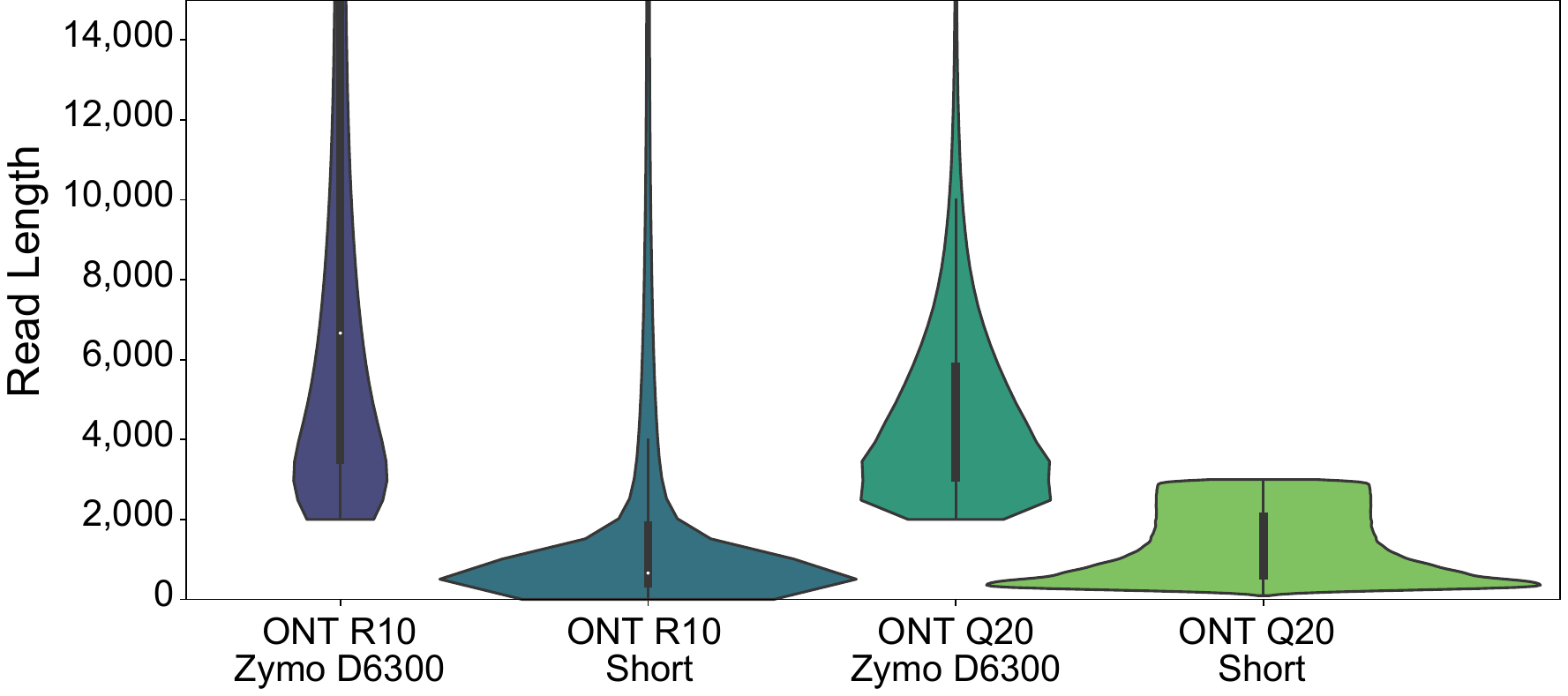

**Supplementary Figure S2.** The average values for HiFi datasets for (A) precision and recall and (B) F1 scores for the species-level analysis based on a minimum threshold of 0.001% of the total reads.

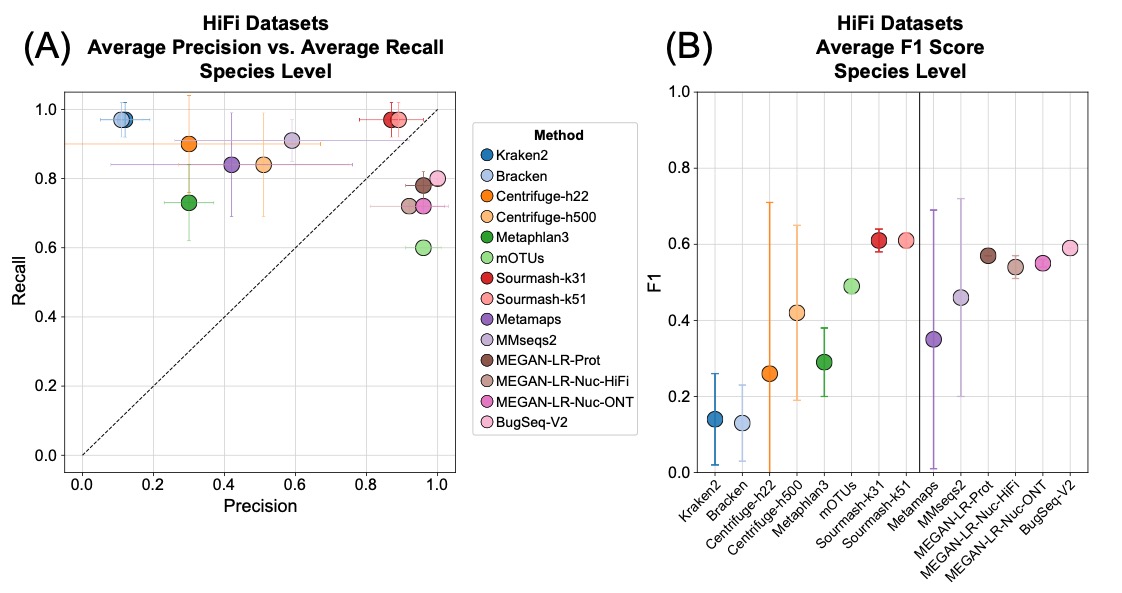

**Supplementary Figure S3.** The average values for ONT datasets for (A) precision and recall and (B) F1 scores for the species-level analysis based on a minimum threshold of 0.001% of the total reads.

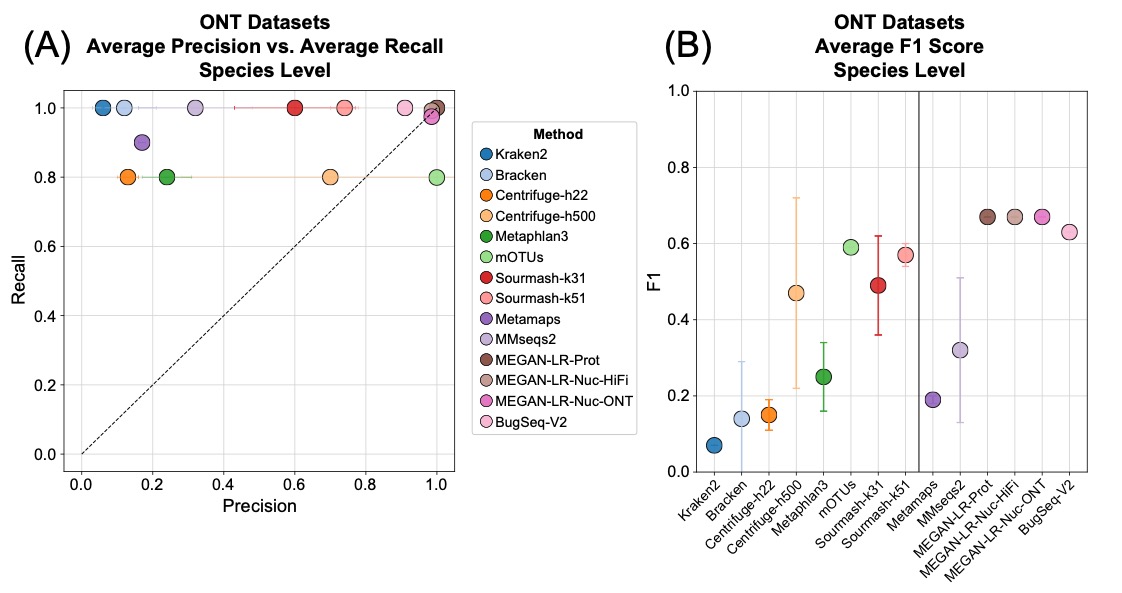

**Supplementary Figure S4.** Precision, recall and F-scores for the species-level analysis based on the minimum 0.1% of total reads threshold.

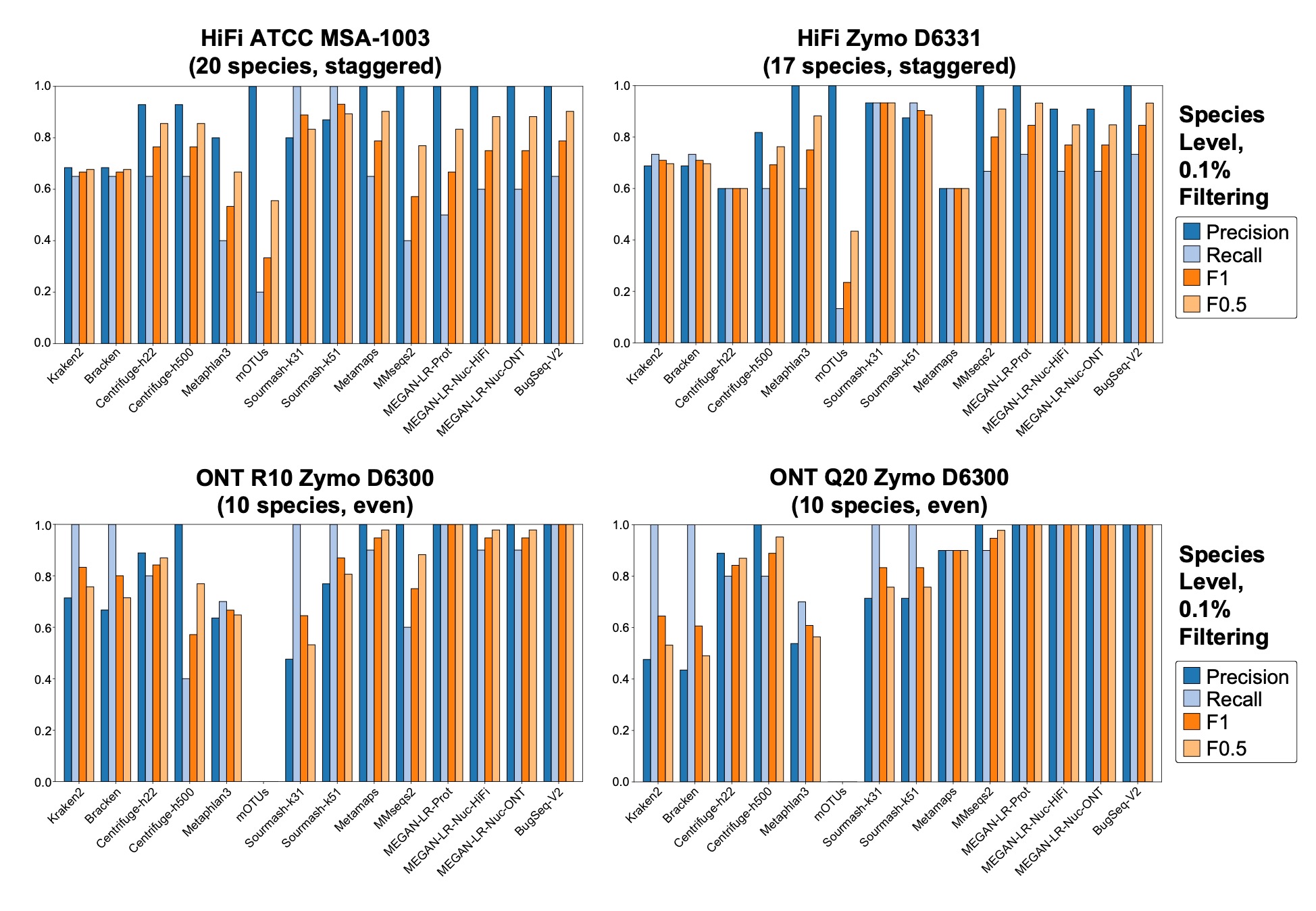

**Supplementary Figure S5.** Precision, recall and F-scores for the genus-level analysis based on the minimum 0.1% of total reads threshold.

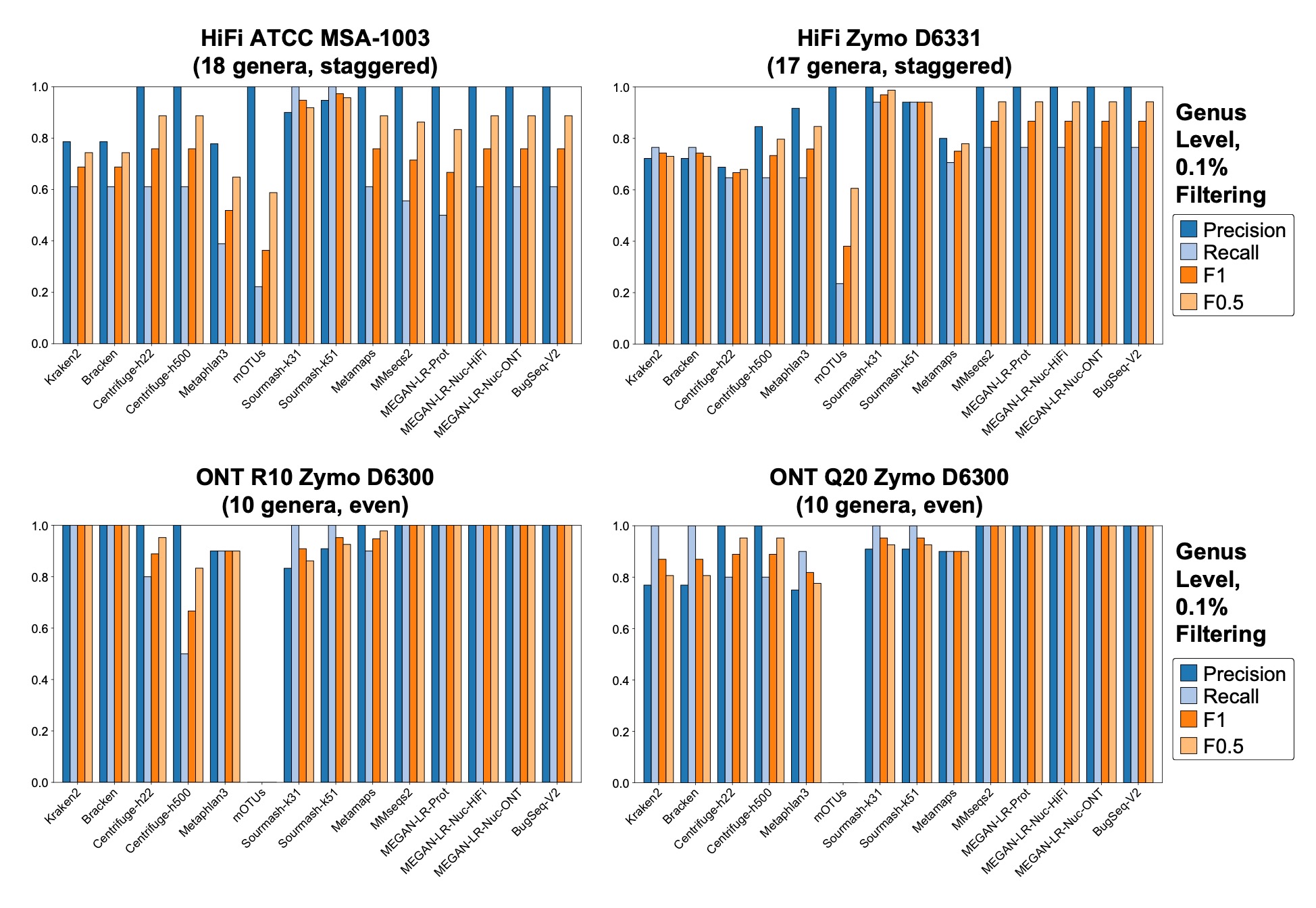

**Supplementary Figure S6.** Precision, recall and F-scores for the species-level analysis based on the minimum 1% of total reads threshold. Empty spaces indicate no species met the minimum required read count based on the minimum threshold.
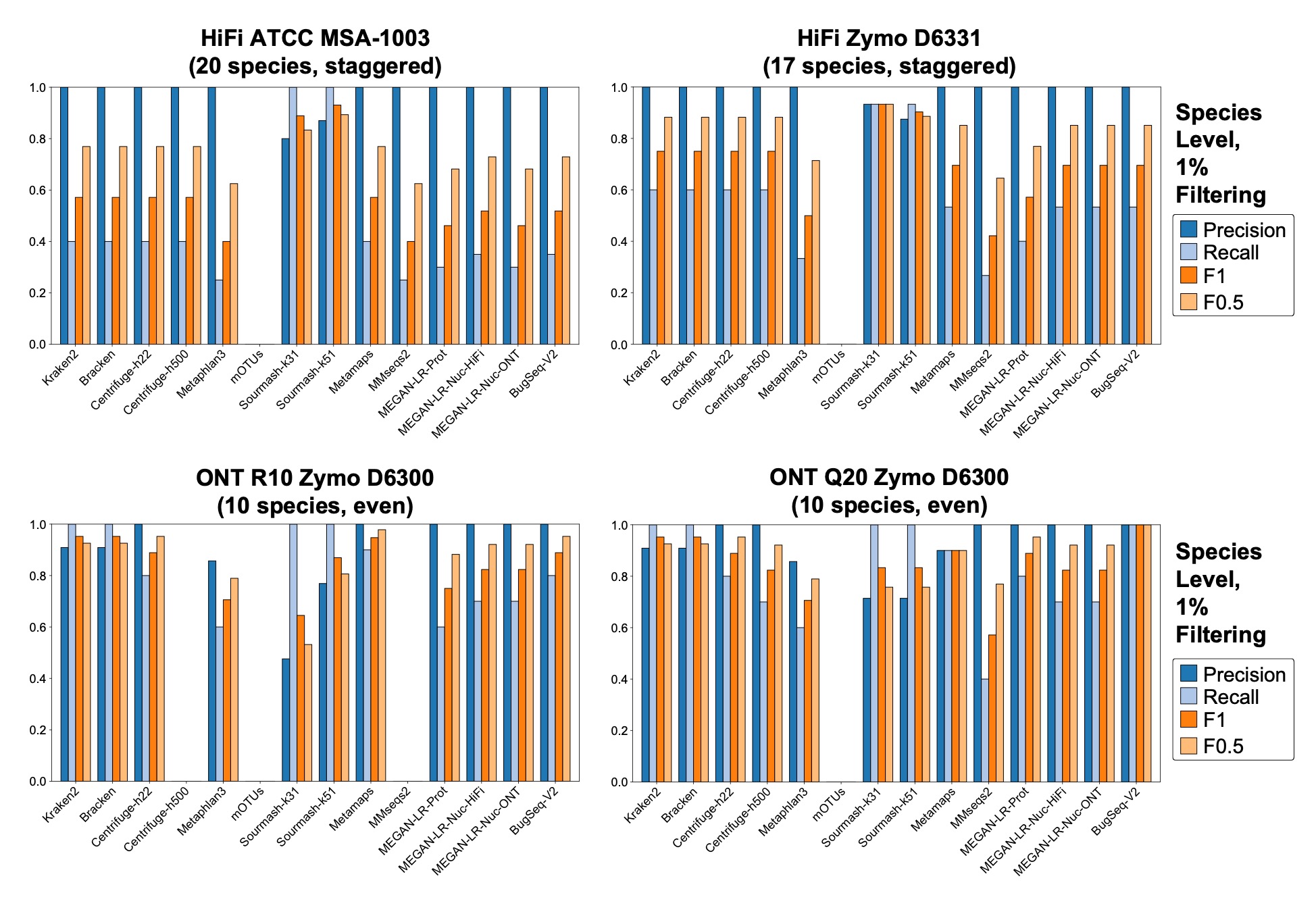

**Supplementary Figure S7.** Precision, recall and F-scores for the genus-level analysis based on the minimum 1% of total reads threshold. Empty spaces indicate no genera met the minimum required read count based on the minimum threshold.

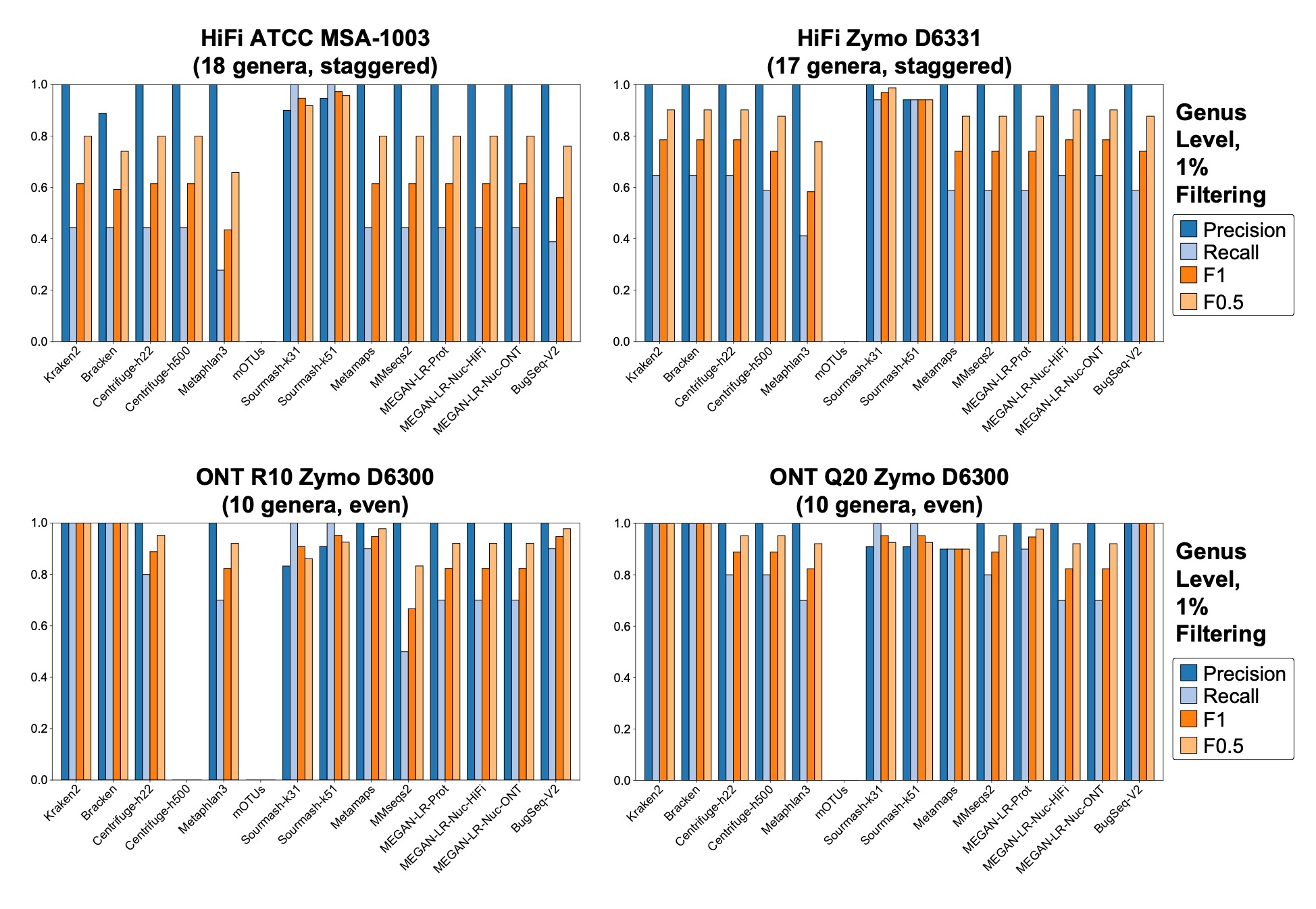

**Supplementary Figure S8.** Precision, recall and F-scores for the species-level analysis of the ONT Short datasets based on all thresholds. Empty spaces indicate no species met the minimum required read count based on the minimum threshold.

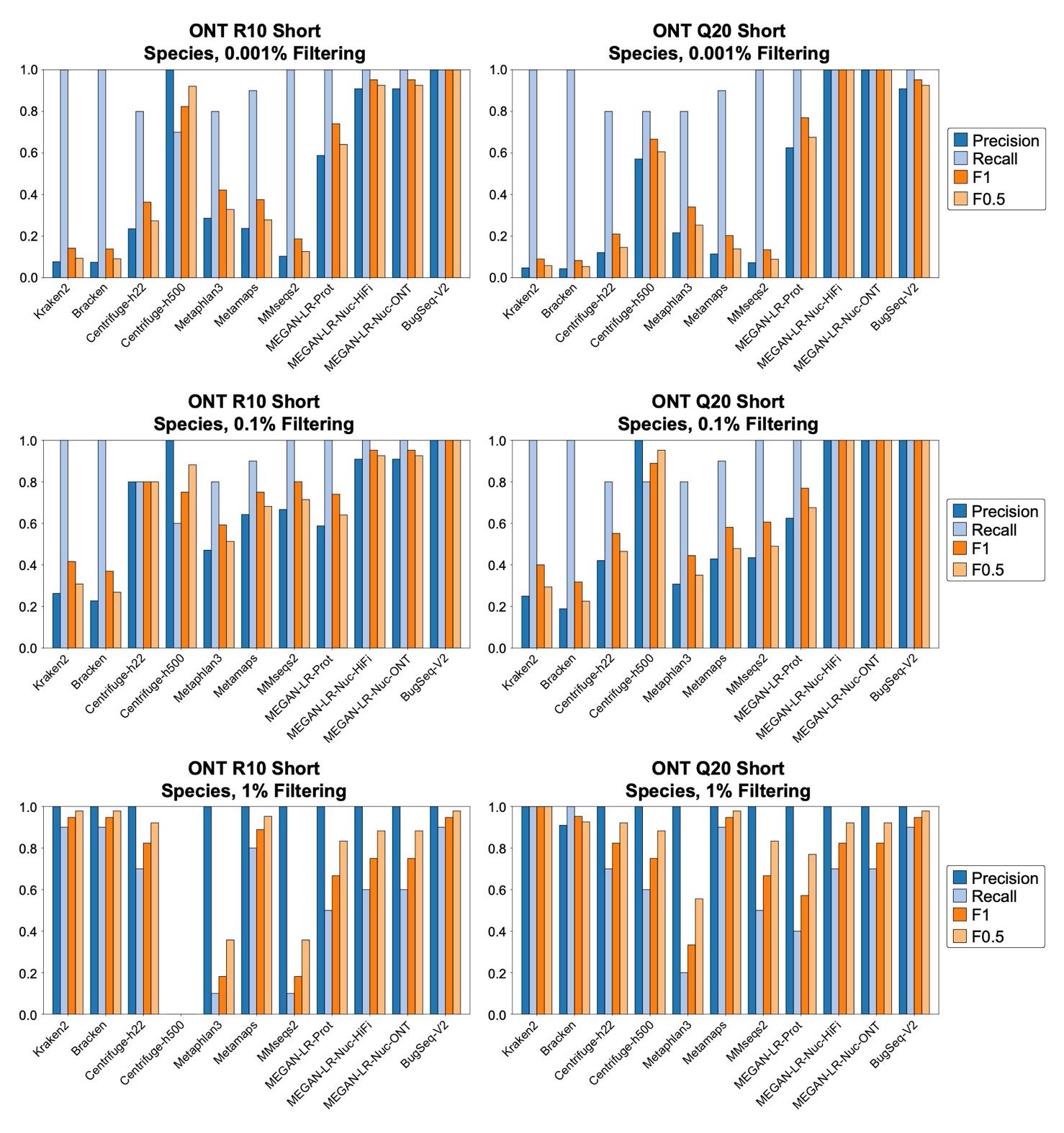

**Supplementary Figure S9.** Species and genus-level relative abundance estimates for the ONT Short datasets. The theoretical distributions are shown on the left and are based on the manufacturer’s specifications. The read counts for false positives were grouped in a category labeled ‘Other’. Asterisks signify methods that failed the chi-squared goodness of fit test (e.g., the abundance estimates were significantly different from the theoretical values).

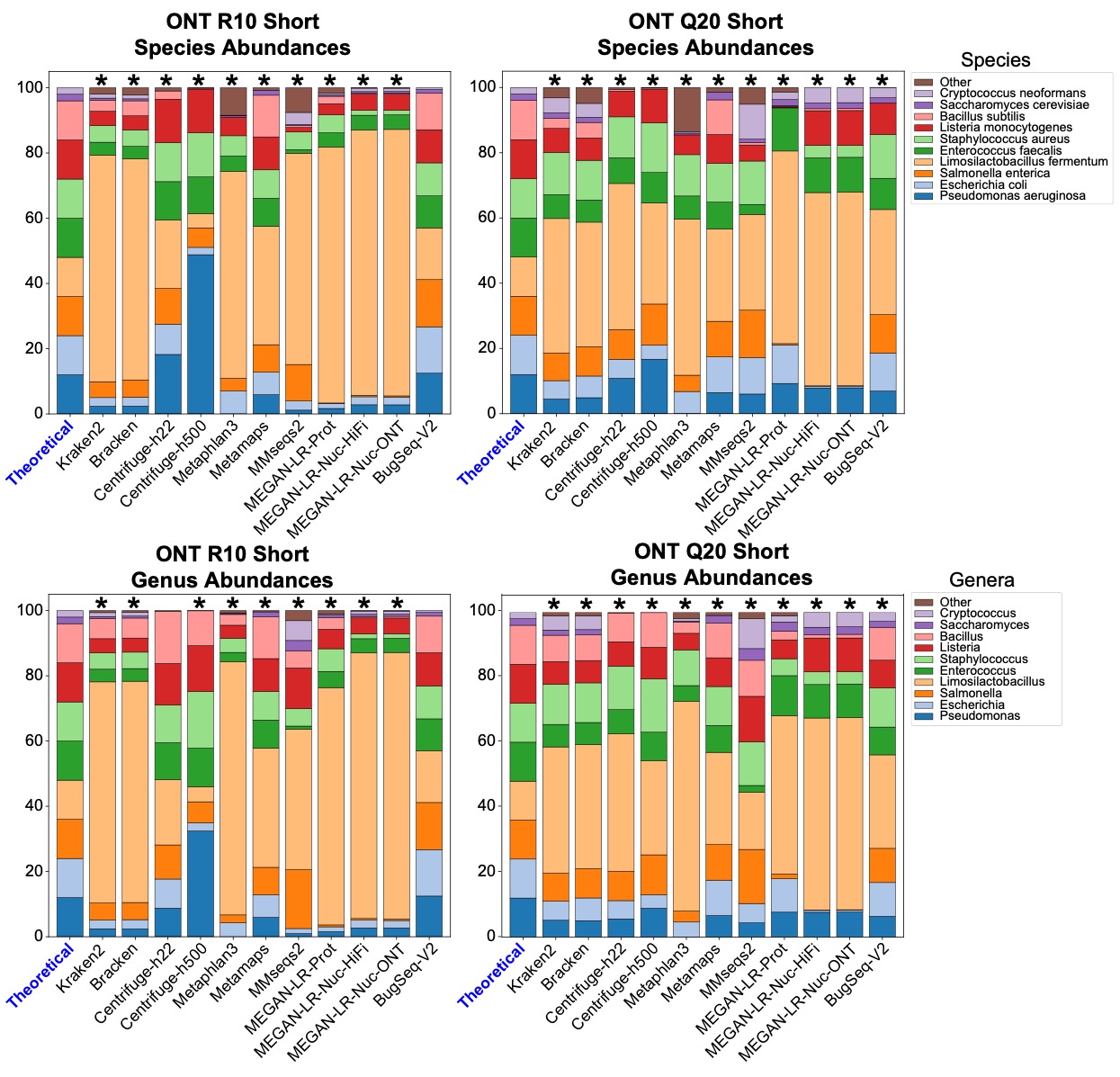

**Supplementary Figure S10.** Precision, recall and F-scores for the species-level analysis of the ATCC and Zymo D6300 short-read datasets based on all thresholds.

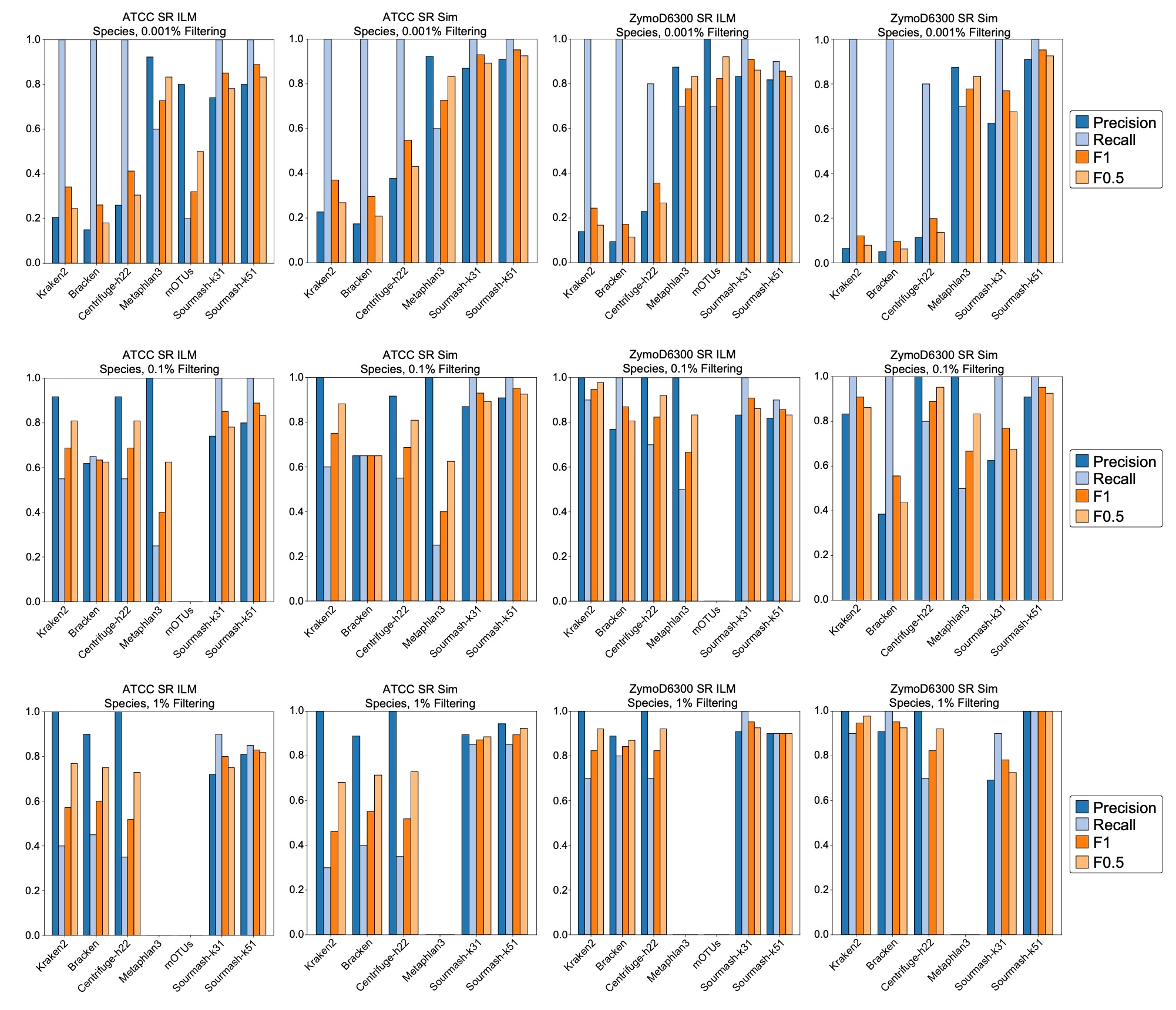

**Supplementary Figure S11.** Species-level relative abundance estimates for the ATCC and Zymo D6300 short-read datasets. The theoretical distributions are shown on the left and are based on the manufacturer’s specifications. The read counts for false positives were grouped in a category labeled ‘Other’. Asterisks signify methods that failed the chi-squared goodness of fit test (e.g., the abundance estimates were significantly different from the theoretical values).

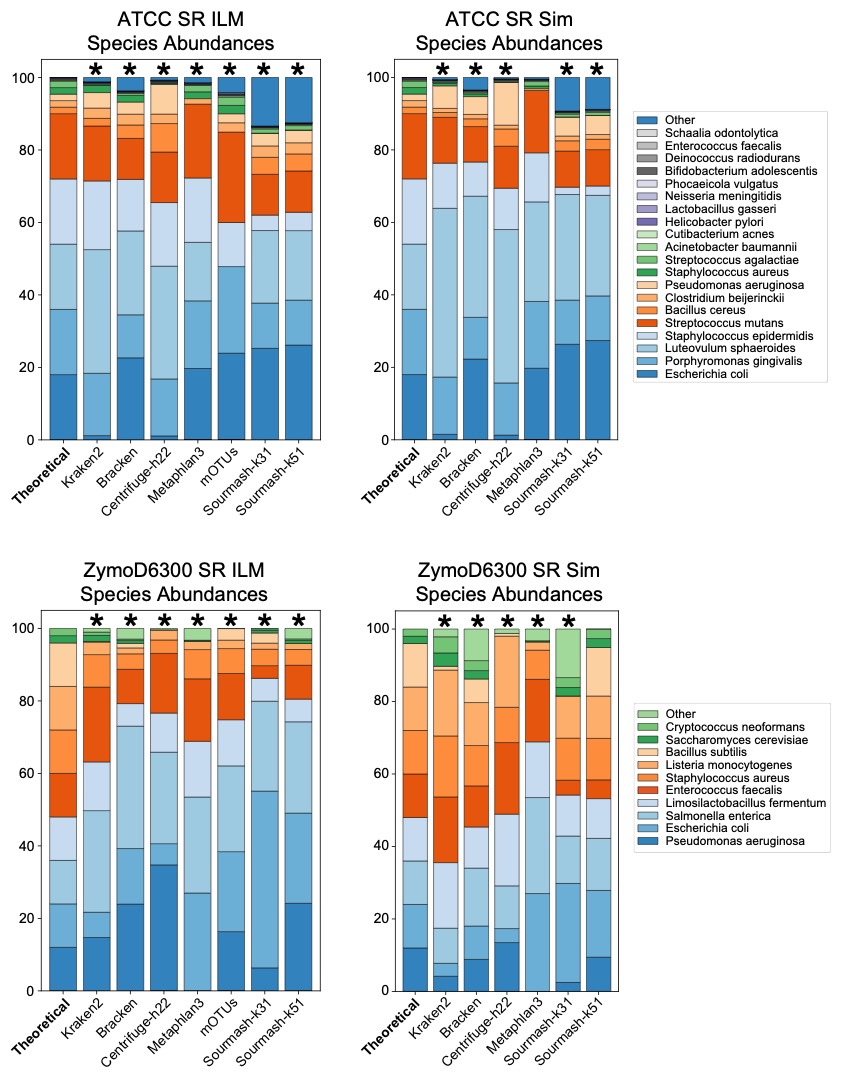

**Supplementary Figure S12.** Genus-level relative abundance estimates for the ATCC and Zymo D6300 short-read datasets. The theoretical distributions are shown on the left and are based on the manufacturer’s specifications. The read counts for false positives were grouped in a category labeled ‘Other’. Asterisks signify methods that failed the chi-squared goodness of fit test (e.g., the abundance estimates were significantly different from the theoretical values).

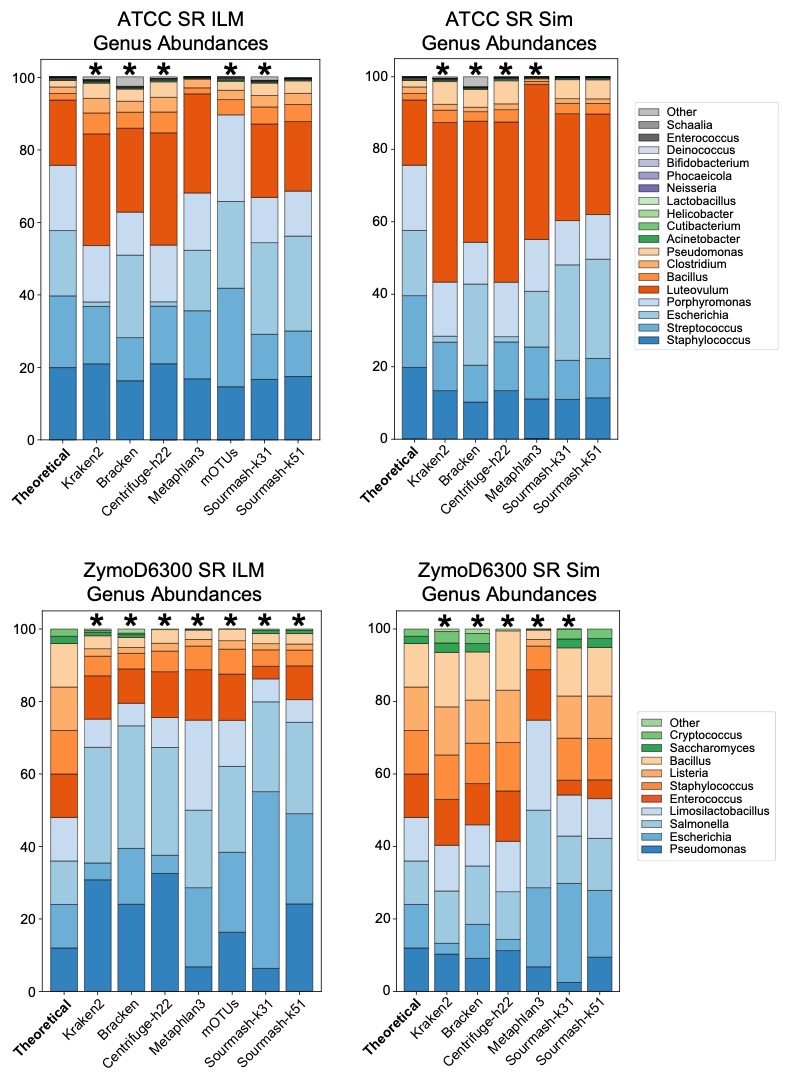

**Supplementary Table S1.** Species-level read counts for HiFi ATCC MSA-1003.

| **Species** | **Theoretical** | **Kraken** | **Bracken** | **Centrifuge-h22** | **Centrifuge-h500** | **Metaphlan3** | **mOTUs** | **Sourmash-k31** | **Sourmash-k51** | **Metamaps** | **MMseqs2** | **MEGAN-Prot** | **MEGAN-Nuc-HiFi** | **MEGAN-Nuc-ONT** | **BugSeq-V2** |
| --- | --- | --- | --- | --- | --- | --- | --- | --- | --- | --- | --- | --- | --- | --- | --- |
| Escherichia coli | 18 | 467,301 | 552,592 | 475,720 | 415,160 | 317,483 | 5,285 | 4930014000 | 4879628000 | 609,822 | 143,757 | 92,628 | 89,722 | 87,293 | 537,965 |
| Porphyromonas gingivalis | 18 | 278,214 | 278,214 | 278,243 | 273,542 | 163,377 | 6,116 | 2199344000 | 2115307000 | 278,196 | 208,314 | 129,938 | 277,930 | 277,858 | 268,073 |
| Luteovulum sphaeroides | 18 | 806,633 | 806,654 | 806,755 | 788,258 | 47,278 | 0 | 5550780000 | 4979545000 | 806,631 | 184,963 | 169,849 | 806,030 | 805,639 | 519,838 |
| Staphylococcus epidermidis | 18 | 231,055 | 231,085 | 232,273 | 226,276 | 191,262 | 3,446 | 347065000 | 423458000 | 232,132 | 8,414 | 68,544 | 226,886 | 225,997 | 214,128 |
| Streptococcus mutans | 18 | 227,952 | 229,905 | 227,935 | 223,088 | 148,270 | 5,078 | 1845396000 | 1761582000 | 227,922 | 104,066 | 86,028 | 226,144 | 220,518 | 227,525 |
| Bacillus cereus | 1.8 | 61,777 | 62,138 | 125,367 | 121,739 | 0 | 0 | 511849000 | 496789000 | 63,155 | 785 | 3,202 | 25,508 | 22,396 | 61,071 |
| Clostridium beijerinckii | 1.8 | 29,443 | 29,566 | 29,366 | 28,012 | 3,690 | 172 | 147740000 | 137691000 | 29,632 | 15,675 | 17,282 | 15,009 | 13,590 | 17,049 |
| Pseudomonas aeruginosa | 1.8 | 116,670 | 116,734 | 234,540 | 229,324 | 0 | 964 | 932163000 | 895719000 | 115,283 | 28,330 | 61,473 | 110,475 | 108,018 | 114,841 |
| Staphylococcus aureus | 1.8 | 17,537 | 17,623 | 14,594 | 13,053 | 6,075 | 172 | 1237000 | 402000 | 16,380 | 8,379 | 11,022 | 14,587 | 14,592 | 10,311 |
| Streptococcus agalactiae | 1.8 | 11,998 | 12,740 | 10,526 | 7,337 | 11,538 | 323 | 130652000 | 128204000 | 17,648 | 880 | 8,266 | 0 | 0 | 15,032 |
| Acinetobacter baumannii | 0.18 | 5,772 | 5,785 | 5,738 | 5,605 | 0 | 67 | 52489999 | 50699999 | 5,840 | 1,612 | 1,526 | 2,664 | 2,427 | 5,778 |
| Cutibacterium acnes | 0.18 | 4,796 | 4,797 | 4,796 | 4,711 | 1,905 | 87 | 39940999 | 38878000 | 4,796 | 666 | 790 | 4,715 | 4,705 | 4,705 |
| Helicobacter pylori | 0.18 | 1,661 | 1,661 | 1,661 | 1,592 | 495 | 53 | 14347000 | 13361000 | 1,661 | 1,653 | 1,356 | 1,660 | 1,658 | 1,768 |
| Lactobacillus gasseri | 0.18 | 1,428 | 1,428 | 1,428 | 1,390 | 492 | 16 | 10059000 | 10635000 | 1,435 | 975 | 470 | 1,350 | 1,331 | 1,206 |
| Neisseria meningitidis | 0.18 | 3,157 | 3,163 | 3,154 | 3,082 | 628 | 69 | 29623000 | 27333000 | 3,155 | 1,501 | 383 | 3,117 | 3,080 | 3,168 |
| Phocaeicola vulgatus | 0.02 | 648 | 648 | 648 | 639 | 23 | 0 | 16535000 | 5244000 | 646 | 17 | 0 | 0 | 0 | 25 |
| Bifidobacterium adolescentis | 0.02 | 108 | 108 | 108 | 104 | 0 | 3 | 481000 | 550000 | 107 | 30 | 0 | 0 | 0 | 0 |
| Deinococcus radiodurans | 0.02 | 542 | 542 | 542 | 534 | 125 | 10 | 4126000 | 3640000 | 542 | 491 | 0 | 0 | 0 | 15 |
| Enterococcus faecalis | 0.02 | 224 | 224 | 205 | 197 | 0 | 2 | 273000 | 1211000 | 201 | 43 | 0 | 0 | 0 | 0 |
| Schaalia odontolytica | 0.02 | 142 | 142 | 142 | 22 | 0 | 0 | 654000 | 637000 | 0 | 67 | 0 | 0 | 0 | 0 |
| Other | 0 | 51,376 | 62,360 | 8,961 | 7,564 | 34,128 | 634 | 1620052999 | 1472289000 | 2,596 | 1,591 | 0 | 0 | 0 | 0 |

Sourmash results are reported as abundance-projected base pair estimates, which is a projection of the number of base pairs that the percent of matched k-mers represents.

**Supplementary Table S2.** Genus-level read counts for HiFi ATCC MSA-1003.

| **Genus** | **Theoretical** | **Kraken** | **Bracken** | **Centrifuge-h22** | **Centrifuge-h500** | **Metaphlan3** | **mOTUs** | **Sourmash-k31** | **Sourmash-k51** | **Metamaps** | **MMseqs2** | **MEGAN-Prot** | **MEGAN-Nuc-HiFi** | **MEGAN-Nuc-ONT** | **BugSeq-V2** |
| --- | --- | --- | --- | --- | --- | --- | --- | --- | --- | --- | --- | --- | --- | --- | --- |
| Staphylococcus | 19.8 | 248,954 | 248,950 | 248,767 | 242,862 | 200,354 | 3618 | 1908684000 | 1895141000 | 248,538 | 58,266 | 165,670 | 248,199 | 247,479 | 224,995 |
| Streptococcus | 19.8 | 245,513 | 245,459 | 245,461 | 240,208 | 159,899 | 5401 | 1976048000 | 1889786000 | 245,570 | 240,774 | 196,348 | 244,823 | 244,813 | 242,557 |
| Escherichia | 18 | 488,686 | 565,952 | 475,848 | 415,503 | 317,483 | 5285 | 4930014000 | 4879628000 | 609,822 | 154,999 | 107,340 | 89,787 | 87,395 | 537,965 |
| Porphyromonas | 18 | 278,214 | 278,214 | 278,243 | 273,542 | 164,142 | 6116 | 2199344000 | 2115307000 | 278,196 | 263,227 | 166,372 | 277,958 | 277,894 | 268,073 |
| Luteovulum | 18 | 806,633 | 806,654 | 806,756 | 788,258 | 94,556 | 0 | 5609127999 | 4979545000 | 806,631 | 211,200 | 350,506 | 806,101 | 805,695 | 519,838 |
| Bacillus | 1.8 | 63,167 | 63,167 | 63,154 | 61,728 | 19,837 | 634 | 512398000 | 497244000 | 63,168 | 44,599 | 43,102 | 46,782 | 43,450 | 61,077 |
| Clostridium | 1.8 | 29,646 | 29,626 | 29,676 | 28,533 | 4,391 | 172 | 147740000 | 137691000 | 29,632 | 29,064 | 25,096 | 29,587 | 29,561 | 17,049 |
| Pseudomonas | 1.8 | 117,038 | 117,058 | 117,272 | 114,667 | 142 | 964 | 932163000 | 895719000 | 115,414 | 44,064 | 75,445 | 110,541 | 108,068 | 114,841 |
| Acinetobacter | 0.18 | 5,838 | 5,838 | 5,837 | 5,724 | 500 | 67 | 52489999 | 50699999 | 5,840 | 3,553 | 3,613 | 2,787 | 2,582 | 5,778 |
| Cutibacterium | 0.18 | 4,796 | 4,797 | 4,796 | 4,712 | 1,905 | 87 | 39940999 | 38878000 | 4,796 | 789 | 1,142 | 4,718 | 4,709 | 4,705 |
| Helicobacter | 0.18 | 1,661 | 1,661 | 1,661 | 1,592 | 495 | 53 | 14347000 | 13361000 | 1,661 | 1,659 | 1,425 | 1,660 | 1,658 | 1,768 |
| Lactobacillus | 0.18 | 1,428 | 1,428 | 1,428 | 1,391 | 492 | 16 | 10059000 | 10635000 | 1,435 | 1,306 | 1,079 | 1,415 | 1,415 | 1,206 |
| Neisseria | 0.18 | 3,158 | 3,163 | 3,154 | 3,082 | 628 | 69 | 29623000 | 27333000 | 3,155 | 3,004 | 2,208 | 3,157 | 3,157 | 3,168 |
| Phocaeicola | 0.02 | 648 | 648 | 648 | 639 | 46 | 0 | 16535000 | 5244000 | 648 | 47 | 0 | 0 | 0 | 25 |
| Bifidobacterium | 0.02 | 108 | 108 | 108 | 104 | 0 | 3 | 481000 | 550000 | 107 | 96 | 0 | 0 | 0 | 0 |
| Deinococcus | 0.02 | 542 | 542 | 542 | 534 | 125 | 10 | 4126000 | 3640000 | 542 | 533 | 316 | 0 | 0 | 15 |
| Enterococcus | 0.02 | 225 | 224 | 205 | 197 | 0 | 2 | 273000 | 1211000 | 201 | 67 | 0 | 0 | 0 | 0 |
| Schaalia | 0.02 | 142 | 142 | 142 | 22 | 0 | 0 | 654000 | 637000 | 92 | 68 | 0 | 0 | 0 | 0 |
| Other | 0 | 42,939 | 44,512 | 760 | 363 | 28,966 | 0 | 774000 | 553000 | 2,136 | 1,990 | 0 | 0 | 0 | 0 |

Sourmash results are reported as abundance-projected base pair estimates, which is a projection of the number of base pairs that the percent of matched k-mers represents.

**Supplementary Table S3.** Species-level read counts for HiFi Zymo D6331.

| **Species** | **Theoretical** | **Kraken** | **Bracken** | **Centrifuge-h22** | **Centrifuge-h500** | **Metaphlan3** | **mOTUs** | **Sourmash-k31** | **Sourmash-k51** | **Metamaps** | **MMseqs2** | **MEGAN-Prot** | **MEGAN-Nuc-HiFi** | **MEGAN-Nuc-ONT** | **BugSeq-V2** |
| --- | --- | --- | --- | --- | --- | --- | --- | --- | --- | --- | --- | --- | --- | --- | --- |
| Faecalibacterium prausnitzii | 14 | 389,244 | 391,410 | 386,140 | 25,671 | 79,782 | 0 | 2232798000 | 2206635000 | 0 | 8,007 | 123,357 | 362,252 | 360,348 | 276,239 |
| Veillonella rogosae | 14 | 0 | 0 | 0 | 0 | 0 | 5064 | 2724365999 | 2182118998 | 0 | 333 | 5,239 | 0 | 0 | 0 |
| Roseburia hominis | 14 | 101,822 | 102,154 | 99,626 | 76,760 | 27,457 | 1540 | 815741000 | 781818999 | 107,429 | 10,001 | 45,264 | 85,589 | 84,925 | 102,836 |
| Bacteroides fragilis | 14 | 370,388 | 372,972 | 371,058 | 345,450 | 61,645 | 2157 | 3345958000 | 3290304999 | 372,729 | 66,083 | 160,813 | 352,672 | 351,235 | 427,436 |
| Prevotella corporis | 14 | 0 | 0 | 0 | 0 | 44,310 | 3452 | 3090188999 | 3092246999 | 0 | 94,825 | 29,504 | 0 | 0 | 0 |
| Escherichia coli | 6 | 296,090 | 318,823 | 295,620 | 279,043 | 142,283 | 2705 | 1000160000 | 831901000 | 321,736 | 83,700 | 46,022 | 46,057 | 63,100 | 374,948 |
| Bifidobacterium adolescentis | 6 | 38,924 | 39,148 | 38,761 | 38,225 | 9,217 | 800 | 354089000 | 338363000 | 38,762 | 10,370 | 13,505 | 36,983 | 36,769 | 39,790 |
| Fusobacterium nucleatum | 6 | 108,383 | 108,395 | 107,585 | 54,547 | 35,421 | 1962 | 758551000 | 743456000 | 107,352 | 60,120 | 55,183 | 26,958 | 25,743 | 85,888 |
| Limosilactobacillus fermentum | 6 | 23,018 | 23,152 | 23,177 | 22,531 | 8,330 | 497 | 164152000 | 156817000 | 22,915 | 10,243 | 12,492 | 14,921 | 14,076 | 17,431 |
| Clostridioides difficile | 1.5 | 53,950 | 54,778 | 53,052 | 51,449 | 14,105 | 596 | 448176000 | 442126999 | 52,865 | 48,928 | 27,256 | 52,783 | 52,788 | 51,852 |
| Akkermansia muciniphila | 1.5 | 39,515 | 39,515 | 39,512 | 38,011 | 10,985 | 785 | 353163000 | 332318000 | 37,945 | 7,215 | 15,450 | 39,269 | 39,255 | 41,083 |
| Candida albicans | 1.5 | 4,532 | 4,533 | 0 | 0 | 470 | 24 | 11060000 | 10294000 | 0 | 3,696 | 316 | 4,509 | 4,509 | 3,247 |
| Saccharomyces cerevisiae | 1.4 | 3,987 | 3,987 | 0 | 0 | 636 | 8 | 12522000 | 3074000 | 3,996 | 810 | 0 | 0 | 0 | 3,134 |
| Methanobrevibacter smithii | 0.1 | 1,045 | 1,045 | 1,047 | 1,016 | 437 | 0 | 153000 | 166000 | 1,047 | 32 | 0 | 1,043 | 1,043 | 1,160 |
| Salmonella enterica | 0.01 | 532 | 565 | 432 | 333 | 12 | 0 | 0 | 0 | 344 | 169 | 0 | 0 | 0 | 0 |
| Enterococcus faecalis | 0.001 | 19 | 19 | 37 | 15 | 0 | 0 | 46414000 | 44771999 | 14 | 0 | 2,322 | 0 | 0 | 0 |
| Clostridium perfringens | 0.0001 | 3 | 0 | 3 | 0 | 0 | 0 | 38849000 | 38605999 | 0 | 0 | 2,196 | 0 | 0 | 0 |
| Other | 0 | 501,337 | 507,076 | 499,102 | 38,262 | 89,558 | 305 | 3315751999 | 3216723998 | 355,765 | 14,994 | 4,734 | 140,685 | 143,048 | 0 |

Sourmash results are reported as abundance-projected base pair estimates, which is a projection of the number of base pairs that the percent of matched k-mers represents.

**Supplementary Table S4.** Genus-level read counts for HiFi Zymo D6331.

| **Genus** | **Theoretical** | **Kraken** | **Bracken** | **Centrifuge-h22** | **Centrifuge-h500** | **Metaphlan3** | **mOTUs** | **Sourmash-k31** | **Sourmash-k51** | **Metamaps** | **MMseqs2** | **MEGAN-Prot** | **MEGAN-Nuc-HiFi** | **MEGAN-Nuc-ONT** | **BugSeq-V2** |
| --- | --- | --- | --- | --- | --- | --- | --- | --- | --- | --- | --- | --- | --- | --- | --- |
| Faecalibacterium | 14 | 389,244 | 391,410 | 386,140 | 25,671 | 79,782 | 0 | 2232798000 | 2245889000 | 0 | 360,076 | 293,611 | 362,265 | 360,366 | 276,239 |
| Veillonella | 14 | 320,863 | 320,904 | 320,202 | 26,267 | 78,789 | 5064 | 2724366000 | 2182119000 | 291,225 | 302,882 | 224,485 | 234,591 | 232,084 | 235,670 |
| Roseburia | 14 | 105,598 | 105,935 | 103,160 | 77,747 | 27,457 | 1540 | 815741000 | 781818999 | 107,429 | 94,312 | 61,749 | 92,416 | 91,976 | 102,836 |
| Bacteroides | 14 | 379,555 | 381,293 | 380,993 | 347,253 | 61,645 | 2157 | 3345958000 | 3290304999 | 372,818 | 312,501 | 280,562 | 364,544 | 365,031 | 427,436 |
| Prevotella | 14 | 128,283 | 128,349 | 117,508 | 2,107 | 44,310 | 3452 | 3090188999 | 3092246999 | 27,225 | 143,826 | 99,498 | 104,312 | 113,480 | 145,616 |
| Escherichia | 6 | 300,479 | 321,116 | 295,918 | 279,679 | 142,283 | 2705 | 1000160000 | 831901000 | 321,736 | 90,137 | 53,788 | 47,839 | 68,082 | 375,159 |
| Bifidobacterium | 6 | 38,966 | 39,181 | 38,836 | 38,269 | 9,360 | 800 | 354089000 | 338363000 | 38,793 | 35,784 | 29,100 | 38,003 | 38,117 | 39,790 |
| Fusobacterium | 6 | 109,137 | 109,143 | 109,245 | 54,802 | 36,353 | 1962 | 758551000 | 743456000 | 108,955 | 107,569 | 89,608 | 34,848 | 33,166 | 85,888 |
| Limosilactobacillus | 6 | 23,589 | 23,716 | 23,285 | 22,531 | 16,660 | 497 | 164152000 | 156817000 | 22,976 | 10,953 | 14,178 | 22,870 | 22,794 | 17,431 |
| Clostridioides | 1.5 | 53,950 | 54,778 | 53,052 | 51,449 | 14,105 | 596 | 448176000 | 442126999 | 52,865 | 48,946 | 27,282 | 52,783 | 52,788 | 51,852 |
| Akkermansia | 1.5 | 39,515 | 39,515 | 39,513 | 38,011 | 10,985 | 785 | 353163000 | 332318000 | 37,945 | 34,289 | 25,790 | 39,275 | 39,263 | 41,083 |
| Methanobrevibacter | 0.1 | 1,045 | 1,045 | 1,056 | 1,016 | 437 | 24 | 11060000 | 10294000 | 1,047 | 1,040 | 800 | 1,043 | 1,043 | 1,160 |
| Salmonella | 0.01 | 637 | 655 | 483 | 353 | 12 | 8 | 12522000 | 3074000 | 344 | 319 | 0 | 0 | 0 | 0 |
| Enterococcus | 0.001 | 419 | 434 | 652 | 15 | 0 | 0 | 153000 | 166000 | 15 | 1 | 0 | 0 | 0 | 0 |
| Clostridium | 0.0001 | 309 | 294 | 422 | 2 | 0 | 0 | 0 | 0 | 44 | 703 | 381 | 0 | 0 | 0 |
| Candida | 1.5 | 4,642 | 4,638 | 116 | 0 | 470 | 0 | 46414000 | 44771999 | 4,379 | 4,177 | 2,322 | 4,509 | 4,509 | 3,247 |
| Saccharomyces | 1.4 | 3,993 | 3,987 | 0 | 0 | 653 | 0 | 39664000 | 38605999 | 3,996 | 3,401 | 2,701 | 3,819 | 3,874 | 3,555 |
| Other | 0 | 41,827 | 41,214 | 54,045 | 9,382 | 9,682 | 0 | 0 | 774000 | 19,907 | 2,193 | 0 | 999 | 1,104 | 0 |

Sourmash results are reported as abundance-projected base pair estimates, which is a projection of the number of base pairs that the percent of matched k-mers represents.

**Supplementary Table S5.** Species-level read counts for ONT R10 Zymo D6300.

| **Species** | **Theoretical** | **Kraken** | **Bracken** | **Centrifuge-h22** | **Centrifuge-h500** | **Metaphlan3** | mOTUs | Sourmash-k31 | Sourmash-k51 | **Metamaps** | **MMseqs2** | **MEGAN-Prot** | **MEGAN-Nuc-HiFi** | **MEGAN-Nuc-ONT** | **BugSeq-V2** |
| --- | --- | --- | --- | --- | --- | --- | --- | --- | --- | --- | --- | --- | --- | --- | --- |
| Pseudomonas aeruginosa | 12 | 22,862 | 22,963 | 45,410 | 1,562 | 0 | 148 | 53062999 | 134319000 | 22,535 | 273 | 5,252 | 21,554 | 21,513 | 36,720 |
| Escherichia coli | 12 | 25,249 | 25,425 | 23,173 | 72 | 13,969 | 120 | 456323000 | 180590000 | 26,006 | 475 | 2,345 | 1,076 | 1,001 | 42,010 |
| Salmonella enterica | 12 | 27,572 | 27,642 | 27,153 | 191 | 6,122 | 186 | 200771999 | 121372000 | 27,556 | 743 | 784 | 256 | 265 | 42,111 |
| Limosilactobacillus fermentum | 12 | 50,208 | 50,214 | 52,225 | 141 | 18,401 | 207 | 110298000 | 65838000 | 50,231 | 897 | 24,180 | 48,170 | 47,940 | 23,157 |
| Enterococcus faecalis | 12 | 29,670 | 29,718 | 29,416 | 359 | 7,873 | 202 | 52268000 | 42066999 | 29,151 | 242 | 10,853 | 27,873 | 27,712 | 26,744 |
| Staphylococcus aureus | 12 | 30,646 | 30,686 | 29,857 | 435 | 9,829 | 226 | 156813000 | 101820000 | 30,376 | 721 | 14,036 | 8,695 | 8,305 | 28,861 |
| Listeria monocytogenes | 12 | 33,962 | 34,004 | 33,103 | 427 | 10,255 | 228 | 144622000 | 94660000 | 34,461 | 353 | 9,893 | 31,754 | 31,601 | 29,642 |
| Bacillus subtilis | 12 | 29,045 | 32,274 | 6,282 | 9 | 0 | 186 | 160620000 | 102853000 | 41,034 | 55 | 5,402 | 1,410 | 1,548 | 32,302 |
| Saccharomyces cerevisiae | 2 | 4,986 | 5,000 | 0 | 0 | 744 | 0 | 20947000 | 14583000 | 5,338 | 111 | 2,184 | 2,994 | 2,926 | 2,606 |
| Cryptococcus neoformans | 2 | 7,554 | 7,556 | 0 | 0 | 245 | 0 | 19694000 | 12079000 | 0 | 1,300 | 774 | 4,894 | 4,705 | 1,306 |
| Other | 0 | 8,713 | 9,216 | 2,443 | 4 | 13,760 | 0 | 2142000 | 551999 | 1,396 | 226 | 0 | 0 | 0 | 90 |

Sourmash results are reported as abundance-projected base pair estimates, which is a projection of the number of base pairs that the percent of matched k-mers represents.

**Supplementary Table S6.** Genus-level read counts for ONT R10 Zymo D6300.

| **Genus** | **Theoretical** | **Kraken** | **Bracken** | **Centrifuge-h22** | **Centrifuge-h500** | **Metaphlan3** | **mOTUs** | **Sourmash-k31** | **Sourmash-k51** | **Metamaps** | **MMseqs2** | **MEGAN-Prot** | **MEGAN-Nuc-HiFi** | **MEGAN-Nuc-ONT** | **BugSeq-V2** |
| --- | --- | --- | --- | --- | --- | --- | --- | --- | --- | --- | --- | --- | --- | --- | --- |
| Pseudomonas | 12 | 23,098 | 22,989 | 22,815 | 985 | 36 | 148 | 53062999 | 134319000 | 22,720 | 616 | 5,365 | 21,669 | 21,650 | 36,720 |
| Escherichia | 12 | 25,498 | 25,624 | 23,264 | 77 | 13,969 | 120 | 456652000 | 180590000 | 26,006 | 505 | 2,551 | 1,085 | 1,010 | 42,010 |
| Salmonella | 12 | 27,625 | 27,655 | 27,237 | 195 | 6,122 | 186 | 200846999 | 121372000 | 27,556 | 10,122 | 1,468 | 509 | 526 | 42,111 |
| Limosilactobacillus | 12 | 50,214 | 50,214 | 52,227 | 141 | 36,802 | 207 | 110298000 | 65838000 | 50,663 | 1,379 | 36,283 | 48,878 | 48,797 | 23,157 |
| Enterococcus | 12 | 29,721 | 29,741 | 29,431 | 359 | 7,873 | 202 | 52268000 | 42066999 | 29,183 | 454 | 14,062 | 27,908 | 27,834 | 26,744 |
| Staphylococcus | 12 | 30,671 | 30,686 | 30,107 | 526 | 11,049 | 226 | 156813000 | 101820000 | 30,485 | 2,507 | 20,135 | 8,803 | 8,424 | 28,861 |
| Listeria | 12 | 33,978 | 34,004 | 33,138 | 428 | 11,569 | 228 | 144622000 | 94660000 | 34,461 | 12,061 | 18,362 | 31,909 | 31,782 | 29,642 |
| Bacillus | 12 | 40,974 | 40,794 | 41,618 | 325 | 9,137 | 186 | 161704000 | 103306000 | 41,051 | 2,987 | 9,187 | 2,224 | 2,544 | 32,302 |
| Saccharomyces | 2 | 4,992 | 5,000 | 0 | 0 | 744 | 0 | 21280000 | 14583000 | 5,338 | 3,210 | 2,765 | 4,530 | 4,491 | 3,272 |
| Cryptococcus | 2 | 7,558 | 7,556 | 0 | 0 | 617 | 0 | 19694000 | 12079000 | 0 | 4,883 | 2,218 | 4,905 | 4,711 | 1,306 |
| Other | 0 | 686 | 446 | 633 | 2 | 1,801 | 0 | 321000 | 99000 | 501 | 128 | 0 | 0 | 0 | 0 |

Sourmash results are reported as abundance-projected base pair estimates, which is a projection of the number of base pairs that the percent of matched k-mers represents.

**Supplementary Table S7.** Species-level read counts for ONT Q20 Zymo D6300.

| **Species** | **Theoretical** | **Kraken** | **Bracken** | **Centrifuge-h22** | **Centrifuge-h500** | **Metaphlan3** | **mOTUs** | **Sourmash-k31** | **Sourmash-k51** | **Metamaps** | **MMseqs2** | **MEGAN-Prot** | **MEGAN-Nuc-HiFi** | **MEGAN-Nuc-ONT** | **BugSeq-V2** |
| --- | --- | --- | --- | --- | --- | --- | --- | --- | --- | --- | --- | --- | --- | --- | --- |
| Pseudomonas aeruginosa | 12 | 181,001 | 183,502 | 371,858 | 225,954 | 0 | 115 | 30668999 | 100564999 | 178,347 | 19,810 | 83,821 | 154,419 | 153,798 | 192,544 |
| Escherichia coli | 12 | 220,685 | 231,957 | 191,294 | 59,701 | 56,566 | 235 | 372175000 | 220446000 | 246,354 | 32,754 | 71,249 | 8,021 | 7,721 | 240,309 |
| Salmonella enterica | 12 | 271,319 | 276,738 | 246,850 | 142,882 | 35,946 | 263 | 184450000 | 167039000 | 266,781 | 52,039 | 19,628 | 5,668 | 5,569 | 253,522 |
| Limosilactobacillus fermentum | 12 | 213,317 | 213,372 | 230,292 | 107,458 | 83,935 | 1543 | 528050000 | 434214000 | 210,701 | 13,766 | 82,785 | 181,931 | 180,905 | 180,016 |
| Enterococcus faecalis | 12 | 230,329 | 231,015 | 234,388 | 125,340 | 61,084 | 354 | 52292000 | 57768000 | 226,222 | 6,494 | 97,732 | 195,631 | 194,170 | 240,292 |
| Staphylococcus aureus | 12 | 218,817 | 219,363 | 217,460 | 130,522 | 65,193 | 535 | 227406000 | 190643999 | 218,809 | 23,736 | 121,395 | 44,954 | 41,888 | 211,288 |
| Listeria monocytogenes | 12 | 238,135 | 238,884 | 235,546 | 136,621 | 54,720 | 335 | 147293000 | 127000000 | 243,540 | 13,874 | 84,355 | 200,790 | 200,024 | 251,609 |
| Bacillus subtilis | 12 | 160,638 | 187,653 | 32,701 | 5,780 | 0 | 296 | 179629000 | 156918000 | 269,877 | 479 | 40,457 | 9,890 | 10,826 | 180,071 |
| Saccharomyces cerevisiae | 2 | 47,370 | 47,491 | 0 | 0 | 4,380 | 0 | 37167000 | 33412000 | 50,814 | 2,992 | 23,127 | 27,246 | 26,841 | 32,140 |
| Cryptococcus neoformans | 2 | 53,681 | 53,708 | 0 | 0 | 1,694 | 0 | 74842000 | 61960000 | 0 | 23,599 | 4,943 | 30,096 | 29,216 | 34,284 |
| Other | 0 | 89,588 | 111,324 | 13,138 | 1,872 | 90,011 | 2 | 2895000 | 1809000 | 33,497 | 1,675 | 0 | 0 | 0 | 418 |

Sourmash results are reported as abundance-projected base pair estimates, which is a projection of the number of base pairs that the percent of matched k-mers represents.

**Supplementary Table S8.** Species-level read counts for ONT Q20 Zymo D6300.

| **Genus** | **Theoretical** | **Kraken** | **Bracken** | **Centrifuge-h22** | **Centrifuge-h500** | **Metaphlan3** | **mOTUs** | **Sourmash-k31** | **Sourmash-k51** | **Metamaps** | **MMseqs2** | **MEGAN-Prot** | **MEGAN-Nuc-HiFi** | **MEGAN-Nuc-ONT** | **BugSeq-V2** |
| --- | --- | --- | --- | --- | --- | --- | --- | --- | --- | --- | --- | --- | --- | --- | --- |
| Pseudomonas | 12 | 185,227 | 184,872 | 187,731 | 120,858 | 120 | 115 | 30668999 | 100564999 | 180,136 | 35,572 | 84,688 | 154,423 | 153,800 | 192,544 |
| Escherichia | 12 | 232,016 | 240,398 | 192,543 | 60,759 | 56,566 | 235 | 372175000 | 220446000 | 246,354 | 34,907 | 76,499 | 8,416 | 8,165 | 240,309 |
| Salmonella | 12 | 273,306 | 277,107 | 248,024 | 146,655 | 35,946 | 263 | 184450000 | 168303000 | 266,781 | 173,434 | 30,985 | 5,851 | 5,736 | 253,522 |
| Limosilactobacillus | 12 | 213,353 | 213,384 | 230,318 | 107,458 | 167,870 | 1543 | 528050000 | 434312000 | 213,033 | 17,468 | 131,359 | 184,631 | 184,093 | 180,016 |
| Enterococcus | 12 | 230,741 | 231,174 | 234,541 | 125,563 | 61,084 | 354 | 52292000 | 57768000 | 226,420 | 10,038 | 119,863 | 195,997 | 195,481 | 240,292 |
| Staphylococcus | 12 | 218,998 | 219,363 | 219,884 | 141,708 | 78,788 | 537 | 227406000 | 190643999 | 219,818 | 79,906 | 146,419 | 45,692 | 42,707 | 211,300 |
| Listeria | 12 | 238,311 | 238,922 | 235,769 | 136,853 | 65,721 | 335 | 147293000 | 127000000 | 243,540 | 169,083 | 135,029 | 202,639 | 201,896 | 251,609 |
| Bacillus | 12 | 268,179 | 268,155 | 276,222 | 140,041 | 45,575 | 296 | 180013000 | 157107000 | 270,062 | 115,376 | 66,935 | 15,495 | 17,453 | 262,251 |
| Saccharomyces | 2 | 47,476 | 47,587 | 0 | 0 | 4,794 | 0 | 39030000 | 33412000 | 50,814 | 35,725 | 27,334 | 39,079 | 38,866 | 39,461 |
| Cryptococcus | 2 | 53,707 | 53,720 | 0 | 0 | 4,948 | 0 | 74842000 | 61960000 | 0 | 41,730 | 4,949 | 30,139 | 29,247 | 34,284 |
| Other | 0 | 21,709 | 20,378 | 4,643 | 922 | 17,156 | 0 | 648000 | 258000 | 26,713 | 1,201 | 0 | 0 | 0 | 0 |

Sourmash results are reported as abundance-projected base pair estimates, which is a projection of the number of base pairs that the percent of matched k-mers represents.

**Supplementary Table S9.** The number of species detected at each relative abundance level in the HiFi ATCC MSA-1003 and HiFi Zymo D6331 datasets, based on the minimum 0.001% of total reads threshold.

|  |  | **Number of Species Detected at Abundance Level** | | | | | | |
| --- | --- | --- | --- | --- | --- | --- | --- | --- |
| **Dataset** | **Method** | **14-18%** | **6%** | **1.5-1.8%** | **0.10-0.18%** | **0.01-0.02%** | **0.001%** | **0.0001%** |
| HiFi ATCC MSA-1003 | Kraken2 | **5/5** | - | **5/5** | **5/5** | **5/5** | - | - |
|  | Bracken | **5/5** | - | **5/5** | **5/5** | **5/5** | - | - |
|  | Centrifuge-h22 | **5/5** | - | **5/5** | **5/5** | **5/5** | - | - |
|  | Centrifuge-h500 | **5/5** | - | **5/5** | **5/5** | 4/5 | - | - |
|  | MetaPhlAn3 | **5/5** | - | 3/5 | 4/5 | 1/5 | - | - |
|  | mOTUs | 4/5 | - | 4/5 | **5/5** | 3/5 | - | - |
|  | Sourmash-K31 | **5/5** | - | **5/5** | **5/5** | **5/5** | - | - |
|  | Sourmash-K51 | **5/5** | - | **5/5** | **5/5** | **5/5** | - | - |
|  | MetaMaps | **5/5** | - | **5/5** | **5/5** | 4/5 | - | - |
|  | MMseqs2 | **5/5** | - | **5/5** | **5/5** | 4/5 | - | - |
|  | MEGAN-LR-prot | **5/5** | - | **5/5** | **5/5** | 0/5 | - | - |
|  | MEGAN-LR-nuc-HiFi | **5/5** | - | **5/5** | **5/5** | 0/5 | - | - |
|  | MEGAN-LR-nuc-ONT | **5/5** | - | **5/5** | **5/5** | 0/5 | - | - |
|  | BugSeq-V2 | **5/5** | - | **5/5** | **5/5** | 1/5 | - | - |
| HiFi Zymo D6331 | Kraken2 | **3/3** | **4/4** | **4/4** | **1/1** | **1/1** | **1/1** | 0/1 |
|  | Bracken | **3/3** | **4/4** | **4/4** | **1/1** | **1/1** | **1/1** | 0/1 |
|  | Centrifuge-h22 | **3/3** | **4/4** | 2/4 | **1/1** | **1/1** | **1/1** | 0/1 |
|  | Centrifuge-h500 | **3/3** | **4/4** | 2/4 | **1/1** | **1/1** | 0/1 | 0/1 |
|  | MetaPhlAn3 | **3/3** | **4/4** | **4/4** | **1/1** | 0/1 | 0/1 | 0/1 |
|  | mOTUs | **3/3** | **4/4** | 2/4 | **1/1** | **1/1** | 0/1 | 0/1 |
|  | Sourmash-K31 | **3/3** | **4/4** | **4/4** | **1/1** | **1/1** | **1/1** | 0/1 |
|  | Sourmash-K51 | **3/3** | **4/4** | **4/4** | **1/1** | **1/1** | **1/1** | 0/1 |
|  | MetaMaps | 2/3 | **4/4** | 3/4 | **1/1** | **1/1** | 0/1 | 0/1 |
|  | MMseqs2 | **3/3** | **4/4** | **4/4** | **1/1** | **1/1** | 0/1 | 0/1 |
|  | MEGAN-LR-prot | **3/3** | **4/4** | **4/4** | **1/1** | 0/1 | 0/1 | 0/1 |
|  | MEGAN-LR-nuc-HiFi | **3/3** | **4/4** | 3/4 | **1/1** | 0/1 | 0/1 | 0/1 |
|  | MEGAN-LR-nuc-ONT | **3/3** | **4/4** | 3/4 | **1/1** | 0/1 | 0/1 | 0/1 |
|  | BugSeq-V2 | **3/3** | **4/4** | **4/4** | **1/1** | 0/1 | 0/1 | 0/1 |

Perfect detection scores are highlighted in bold. For Zymo D6331 two species were excluded at the 14% abundance level, which were missing from the databases for several methods.

**Supplementary Table S10.** Genus-level detection results based on the minimum 0.001% of total reads threshold.

| **Dataset** | **Method Type** | **Profiling Method** | **True Positives** | **False Positives** | **False Negatives** | **Precision** | **Recall** | **F1** | **F0.5** | **L1** |
| --- | --- | --- | --- | --- | --- | --- | --- | --- | --- | --- |
| HiFi ATCC-MSA1003 | Short read | Kraken2 | 18 | 28 | 0 | 0.39 | 1.00 | 0.56 | 0.45 | 50.8 |
| (18 genera, staggered) |  | Bracken | 18 | 28 | 0 | 0.39 | 1.00 | 0.56 | 0.45 | 53.1 |
|  |  | Centrifuge-h22 | 18 | 5 | 0 | 0.78 | 1.00 | 0.88 | 0.82 | 49.2 |
|  |  | Centrifuge-h500 | 17 | 4 | 1 | 0.81 | 0.94 | 0.87 | 0.83 | 47.5 |
|  |  | Metaphlan3 | 15 | 28 | 3 | 0.35 | 0.83 | 0.49 | 0.39 | 34.8 |
|  |  | mOTUs | 11 | 0 | 7 | 1.00 | 0.61 | 0.76 | 0.89 | 45.8 |
|  | General | Sourmash-k31 | 18 | 2 | 0 | 0.90 | 1.00 | 0.95 | 0.92 | 51.6 |
|  |  | Sourmash-k51 | 18 | 1 | 0 | 0.95 | 1.00 | 0.97 | 0.96 | 50.2 |
|  | Long read | Metamaps | 18 | 4 | 0 | 0.82 | 1.00 | 0.90 | 0.85 | 53.1 |
|  |  | MMseqs2 | 18 | 2 | 0 | 0.90 | 1.00 | 0.95 | 0.92 | 35.8 |
|  |  | MEGAN-LR-Prot | 14 | 0 | 4 | 1.00 | 0.78 | 0.87 | 0.94 | 40.2 |
|  |  | MEGAN-LR-Nuc-HiFi | 13 | 0 | 5 | 1.00 | 0.72 | 0.84 | 0.93 | 60.1 |
|  |  | MEGAN-LR-Nuc-ONT | 13 | 0 | 5 | 1.00 | 0.72 | 0.84 | 0.93 | 60.0 |
|  |  | BugSeq-V2 | 14 | 0 | 4 | 1.00 | 0.78 | 0.88 | 0.95 | 44.3 |
| HiFi Zymo-D6331 | Short read | Kraken2 | 17 | 130 | 0 | 0.12 | 1.00 | 0.21 | 0.14 | 40.4 |
| (17 genera, staggered) |  | Bracken | 17 | 122 | 0 | 0.12 | 1.00 | 0.22 | 0.15 | 40.7 |
|  |  | Centrifuge-h22 | 16 | 213 | 1 | 0.07 | 0.94 | 0.13 | 0.09 | 41.3 |
|  |  | Centrifuge-h500 | 13 | 16 | 4 | 0.45 | 0.77 | 0.57 | 0.49 | 87.0 |
|  |  | Metaphlan3 | 14 | 17 | 3 | 0.45 | 0.82 | 0.58 | 0.50 | 42.2 |
|  |  | mOTUs | 11 | 0 | 6 | 1.00 | 0.65 | 0.79 | 0.90 | 62.8 |
|  | General | Sourmash-k31 | 16 | 0 | 1 | 1.00 | 0.94 | 0.97 | 0.99 | 41.6 |
|  |  | Sourmash-k51 | 16 | 1 | 1 | 0.94 | 0.94 | 0.94 | 0.94 | 41.5 |
|  | Long read | Metamaps | 15 | 35 | 2 | 0.30 | 0.88 | 0.45 | 0.35 | 68.8 |
|  |  | MMseqs2 | 16 | 9 | 1 | 0.64 | 0.94 | 0.76 | 0.68 | 55.1 |
|  |  | MEGAN-LR-Prot | 15 | 0 | 2 | 1.00 | 0.88 | 0.93 | 0.97 | 58.7 |
|  |  | MEGAN-LR-Nuc-HiFi | 14 | 1 | 3 | 0.93 | 0.82 | 0.88 | 0.91 | 63.1 |
|  |  | MEGAN-LR-Nuc-ONT | 14 | 1 | 3 | 0.93 | 0.82 | 0.88 | 0.91 | 61.1 |
|  |  | BugSeq-V2 | 14 | 0 | 3 | 1.00 | 0.82 | 0.90 | 0.96 | 43.8 |
| ONT R10 Zymo-D6300 | Short read | Kraken2 | 10 | 46 | 0 | 0.18 | 1.00 | 0.30 | 0.21 | 21.0 |
| (10 genera, even) |  | Bracken | 10 | 6 | 0 | 0.63 | 1.00 | 0.77 | 0.68 | 20.8 |
|  |  | Centrifuge-h22 | 8 | 16 | 2 | 0.33 | 0.80 | 0.47 | 0.38 | 26.0 |
|  |  | Centrifuge-h500 | 8 | 0 | 2 | 1.00 | 0.80 | 0.89 | 0.95 | 55.8 |
|  |  | Metaphlan3 | 10 | 10 | 0 | 0.50 | 1.00 | 0.67 | 0.56 | 57.4 |
|  |  | mOTUs | 8 | 0 | 2 | 1.00 | 0.80 | 0.89 | 0.95 | 20.3 |
|  | General | Sourmash-k31 | 10 | 2 | 0 | 0.83 | 1.00 | 0.91 | 0.86 | 47.5 |
|  |  | Sourmash-k51 | 10 | 1 | 0 | 0.91 | 1.00 | 0.95 | 0.93 | 28.2 |
|  | Long read | Metamaps | 9 | 18 | 1 | 0.33 | 0.90 | 0.49 | 0.38 | 22.5 |
|  |  | MMseqs2 | 10 | 21 | 0 | 0.32 | 1.00 | 0.49 | 0.37 | 100.5 |
|  |  | MEGAN-LR-Prot | 10 | 0 | 0 | 1.00 | 1.00 | 1.00 | 1.00 | 63.0 |
|  |  | MEGAN-LR-Nuc-HiFi | 10 | 0 | 0 | 1.00 | 1.00 | 1.00 | 1.00 | 79.4 |
|  |  | MEGAN-LR-Nuc-ONT | 10 | 0 | 0 | 1.00 | 1.00 | 1.00 | 1.00 | 79.5 |
|  |  | BugSeq-V2 | 10 | 0 | 0 | 1.00 | 1.00 | 1.00 | 1.00 | 19.1 |
| ONT Q20 Zymo-D6300 | Short read | Kraken2 | 10 | 29 | 0 | 0.26 | 1.00 | 0.41 | 0.30 | 11.0 |
| (10 genera, even) |  | Bracken | 10 | 24 | 0 | 0.29 | 1.00 | 0.45 | 0.34 | 10.9 |
|  |  | Centrifuge-h22 | 8 | 16 | 2 | 0.33 | 0.80 | 0.47 | 0.38 | 14.4 |
|  |  | Centrifuge-h500 | 8 | 4 | 2 | 0.67 | 0.80 | 0.73 | 0.69 | 21.7 |
|  |  | Metaphlan3 | 10 | 21 | 0 | 0.32 | 1.00 | 0.49 | 0.37 | 50.4 |
|  |  | mOTUs | 8 | 0 | 2 | 1.00 | 0.80 | 0.89 | 0.95 | 65.1 |
|  | General | Sourmash-k31 | 10 | 1 | 0 | 0.91 | 1.00 | 0.95 | 0.93 | 55.2 |
|  |  | Sourmash-k51 | 10 | 1 | 0 | 0.91 | 1.00 | 0.95 | 0.93 | 41.3 |
|  | Long read | Metamaps | 9 | 22 | 1 | 0.29 | 0.90 | 0.44 | 0.34 | 13.6 |
|  |  | MMseqs2 | 10 | 8 | 0 | 0.56 | 1.00 | 0.71 | 0.61 | 70.2 |
|  |  | MEGAN-LR-Prot | 10 | 0 | 0 | 1.00 | 1.00 | 1.00 | 1.00 | 35.9 |
|  |  | MEGAN-LR-Nuc-HiFi | 10 | 0 | 0 | 1.00 | 1.00 | 1.00 | 1.00 | 78.9 |
|  |  | MEGAN-LR-Nuc-ONT | 10 | 0 | 0 | 1.00 | 1.00 | 1.00 | 1.00 | 79.1 |
|  |  | BugSeq-V2 | 10 | 0 | 0 | 1.00 | 1.00 | 1.00 | 1.00 | 11.1 |

**Supplementary Table S11.** Species-level detection results based on the minimum 0.1% of total reads threshold.

| **Dataset** | **Method Type** | **Profiling Method** | **True Positives** | **False Positives** | **False Negatives** | **Precision** | **Recall** | **F1** | **F0.5** |
| --- | --- | --- | --- | --- | --- | --- | --- | --- | --- |
| HiFi ATCC-MSA1003 | Short read | Kraken2 | 13 | 6 | 7 | 0.68 | 0.65 | 0.67 | 0.68 |
| (20 species, staggered) |  | Bracken | 13 | 6 | 7 | 0.68 | 0.65 | 0.67 | 0.68 |
|  |  | Centrifuge-h22 | 13 | 1 | 7 | 0.93 | 0.65 | 0.76 | 0.86 |
|  |  | Centrifuge-h500 | 13 | 1 | 7 | 0.93 | 0.65 | 0.76 | 0.86 |
|  |  | Metaphlan3 | 8 | 2 | 12 | 0.80 | 0.40 | 0.53 | 0.67 |
|  |  | mOTUs | 4 | 0 | 16 | 1.00 | 0.20 | 0.33 | 0.56 |
|  | General | Sourmash-k31 | 20 | 5 | 0 | 0.80 | 1.00 | 0.89 | 0.83 |
|  |  | Sourmash-k51 | 20 | 3 | 0 | 0.87 | 1.00 | 0.93 | 0.89 |
|  | Long read | Metamaps | 13 | 0 | 7 | 1.00 | 0.65 | 0.79 | 0.90 |
|  |  | MMseqs2 | 8 | 0 | 12 | 1.00 | 0.40 | 0.57 | 0.77 |
|  |  | MEGAN-LR-Prot | 10 | 0 | 10 | 1.00 | 0.50 | 0.67 | 0.83 |
|  |  | MEGAN-LR-Nuc-HiFi | 12 | 0 | 8 | 1.00 | 0.60 | 0.75 | 0.88 |
|  |  | MEGAN-LR-Nuc-ONT | 12 | 0 | 8 | 1.00 | 0.60 | 0.75 | 0.88 |
|  |  | BugSeq-V2 | 13 | 0 | 7 | 1.00 | 0.65 | 0.79 | 0.90 |
| HiFi Zymo-D6331 | Short read | Kraken2* | 11 | 5 | 4 | 0.69 | 0.73 | 0.71 | 0.70 |
| (17 species, staggered) |  | Bracken* | 11 | 5 | 4 | 0.69 | 0.73 | 0.71 | 0.70 |
| *based on 15 species |  | Centrifuge-h22* | 9 | 6 | 6 | 0.60 | 0.60 | 0.60 | 0.60 |
|  |  | Centrifuge-h500* | 9 | 2 | 6 | 0.82 | 0.60 | 0.69 | 0.76 |
|  |  | Metaphlan3* | 9 | 0 | 6 | 1.00 | 0.60 | 0.75 | 0.88 |
|  |  | mOTUs | 2 | 0 | 13 | 1.00 | 0.13 | 0.23 | 0.43 |
|  | General | Sourmash-k31 | 14 | 1 | 1 | 0.93 | 0.93 | 0.93 | 0.93 |
|  |  | Sourmash-k51 | 14 | 2 | 1 | 0.88 | 0.93 | 0.90 | 0.89 |
|  | Long read | Metamaps* | 9 | 6 | 6 | 0.60 | 0.60 | 0.60 | 0.60 |
|  |  | MMseqs2* | 10 | 0 | 5 | 1.00 | 0.67 | 0.80 | 0.91 |
|  |  | MEGAN-LR-Prot* | 11 | 0 | 4 | 1.00 | 0.73 | 0.85 | 0.93 |
|  |  | MEGAN-LR-Nuc-HiFi* | 10 | 1 | 5 | 0.91 | 0.67 | 0.77 | 0.85 |
|  |  | MEGAN-LR-Nuc-ONT* | 10 | 1 | 5 | 0.91 | 0.67 | 0.77 | 0.85 |
|  |  | BugSeq-V2* | 11 | 0 | 4 | 1.00 | 0.73 | 0.85 | 0.93 |
| ONT R10 Zymo-D6300 | Short read | Kraken2 | 10 | 4 | 0 | 0.71 | 1.00 | 0.83 | 0.76 |
| (10 species, even) |  | Bracken | 10 | 5 | 0 | 0.67 | 1.00 | 0.80 | 0.71 |
|  |  | Centrifuge-h22 | 8 | 1 | 2 | 0.89 | 0.80 | 0.84 | 0.87 |
|  |  | Centrifuge-h500 | 4 | 0 | 6 | 1.00 | 0.40 | 0.57 | 0.77 |
|  |  | Metaphlan3 | 7 | 4 | 3 | 0.64 | 0.70 | 0.67 | 0.65 |
|  |  | mOTUs | 0 | 0 | 10 | 0.00 | 0.00 | 0.00 | 0.00 |
|  | General | Sourmash-k31 | 10 | 11 | 0 | 0.48 | 1.00 | 0.64 | 0.53 |
|  |  | Sourmash-k51 | 10 | 3 | 0 | 0.77 | 1.00 | 0.87 | 0.81 |
|  | Long read | Metamaps | 9 | 0 | 1 | 1.00 | 0.90 | 0.95 | 0.98 |
|  |  | MMseqs2 | 6 | 0 | 4 | 1.00 | 0.60 | 0.75 | 0.88 |
|  |  | MEGAN-LR-Prot | 10 | 0 | 0 | 1.00 | 1.00 | 1.00 | 1.00 |
|  |  | MEGAN-LR-Nuc-HiFi | 9 | 0 | 1 | 1.00 | 0.90 | 0.95 | 0.98 |
|  |  | MEGAN-LR-Nuc-ONT | 9 | 0 | 1 | 1.00 | 0.90 | 0.95 | 0.98 |
|  |  | BugSeq-V2 | 10 | 0 | 0 | 1.00 | 1.00 | 1.00 | 1.00 |
| ONT Q20 Zymo-D6300 | Short read | Kraken2 | 10 | 11 | 0 | 0.48 | 1.00 | 0.64 | 0.53 |
| (10 species, even) |  | Bracken | 10 | 13 | 0 | 0.44 | 1.00 | 0.61 | 0.49 |
|  |  | Centrifuge-h22 | 8 | 1 | 2 | 0.89 | 0.80 | 0.84 | 0.87 |
|  |  | Centrifuge-h500 | 8 | 0 | 2 | 1.00 | 0.80 | 0.89 | 0.95 |
|  |  | Metaphlan3 | 7 | 6 | 3 | 0.54 | 0.70 | 0.61 | 0.56 |
|  |  | mOTUs | 0 | 0 | 10 | 0.00 | 0.00 | 0.00 | 0.00 |
|  | General | Sourmash-k31 | 10 | 4 | 0 | 0.71 | 1.00 | 0.83 | 0.76 |
|  |  | Sourmash-k51 | 10 | 4 | 0 | 0.71 | 1.00 | 0.83 | 0.76 |
|  | Long read | Metamaps | 9 | 1 | 1 | 0.90 | 0.90 | 0.90 | 0.90 |
|  |  | MMseqs2 | 9 | 0 | 1 | 1.00 | 0.90 | 0.95 | 0.98 |
|  |  | MEGAN-LR-Prot | 10 | 0 | 0 | 1.00 | 1.00 | 1.00 | 1.00 |
|  |  | MEGAN-LR-Nuc-HiFi | 10 | 0 | 0 | 1.00 | 1.00 | 1.00 | 1.00 |
|  |  | MEGAN-LR-Nuc-ONT | 10 | 0 | 0 | 1.00 | 1.00 | 1.00 | 1.00 |
|  |  | BugSeq-V2 | 10 | 0 | 0 | 1.00 | 1.00 | 1.00 | 1.00 |

**Supplementary Table S12.** The number of species detected at each relative abundance level in the HiFi ATCC MSA-1003 and HiFi Zymo D6331 datasets, based on the minimum 0.1% of total reads threshold.

|  |  | **Species Detected at Relative Abundance Level** | | | | | | |
| --- | --- | --- | --- | --- | --- | --- | --- | --- |
| **Dataset** | **Method** | **14-18%** | **6%** | **1.5-1.8%** | **0.10-0.18%** | **0.01-0.02%** | **0.001%** | **0.0001%** |
| HiFi ATCC MSA-1003 | Kraken2 | **5/5** | - | **5/5** | 3/5 | 0/5 | - | - |
|  | Bracken | **5/5** | - | **5/5** | 3/5 | 0/5 | - | - |
|  | Centrifuge-h22 | **5/5** | - | **5/5** | 3/5 | 0/5 | - | - |
|  | Centrifuge-h500 | **5/5** | - | **5/5** | 3/5 | 0/5 | - | - |
|  | MetaPhlAn3 | **5/5** | - | 3/5 | 0/5 | 0/5 | - | - |
|  | mOTUs | 4/5 | - | 0/5 | 0/5 | 0/5 | - | - |
|  | Sourmash-k31 | **5/5** | - | **5/5** | 3/5 | 0/5 | - | - |
|  | Sourmash-k51 | **5/5** | - | **5/5** | 3/5 | 0/5 | - | - |
|  | MetaMaps | **5/5** | - | **5/5** | 3/5 | 0/5 | - | - |
|  | MMseqs2 | **5/5** | - | 3/5 | 0/5 | 0/5 | - | - |
|  | MEGAN-LR-prot | **5/5** | - | **5/5** | 0/5 | 0/5 | - | - |
|  | MEGAN-LR-nuc-HiFi | **5/5** | - | **5/5** | 3/5 | 0/5 | - | - |
|  | MEGAN-LR-nuc-ONT | **5/5** | - | 4/5 | 3/5 | 0/5 | - | - |
|  | BugSeq-V2 | **5/5** | - | **5/5** | 3/5 | 0/5 | - | - |
| HiFi Zymo D6331 | Kraken2 | **3/3** | **4/4** | **4/4** | 0/1 | 0/1 | 0/1 | 0/1 |
|  | Bracken | **3/3** | **4/4** | **4/4** | 0/1 | 0/1 | 0/1 | 0/1 |
|  | Centrifuge-h22 | **3/3** | **4/4** | 2/4 | 0/1 | 0/1 | 0/1 | 0/1 |
|  | Centrifuge-h500 | **3/3** | **4/4** | 2/4 | 0/1 | 0/1 | 0/1 | 0/1 |
|  | MetaPhlAn3 | **3/3** | **4/4** | 2/4 | 0/1 | 0/1 | 0/1 | 0/1 |
|  | mOTUs | **3/3** | 1/4 | 0/4 | 0/1 | 0/1 | 0/1 | 0/1 |
|  | Sourmash-k31 | **3/3** | **4/4** | **4/4** | 0/1 | 0/1 | 0/1 | 0/1 |
|  | Sourmash-k51 | **3/3** | **4/4** | **4/4** | 0/1 | 0/1 | 0/1 | 0/1 |
|  | MetaMaps | 2/3 | **4/4** | 3/4 | 0/1 | 0/1 | 0/1 | 0/1 |
|  | MMseqs2 | **3/3** | **4/4** | **3/4** | 0/1 | 0/1 | 0/1 | 0/1 |
|  | MEGAN-LR-prot | **3/3** | **4/4** | **4/4** | 0/1 | 0/1 | 0/1 | 0/1 |
|  | MEGAN-LR-nuc-HiFi | **3/3** | **4/4** | 3/4 | 0/1 | 0/1 | 0/1 | 0/1 |
|  | MEGAN-LR-nuc-ONT | **3/3** | **4/4** | 3/4 | 0/1 | 0/1 | 0/1 | 0/1 |
|  | BugSeq-V2 | **3/3** | **4/4** | **4/4** | 0/1 | 0/1 | 0/1 | 0/1 |

Perfect detection scores are highlighted in bold. For Zymo D6331 two species were excluded at the 14% abundance level, which were missing from the databases for several methods.

**Supplementary Table S13.** Genus-level detection results based on the minimum 0.1% of total reads threshold.

| **Dataset** | **Method Type** | **Profiling Method** | **True Positives** | **False Positives** | **False Negatives** | **Precision** | **Recall** | **F1** | **F0.5** |
| --- | --- | --- | --- | --- | --- | --- | --- | --- | --- |
| HiFi ATCC-MSA1003 | Short read | Kraken2 | 11 | 3 | 7 | 0.79 | 0.61 | 0.69 | 0.74 |
| (18 genera, staggered) |  | Bracken | 11 | 3 | 7 | 0.79 | 0.61 | 0.69 | 0.74 |
|  |  | Centrifuge-h22 | 11 | 0 | 7 | 1.00 | 0.61 | 0.76 | 0.89 |
|  |  | Centrifuge-h500 | 11 | 0 | 7 | 1.00 | 0.61 | 0.76 | 0.89 |
|  |  | Metaphlan3 | 7 | 2 | 11 | 0.78 | 0.39 | 0.52 | 0.65 |
|  |  | mOTUs | 4 | 0 | 14 | 1.00 | 0.22 | 0.36 | 0.59 |
|  | General | Sourmash-k31 | 18 | 2 | 0 | 0.90 | 1.00 | 0.95 | 0.92 |
|  |  | Sourmash-k51 | 18 | 1 | 0 | 0.95 | 1.00 | 0.97 | 0.96 |
|  | Long read | Metamaps | 11 | 0 | 7 | 1.00 | 0.61 | 0.76 | 0.89 |
|  |  | MMseqs2 | 10 | 0 | 8 | 1.00 | 0.56 | 0.71 | 0.86 |
|  |  | MEGAN-LR-Prot | 9 | 0 | 9 | 1.00 | 0.50 | 0.67 | 0.83 |
|  |  | MEGAN-LR-Nuc-HiFi | 11 | 0 | 7 | 1.00 | 0.61 | 0.76 | 0.89 |
|  |  | MEGAN-LR-Nuc-ONT | 11 | 0 | 7 | 1.00 | 0.61 | 0.76 | 0.89 |
|  |  | BugSeq-V2 | 11 | 0 | 7 | 1.00 | 0.61 | 0.76 | 0.89 |
| HiFi Zymo-D6331 | Short read | Kraken2 | 13 | 5 | 4 | 0.72 | 0.77 | 0.74 | 0.73 |
| (17 genera, staggered) |  | Bracken | 13 | 5 | 4 | 0.72 | 0.77 | 0.74 | 0.73 |
|  |  | Centrifuge-h22 | 11 | 5 | 6 | 0.69 | 0.65 | 0.67 | 0.68 |
|  |  | Centrifuge-h500 | 11 | 2 | 6 | 0.85 | 0.65 | 0.73 | 0.80 |
|  |  | Metaphlan3 | 11 | 1 | 6 | 0.92 | 0.65 | 0.76 | 0.85 |
|  |  | mOTUs | 0 | 0 | 17 | 0.00 | 0.00 | 0.00 | 0.00 |
|  | General | Sourmash-k31 | 16 | 0 | 1 | 1.00 | 0.94 | 0.97 | 0.99 |
|  |  | Sourmash-k51 | 16 | 1 | 1 | 0.94 | 0.94 | 0.94 | 0.94 |
|  | Long read | Metamaps | 12 | 3 | 5 | 0.80 | 0.71 | 0.75 | 0.78 |
|  |  | MMseqs2 | 13 | 0 | 4 | 1.00 | 0.77 | 0.87 | 0.94 |
|  |  | MEGAN-LR-Prot | 13 | 0 | 4 | 1.00 | 0.77 | 0.87 | 0.94 |
|  |  | MEGAN-LR-Nuc-HiFi | 13 | 0 | 4 | 1.00 | 0.77 | 0.87 | 0.94 |
|  |  | MEGAN-LR-Nuc-ONT | 13 | 0 | 4 | 1.00 | 0.77 | 0.87 | 0.94 |
|  |  | BugSeq-V2 | 13 | 0 | 4 | 1.00 | 0.77 | 0.87 | 0.94 |
| ONT R10 Zymo-D6300 | Short read | Kraken2 | 10 | 0 | 0 | 1.00 | 1.00 | 1.00 | 1.00 |
| (10 genera, even) |  | Bracken | 10 | 0 | 0 | 1.00 | 1.00 | 1.00 | 1.00 |
|  |  | Centrifuge-h22 | 8 | 0 | 2 | 1.00 | 0.80 | 0.89 | 0.95 |
|  |  | Centrifuge-h500 | 5 | 0 | 5 | 1.00 | 0.50 | 0.67 | 0.83 |
|  |  | Metaphlan3 | 9 | 1 | 1 | 0.90 | 0.90 | 0.90 | 0.90 |
|  |  | mOTUs | 0 | 0 | 10 | 0.00 | 0.00 | 0.00 | 0.00 |
|  | General | Sourmash-k31 | 10 | 2 | 0 | 0.83 | 1.00 | 0.91 | 0.86 |
|  |  | Sourmash-k51 | 10 | 1 | 0 | 0.91 | 1.00 | 0.95 | 0.93 |
|  | Long read | Metamaps | 9 | 0 | 1 | 1.00 | 0.90 | 0.95 | 0.98 |
|  |  | MMseqs2 | 10 | 0 | 0 | 1.00 | 1.00 | 1.00 | 1.00 |
|  |  | MEGAN-LR-Prot | 10 | 0 | 0 | 1.00 | 1.00 | 1.00 | 1.00 |
|  |  | MEGAN-LR-Nuc-HiFi | 10 | 0 | 0 | 1.00 | 1.00 | 1.00 | 1.00 |
|  |  | MEGAN-LR-Nuc-ONT | 10 | 0 | 0 | 1.00 | 1.00 | 1.00 | 1.00 |
|  |  | BugSeq-V2 | 10 | 0 | 0 | 1.00 | 1.00 | 1.00 | 1.00 |
| ONT Q20 Zymo-D6300 | Short read | Kraken2 | 10 | 3 | 0 | 0.77 | 1.00 | 0.87 | 0.81 |
| (10 genera, even) |  | Bracken | 10 | 3 | 0 | 0.77 | 1.00 | 0.87 | 0.81 |
|  |  | Centrifuge-h22 | 8 | 0 | 2 | 1.00 | 0.80 | 0.89 | 0.95 |
|  |  | Centrifuge-h500 | 8 | 0 | 2 | 1.00 | 0.80 | 0.89 | 0.95 |
|  |  | Metaphlan3 | 9 | 3 | 1 | 0.75 | 0.90 | 0.82 | 0.78 |
|  |  | mOTUs | 0 | 0 | 10 | 0.00 | 0.00 | 0.00 | 0.00 |
|  | General | Sourmash-k31 | 10 | 1 | 0 | 0.91 | 1.00 | 0.95 | 0.93 |
|  |  | Sourmash-k51 | 10 | 1 | 0 | 0.91 | 1.00 | 0.95 | 0.93 |
|  | Long read | Metamaps | 9 | 1 | 1 | 0.90 | 0.90 | 0.90 | 0.90 |
|  |  | MMseqs2 | 10 | 0 | 0 | 1.00 | 1.00 | 1.00 | 1.00 |
|  |  | MEGAN-LR-Prot | 10 | 0 | 0 | 1.00 | 1.00 | 1.00 | 1.00 |
|  |  | MEGAN-LR-Nuc-HiFi | 10 | 0 | 0 | 1.00 | 1.00 | 1.00 | 1.00 |
|  |  | MEGAN-LR-Nuc-ONT | 10 | 0 | 0 | 1.00 | 1.00 | 1.00 | 1.00 |
|  |  | BugSeq-V2 | 10 | 0 | 0 | 1.00 | 1.00 | 1.00 | 1.00 |

**Supplementary Table S14.** Species-level detection results based on the minimum 1% of total reads threshold.

| **Dataset** | **Method Type** | **Profiling Method** | **True Positives** | **False Positives** | **False Negatives** | **Precision** | **Recall** | **F1** | **F0.5** |
| --- | --- | --- | --- | --- | --- | --- | --- | --- | --- |
| HiFi ATCC-MSA1003 | Short read | Kraken2 | 8 | 0 | 12 | 1.00 | 0.40 | 0.57 | 0.77 |
| (20 species, staggered) |  | Bracken | 8 | 0 | 12 | 1.00 | 0.40 | 0.57 | 0.77 |
|  |  | Centrifuge-h22 | 8 | 0 | 12 | 1.00 | 0.40 | 0.57 | 0.77 |
|  |  | Centrifuge-h500 | 8 | 0 | 12 | 1.00 | 0.40 | 0.57 | 0.77 |
|  |  | Metaphlan3 | 5 | 0 | 15 | 1.00 | 0.25 | 0.40 | 0.63 |
|  |  | mOTUs | 0 | 0 | 20 | 0.00 | 0.00 | 0.00 | 0.00 |
|  | General | Sourmash-k31 | 20 | 5 | 0 | 0.80 | 1.00 | 0.89 | 0.83 |
|  |  | Sourmash-k51 | 20 | 3 | 0 | 0.87 | 1.00 | 0.93 | 0.89 |
|  | Long read | Metamaps | 8 | 0 | 12 | 1.00 | 0.40 | 0.57 | 0.77 |
|  |  | MMseqs2 | 5 | 0 | 15 | 1.00 | 0.25 | 0.40 | 0.63 |
|  |  | MEGAN-LR-Prot | 6 | 0 | 14 | 1.00 | 0.30 | 0.46 | 0.68 |
|  |  | MEGAN-LR-Nuc-HiFi | 7 | 0 | 13 | 1.00 | 0.35 | 0.52 | 0.73 |
|  |  | MEGAN-LR-Nuc-ONT | 6 | 0 | 14 | 1.00 | 0.30 | 0.46 | 0.68 |
|  |  | BugSeq-V2 | 7 | 0 | 13 | 1.00 | 0.35 | 0.52 | 0.73 |
| HiFi Zymo-D6331 | Short read | Kraken2* | 9 | 0 | 6 | 1.00 | 0.60 | 0.75 | 0.88 |
| (17 species, staggered) |  | Bracken* | 9 | 0 | 6 | 1.00 | 0.60 | 0.75 | 0.88 |
| *based on 15 species |  | Centrifuge-h22* | 9 | 0 | 6 | 1.00 | 0.60 | 0.75 | 0.88 |
|  |  | Centrifuge-h500* | 9 | 0 | 6 | 1.00 | 0.60 | 0.75 | 0.88 |
|  |  | Metaphlan3* | 5 | 0 | 10 | 1.00 | 0.33 | 0.50 | 0.71 |
|  |  | mOTUs | 0 | 0 | 15 | 0.00 | 0.00 | 0.00 | 0.00 |
|  | General | Sourmash-k31 | 14 | 1 | 1 | 0.93 | 0.93 | 0.93 | 0.93 |
|  |  | Sourmash-k51 | 14 | 2 | 1 | 0.88 | 0.93 | 0.90 | 0.89 |
|  | Long read | Metamaps* | 8 | 0 | 7 | 1.00 | 0.53 | 0.70 | 0.85 |
|  |  | MMseqs2* | 4 | 0 | 11 | 1.00 | 0.27 | 0.42 | 0.65 |
|  |  | MEGAN-LR-Prot* | 6 | 0 | 9 | 1.00 | 0.40 | 0.57 | 0.77 |
|  |  | MEGAN-LR-Nuc-HiFi* | 8 | 0 | 7 | 1.00 | 0.53 | 0.70 | 0.85 |
|  |  | MEGAN-LR-Nuc-ONT* | 8 | 0 | 7 | 1.00 | 0.53 | 0.70 | 0.85 |
|  |  | BugSeq-V2* | 8 | 0 | 7 | 1.00 | 0.53 | 0.70 | 0.85 |
| ONT R10 Zymo-D6300 | Short read | Kraken2 | 10 | 1 | 0 | 0.91 | 1.00 | 0.95 | 0.93 |
| (10 species, even) |  | Bracken | 10 | 1 | 0 | 0.91 | 1.00 | 0.95 | 0.93 |
|  |  | Centrifuge-h22 | 8 | 0 | 2 | 1.00 | 0.80 | 0.89 | 0.95 |
|  |  | Centrifuge-h500 | 0 | 0 | 10 | 0.00 | 0.00 | 0.00 | 0.00 |
|  |  | Metaphlan3 | 6 | 1 | 4 | 0.86 | 0.60 | 0.71 | 0.79 |
|  |  | mOTUs | 0 | 0 | 10 | 0.00 | 0.00 | 0.00 | 0.00 |
|  | General | Sourmash-k31 | 10 | 11 | 0 | 0.48 | 1.00 | 0.64 | 0.53 |
|  |  | Sourmash-k51 | 10 | 3 | 0 | 0.77 | 1.00 | 0.87 | 0.81 |
|  | Long read | Metamaps | 9 | 0 | 1 | 1.00 | 0.90 | 0.95 | 0.98 |
|  |  | MMseqs2 | 0 | 0 | 10 | 0.00 | 0.00 | 0.00 | 0.00 |
|  |  | MEGAN-LR-Prot | 6 | 0 | 4 | 1.00 | 0.60 | 0.75 | 0.88 |
|  |  | MEGAN-LR-Nuc-HiFi | 7 | 0 | 3 | 1.00 | 0.70 | 0.82 | 0.92 |
|  |  | MEGAN-LR-Nuc-ONT | 7 | 0 | 3 | 1.00 | 0.70 | 0.82 | 0.92 |
|  |  | BugSeq-V2 | 8 | 0 | 2 | 1.00 | 0.80 | 0.89 | 0.95 |
| ONT Q20 Zymo-D6300 | Short read | Kraken2 | 10 | 1 | 0 | 0.91 | 1.00 | 0.95 | 0.93 |
| (10 species, even) |  | Bracken | 10 | 1 | 0 | 0.91 | 1.00 | 0.95 | 0.93 |
|  |  | Centrifuge-h22 | 8 | 0 | 2 | 1.00 | 0.80 | 0.89 | 0.95 |
|  |  | Centrifuge-h500 | 7 | 0 | 3 | 1.00 | 0.70 | 0.82 | 0.92 |
|  |  | Metaphlan3 | 6 | 1 | 4 | 0.86 | 0.60 | 0.71 | 0.79 |
|  |  | mOTUs | 0 | 0 | 10 | 0.00 | 0.00 | 0.00 | 0.00 |
|  | General | Sourmash-k31 | 10 | 4 | 0 | 0.71 | 1.00 | 0.83 | 0.76 |
|  |  | Sourmash-k51 | 10 | 4 | 0 | 0.71 | 1.00 | 0.83 | 0.76 |
|  | Long read | Metamaps | 9 | 1 | 1 | 0.90 | 0.90 | 0.90 | 0.90 |
|  |  | MMseqs2 | 4 | 0 | 6 | 1.00 | 0.40 | 0.57 | 0.77 |
|  |  | MEGAN-LR-Prot | 8 | 0 | 2 | 1.00 | 0.80 | 0.89 | 0.95 |
|  |  | MEGAN-LR-Nuc-HiFi | 7 | 0 | 3 | 1.00 | 0.70 | 0.82 | 0.92 |
|  |  | MEGAN-LR-Nuc-ONT | 7 | 0 | 3 | 1.00 | 0.70 | 0.82 | 0.92 |
|  |  | BugSeq-V2 | 10 | 0 | 0 | 1.00 | 1.00 | 1.00 | 1.00 |

**Supplementary Table S15.** The number of species detected at each relative abundance level in the HiFi ATCC MSA-1003 and HiFi Zymo D6331 datasets, based on the minimum 1% of total reads threshold.

|  |  | **Species Detected at Relative Abundance Level** | | | | | | |
| --- | --- | --- | --- | --- | --- | --- | --- | --- |
| **Dataset** | **Method** | **14-18%** | **6%** | **1.5-1.8%** | **0.10-0.18%** | **0.01-0.02%** | **0.001%** | **0.0001%** |
| HiFi ATCC MSA-1003 | Kraken2 | **5/5** | - | 3/5 | 0/5 | 0/5 | - | - |
|  | Bracken | **5/5** | - | 3/5 | 0/5 | 0/5 | - | - |
|  | Centrifuge-h22 | **5/5** | - | 3/5 | 0/5 | 0/5 | - | - |
|  | Centrifuge-h500 | **5/5** | - | 3/5 | 0/5 | 0/5 | - | - |
|  | MetaPhlAn3 | **5/5** | - | 0/5 | 0/5 | 0/5 | - | - |
|  | mOTUs | 0/5 | - | 0/5 | 0/5 | 0/5 | - | - |
|  | Sourmash-k31 | **5/5** | - | 2/5 | 0/5 | 0/5 | - | - |
|  | Sourmash-k51 | **5/5** | - | 2/5 | 0/5 | 0/5 | - | - |
|  | MetaMaps | **5/5** | - | 3/5 | 0/5 | 0/5 | - | - |
|  | MMseqs2 | 4/5 | - | 1/5 | 0/5 | 0/5 | - | - |
|  | MEGAN-LR-prot | **5/5** | - | 1/5 | 0/5 | 0/5 | - | - |
|  | MEGAN-LR-nuc-HiFi | **5/5** | - | 2/5 | 0/5 | 0/5 | - | - |
|  | MEGAN-LR-nuc-ONT | **5/5** | - | 1/5 | 0/5 | 0/5 | - | - |
|  | BugSeq-V2 | **5/5** | - | 2/5 | 0/5 | 0/5 | - | - |
| HiFi Zymo D6331 | Kraken2 | **3/3** | **4/4** | 2/4 | 0/1 | 0/1 | 0/1 | 0/1 |
|  | Bracken | **3/3** | **4/4** | 2/4 | 0/1 | 0/1 | 0/1 | 0/1 |
|  | Centrifuge-h22 | **3/3** | **4/4** | 2/4 | 0/1 | 0/1 | 0/1 | 0/1 |
|  | Centrifuge-h500 | **3/3** | **4/4** | 2/4 | 0/1 | 0/1 | 0/1 | 0/1 |
|  | MetaPhlAn3 | **3/3** | 2/4 | 0/4 | 0/1 | 0/1 | 0/1 | 0/1 |
|  | mOTUs | 0/3 | 0/4 | 0/4 | 0/1 | 0/1 | 0/1 | 0/1 |
|  | Sourmash-k31 | **3/3** | **4/4** | 2/4 | 0/1 | 0/1 | 0/1 | 0/1 |
|  | Sourmash-k51 | **3/3** | **4/4** | 2/4 | 0/1 | 0/1 | 0/1 | 0/1 |
|  | MetaMaps | 2/3 | **4/4** | 2/4 | 0/1 | 0/1 | 0/1 | 0/1 |
|  | MMseqs2 | 1/3 | 2/4 | 1/4 | 0/1 | 0/1 | 0/1 | 0/1 |
|  | MEGAN-LR-prot | **3/3** | 1/4 | 1/4 | 0/1 | 0/1 | 0/1 | 0/1 |
|  | MEGAN-LR-nuc-HiFi | **3/3** | 3/4 | 2/4 | 0/1 | 0/1 | 0/1 | 0/1 |
|  | MEGAN-LR-nuc-ONT | **3/3** | 3/4 | 2/4 | 0/1 | 0/1 | 0/1 | 0/1 |
|  | BugSeq-V2 | **3/3** | 3/4 | 2/4 | 0/1 | 0/1 | 0/1 | 0/1 |

Perfect detection scores are highlighted in bold. For Zymo D6331 two species were excluded at the 14% abundance level, which were missing from the databases for several methods.

**Supplementary Table S16.** Genus-level detection results based on the minimum 1% of total reads threshold.

| **Dataset** | **Method Type** | **Profiling Method** | **True Positives** | **False Positives** | **False Negatives** | **Precision** | **Recall** | **F1** | **F0.5** |
| --- | --- | --- | --- | --- | --- | --- | --- | --- | --- |
| HiFi ATCC-MSA1003 | Short read | Kraken2 | 8 | 0 | 10 | 1.00 | 0.44 | 0.61 | 0.80 |
| (18 genera, staggered) |  | Bracken | 8 | 1 | 10 | 0.89 | 0.44 | 0.59 | 0.74 |
|  |  | Centrifuge-h22 | 8 | 0 | 10 | 1.00 | 0.44 | 0.61 | 0.80 |
|  |  | Centrifuge-h500 | 8 | 0 | 10 | 1.00 | 0.44 | 0.61 | 0.80 |
|  |  | Metaphlan3 | 5 | 0 | 13 | 1.00 | 0.28 | 0.44 | 0.66 |
|  |  | mOTUs | 0 | 0 | 18 | 0.00 | 0.00 | 0.00 | 0.00 |
|  | General | Sourmash-k31 | 18 | 2 | 0 | 0.90 | 1.00 | 0.95 | 0.92 |
|  |  | Sourmash-k51 | 18 | 1 | 0 | 0.95 | 1.00 | 0.97 | 0.96 |
|  | Long read | Metamaps | 8 | 0 | 10 | 1.00 | 0.44 | 0.61 | 0.80 |
|  |  | MMseqs2 | 8 | 0 | 10 | 1.00 | 0.44 | 0.61 | 0.80 |
|  |  | MEGAN-LR-Prot | 8 | 0 | 10 | 1.00 | 0.44 | 0.61 | 0.80 |
|  |  | MEGAN-LR-Nuc-HiFi | 8 | 0 | 10 | 1.00 | 0.44 | 0.61 | 0.80 |
|  |  | MEGAN-LR-Nuc-ONT | 8 | 0 | 10 | 1.00 | 0.44 | 0.61 | 0.80 |
|  |  | BugSeq-V2 | 7 | 0 | 11 | 1.00 | 0.39 | 0.56 | 0.76 |
| HiFi Zymo-D6331 | Short read | Kraken2 | 11 | 0 | 6 | 1.00 | 0.65 | 0.79 | 0.90 |
| (17 genera, staggered) |  | Bracken | 11 | 0 | 6 | 1.00 | 0.65 | 0.79 | 0.90 |
|  |  | Centrifuge-h22 | 11 | 0 | 6 | 1.00 | 0.65 | 0.79 | 0.90 |
|  |  | Centrifuge-h500 | 10 | 0 | 7 | 1.00 | 0.59 | 0.74 | 0.88 |
|  |  | Metaphlan3 | 7 | 0 | 10 | 1.00 | 0.41 | 0.58 | 0.78 |
|  |  | mOTUs | 0 | 0 | 17 | 0.00 | 0.00 | 0.00 | 0.00 |
|  | General | Sourmash-k31 | 16 | 0 | 1 | 1.00 | 0.94 | 0.97 | 0.99 |
|  |  | Sourmash-k51 | 16 | 1 | 1 | 0.94 | 0.94 | 0.94 | 0.94 |
|  | Long read | Metamaps | 10 | 0 | 7 | 1.00 | 0.59 | 0.74 | 0.88 |
|  |  | MMseqs2 | 10 | 0 | 7 | 1.00 | 0.59 | 0.74 | 0.88 |
|  |  | MEGAN-LR-Prot | 10 | 0 | 7 | 1.00 | 0.59 | 0.74 | 0.88 |
|  |  | MEGAN-LR-Nuc-HiFi | 11 | 0 | 6 | 1.00 | 0.65 | 0.79 | 0.90 |
|  |  | MEGAN-LR-Nuc-ONT | 11 | 0 | 6 | 1.00 | 0.65 | 0.79 | 0.90 |
|  |  | BugSeq-V2 | 10 | 0 | 7 | 1.00 | 0.59 | 0.74 | 0.88 |
| ONT R10 Zymo-D6300 | Short read | Kraken2 | 10 | 0 | 0 | 1.00 | 1.00 | 1.00 | 1.00 |
| (10 genera, even) |  | Bracken | 10 | 0 | 0 | 1.00 | 1.00 | 1.00 | 1.00 |
|  |  | Centrifuge-h22 | 8 | 0 | 2 | 1.00 | 0.80 | 0.89 | 0.95 |
|  |  | Centrifuge-h500 | 0 | 0 | 10 | 0.00 | 0.00 | 0.00 | 0.00 |
|  |  | Metaphlan3 | 7 | 0 | 3 | 1.00 | 0.70 | 0.82 | 0.92 |
|  |  | mOTUs | 0 | 0 | 10 | 0.00 | 0.00 | 0.00 | 0.00 |
|  | General | Sourmash-k31 | 10 | 2 | 0 | 0.83 | 1.00 | 0.91 | 0.86 |
|  |  | Sourmash-k51 | 10 | 1 | 0 | 0.91 | 1.00 | 0.95 | 0.93 |
|  | Long read | Metamaps | 9 | 0 | 1 | 1.00 | 0.90 | 0.95 | 0.98 |
|  |  | MMseqs2 | 5 | 0 | 5 | 1.00 | 0.50 | 0.67 | 0.83 |
|  |  | MEGAN-LR-Prot | 7 | 0 | 3 | 1.00 | 0.70 | 0.82 | 0.92 |
|  |  | MEGAN-LR-Nuc-HiFi | 7 | 0 | 3 | 1.00 | 0.70 | 0.82 | 0.92 |
|  |  | MEGAN-LR-Nuc-ONT | 7 | 0 | 3 | 1.00 | 0.70 | 0.82 | 0.92 |
|  |  | BugSeq-V2 | 9 | 0 | 1 | 1.00 | 0.90 | 0.95 | 0.98 |
| ONT Q20 Zymo-D6300 | Short read | Kraken2 | 10 | 0 | 0 | 1.00 | 1.00 | 1.00 | 1.00 |
| (10 genera, even) |  | Bracken | 10 | 0 | 0 | 1.00 | 1.00 | 1.00 | 1.00 |
|  |  | Centrifuge-h22 | 8 | 0 | 2 | 1.00 | 0.80 | 0.89 | 0.95 |
|  |  | Centrifuge-h500 | 8 | 0 | 2 | 1.00 | 0.80 | 0.89 | 0.95 |
|  |  | Metaphlan3 | 7 | 0 | 3 | 1.00 | 0.70 | 0.82 | 0.92 |
|  |  | mOTUs | 0 | 0 | 10 | 0.00 | 0.00 | 0.00 | 0.00 |
|  | General | Sourmash-k31 | 10 | 1 | 0 | 0.91 | 1.00 | 0.95 | 0.93 |
|  |  | Sourmash-k51 | 10 | 1 | 0 | 0.91 | 1.00 | 0.95 | 0.93 |
|  | Long read | Metamaps | 9 | 1 | 1 | 0.90 | 0.90 | 0.90 | 0.90 |
|  |  | MMseqs2 | 8 | 0 | 2 | 1.00 | 0.80 | 0.89 | 0.95 |
|  |  | MEGAN-LR-Prot | 9 | 0 | 1 | 1.00 | 0.90 | 0.95 | 0.98 |
|  |  | MEGAN-LR-Nuc-HiFi | 7 | 0 | 3 | 1.00 | 0.70 | 0.82 | 0.92 |
|  |  | MEGAN-LR-Nuc-ONT | 7 | 0 | 3 | 1.00 | 0.70 | 0.82 | 0.92 |
|  |  | BugSeq-V2 | 10 | 0 | 0 | 1.00 | 1.00 | 1.00 | 1.00 |

**Supplementary Table S17.** Chi-Squared test results for the species level relative abundances.

|  | **HiFi ATCC MSA-1003** | | | | **HiFi Zymo D6331** | | | | **ONT R10 Zymo D6300** | | | | **ONT Q20 Zymo D6300** | | | |
| --- | --- | --- | --- | --- | --- | --- | --- | --- | --- | --- | --- | --- | --- | --- | --- | --- |
| **Method** | **df** | **Chi-Squared Statistic** | **Critical Value** | **P-value** | **df** | **Chi-Squared Statistic** | **Critical Value** | **P-value** | **df** | **Chi-Squared Statistic** | **Critical Value** | **P-value** | **df** | **Chi-Squared Statistic** | **Critical Value** | **P-value** |
| Kraken2 | 20 | 519.96 | 31.41 | **<0.001** | 17 | 44411.1 | 27.59 | **<0.001** | 10 | 1037.46 | 18.31 | **<0.001** | 10 | 2159.47 | 18.31 | **<0.001** |
| Bracken | 20 | 692.00 | 31.41 | **<0.001** | 17 | 43996.82 | 27.59 | **<0.001** | 10 | 1124.69 | 18.31 | **<0.001** | 10 | 3104.71 | 18.31 | **<0.001** |
| Centrifuge-h22 | 20 | 76.98 | 31.41 | **<0.001** | 17 | 45187.2 | 27.59 | **<0.001** | 10 | 116.53 | 18.31 | **<0.001** | 10 | 73.48 | 18.31 | **<0.001** |
| Centrifuge-h500 | 20 | 77.29 | 31.41 | **<0.001** | 17 | 1549.28 | 27.59 | **<0.001** | 10 | 145.87 | 18.31 | **<0.001** | 10 | 35.26 | 18.31 | **<0.001** |
| Metaphlan3 | 20 | 1379.35 | 31.41 | **<0.001** | 17 | 22021.22 | 27.59 | **<0.001** | 10 | 28723.48 | 18.31 | **<0.001** | 10 | 39381.31 | 18.31 | **<0.001** |
| mOTUs | 20 | 821.56 | 31.41 | **<0.001** | 17 | 282.93 | 27.59 | **<0.001** | 10 | 7.8 | 18.31 | 0.648 | 10 | 93.11 | 18.31 | **<0.001** |
| Sourmash-k31 | 20 | 7789.88 | 31.41 | **<0.001** | 17 | 31383.24 | 27.59 | **<0.001** | 10 | 52.84 | 18.31 | **<0.001** | 10 | 51.33 | 18.31 | **<0.001** |
| Sourmash-k51 | 20 | 7148.22 | 31.41 | **<0.001** | 17 | 32967.88 | 27.59 | **<0.001** | 10 | 14.22 | 18.31 | 0.163 | 10 | 34.72 | 18.31 | **<0.001** |
| Metamaps | 20 | 34.31 | 31.41 | 0.024 | 17 | 41737.33 | 27.59 | **<0.001** | 10 | 34.75 | 18.31 | **<0.001** | 10 | 296.81 | 18.31 | **<0.001** |
| MMseqs2 | 20 | 38.07 | 31.41 | 0.009 | 17 | 1374.97 | 27.59 | **<0.001** | 10 | 2014.14 | 18.31 | **<0.001** | 10 | 171.75 | 18.31 | **<0.001** |
| MEGAN-LR-Prot | 20 | 43.35 | 31.41 | 0.005 | 17 | 128.73 | 27.59 | **<0.001** | 10 | 58.86 | 18.31 | **<0.001** | 10 | 17.23 | 18.31 | 0.069 |
| MEGAN-LR-Nuc-HiFi | 20 | 66.10 | 31.41 | **<0.001** | 17 | 11752.56 | 27.59 | **<0.001** | 10 | 82.72 | 18.31 | **<0.001** | 10 | 66.88 | 18.31 | **<0.001** |
| MEGAN-LR-Nuc-ONT | 20 | 67.23 | 31.41 | **<0.001** | 17 | 11841.98 | 27.59 | **<0.001** | 10 | 83.22 | 18.31 | **<0.001** | 10 | 67.45 | 18.31 | **<0.001** |
| BugSeq-V2 | 20 | 26.30 | 31.41 | 0.156 | 17 | 66.41 | 27.59 | **<0.001** | 10 | 5.82 | 18.31 | 0.830 | 10 | 1.81 | 18.31 | 0.998 |

The Bonferroni correction for 11 tests sets the alpha to 0.0045, and tests that are significant after the correction are shown in bold. P-values in bold indicate rejection of the null hypothesis that there is no difference between the distributions (signifying the abundance estimates are significantly different).

**Supplementary Table S18.** Chi-Squared test results for the genus level relative abundances.

|  | **HiFi ATCC MSA-1003** | | | | **HiFi Zymo D6331** | | | | **ONT R10 Zymo D6300** | | | | **ONT Q20 Zymo D6300** | | | |
| --- | --- | --- | --- | --- | --- | --- | --- | --- | --- | --- | --- | --- | --- | --- | --- | --- |
| **Method** | **df** | **Chi-Squared Statistic** | **Critical Value** | **P-value** | **df** | **Chi-Squared Statistic** | **Critical Value** | **P-value** | **df** | **Chi-Squared Statistic** | **Critical Value** | **P-value** | **df** | **Chi-Squared Statistic** | **Critical Value** | **P-value** |
| Kraken2 | 18 | 366.01 | 28.87 | **<0.001** | 17 | 483.13 | 27.59 | **<0.001** | 10 | 12.20 | 18.31 | 0.272 | 10 | 119.27 | 18.31 | **<0.001** |
| Bracken | 18 | 367.40 | 28.87 | **<0.001** | 17 | 457.86 | 27.59 | **<0.001** | 10 | 8.77 | 18.31 | 0.554 | 10 | 103.95 | 18.31 | **<0.001** |
| Centrifuge-h22 | 18 | 34.24 | 28.87 | 0.012 | 17 | 810.43 | 27.59 | **<0.001** | 10 | 18.11 | 18.31 | 0.053 | 10 | 11.53 | 18.31 | 0.318 |
| Centrifuge-h500 | 18 | 35.21 | 28.87 | 0.009 | 17 | 185.49 | 27.59 | **<0.001** | 10 | 56.50 | 18.31 | **<0.001** | 10 | 9.64 | 18.31 | 0.473 |
| Metaphlan3 | 18 | 862.31 | 28.87 | **<0.001** | 17 | 352.98 | 27.59 | **<0.001** | 10 | 393.30 | 18.31 | **<0.001** | 10 | 1056.36 | 18.31 | **<0.001** |
| mOTUs | 18 | 31.3 | 28.87 | 0.028 | 17 | 52.55 | 27.59 | **<0.001** | 10 | 7.8 | 18.31 | 0.648 | 10 | 92.95 | 18.31 | **<0.001** |
| Sourmash-k31 | 18 | 31.2 | 28.87 | 0.027 | 17 | 24.04 | 27.59 | 0.118 | 10 | 50.81 | 18.31 | **<0.001** | 10 | 49.21 | 18.31 | **<0.001** |
| Sourmash-k51 | 18 | 29.34 | 28.87 | 0.044 | 17 | 24.96 | 27.59 | 0.096 | 10 | 13.94 | 18.31 | 0.176 | 10 | 33.58 | 18.31 | **<0.001** |
| Metamaps | 18 | 33.93 | 28.87 | 0.013 | 17 | 247.08 | 27.59 | **<0.001** | 10 | 11.94 | 18.31 | 0.289 | 10 | 189.71 | 18.31 | **<0.001** |
| MMseqs2 | 18 | 24.54 | 28.87 | 0.138 | 17 | 55.32 | 27.59 | **<0.001** | 10 | 170.75 | 18.31 | **<0.001** | 10 | 65.06 | 18.31 | **<0.001** |
| MEGAN-LR-Prot | 18 | 24.51 | 28.87 | 0.139 | 17 | 42.63 | 27.59 | **0.001** | 10 | 61.89 | 18.31 | **<0.001** | 10 | 15.84 | 18.31 | 0.104 |
| MEGAN-LR-Nuc-HiFi | 18 | 59.77 | 28.87 | **<0.001** | 17 | 46.79 | 27.59 | **<0.001** | 10 | 79.61 | 18.31 | **<0.001** | 10 | 65.33 | 18.31 | **<0.001** |
| MEGAN-LR-Nuc-ONT | 18 | 59.82 | 28.87 | **<0.001** | 17 | 43.66 | 27.59 | **<0.001** | 10 | 79.73 | 18.31 | **<0.001** | 10 | 65.55 | 18.31 | **<0.001** |
| BugSeq-V2 | 18 | 26.03 | 28.87 | 0.099 | 17 | 26.52 | 27.59 | 0.065 | 10 | 5.53 | 18.31 | 0.853 | 10 | 1.53 | 18.31 | 0.999 |

The Bonferroni correction for 11 tests sets the alpha to 0.0045, and tests that are significant after the correction are shown in bold. P-values in bold indicate rejection of the null hypothesis that there is no difference between the distributions (signifying the abundance estimates are significantly different).

**Supplementary Table S19.** Species-level detection results for the ONT Short datasets.

| **Dataset** | **Filtering Level** | **Method Type** | **Method** | **True Positives** | **False Positives** | **False Negatives** | **Precision** | **Recall** | **F_1_** | **F_0.5_** | **L1** |
| --- | --- | --- | --- | --- | --- | --- | --- | --- | --- | --- | --- |
| ONT R10 Short | 0.001 percent | Short read | Kraken2 | 10 | 121 | 0 | 0.08 | 1.00 | 0.14 | 0.09 | 118.8 |
|  |  |  | Bracken | 10 | 125 | 0 | 0.07 | 1.00 | 0.14 | 0.09 | 115.9 |
|  |  |  | Centrifuge-h22 | 8 | 26 | 2 | 0.24 | 0.80 | 0.36 | 0.27 | 34.9 |
|  |  |  | Centrifuge-h500 | 7 | 0 | 3 | 1.00 | 0.70 | 0.82 | 0.92 | 79.7 |
|  |  |  | Metaphlan3 | 8 | 20 | 2 | 0.29 | 0.80 | 0.42 | 0.33 | 119.8 |
|  |  | Long read | Metamaps | 9 | 29 | 1 | 0.24 | 0.90 | 0.38 | 0.28 | 51.8 |
|  |  |  | MMseqs2 | 10 | 87 | 0 | 0.10 | 1.00 | 0.19 | 0.13 | 124.1 |
|  |  |  | MEGAN-LR-Prot | 10 | 7 | 0 | 0.59 | 1.00 | 0.74 | 0.64 | 136.2 |
|  |  |  | MEGAN-LR-Nuc-HiFi | 10 | 1 | 0 | 0.91 | 1.00 | 0.95 | 0.93 | 138.9 |
|  |  |  | MEGAN-LR-Nuc-ONT | 10 | 1 | 0 | 0.91 | 1.00 | 0.95 | 0.93 | 139.6 |
|  |  |  | BugSeq-V2 | 10 | 0 | 0 | 1.00 | 1.00 | 1.00 | 1.00 | 18.1 |
|  | 0.1 percent | Short read | Kraken2 | 10 | 28 | 0 | 0.26 | 1.00 | 0.42 | 0.31 |  |
|  |  |  | Bracken | 10 | 34 | 0 | 0.23 | 1.00 | 0.37 | 0.27 |  |
|  |  |  | Centrifuge-h22 | 8 | 2 | 2 | 0.80 | 0.80 | 0.80 | 0.80 |  |
|  |  |  | Centrifuge-h500 | 6 | 0 | 4 | 1.00 | 0.60 | 0.75 | 0.88 |  |
|  |  |  | Metaphlan3 | 8 | 9 | 2 | 0.47 | 0.80 | 0.59 | 0.51 |  |
|  |  | Long read | Metamaps | 9 | 5 | 1 | 0.64 | 0.90 | 0.75 | 0.68 |  |
|  |  |  | MMseqs2 | 10 | 5 | 0 | 0.67 | 1.00 | 0.80 | 0.71 |  |
|  |  |  | MEGAN-LR-Prot | 10 | 7 | 0 | 0.59 | 1.00 | 0.74 | 0.64 |  |
|  |  |  | MEGAN-LR-Nuc-HiFi | 10 | 1 | 0 | 0.91 | 1.00 | 0.95 | 0.93 |  |
|  |  |  | MEGAN-LR-Nuc-ONT | 10 | 1 | 0 | 0.91 | 1.00 | 0.95 | 0.93 |  |
|  |  |  | BugSeq-V2 | 10 | 0 | 0 | 1.00 | 1.00 | 1.00 | 1.00 |  |
|  | 1 percent | Short read | Kraken2 | 9 | 0 | 1 | 1.00 | 0.90 | 0.95 | 0.98 |  |
|  |  |  | Bracken | 9 | 0 | 1 | 1.00 | 0.90 | 0.95 | 0.98 |  |
|  |  |  | Centrifuge-h22 | 7 | 0 | 3 | 1.00 | 0.70 | 0.82 | 0.92 |  |
|  |  |  | Centrifuge-h500 | 0 | 0 | 10 | 0.00 | 0.00 | 0.00 | 0.00 |  |
|  |  |  | Metaphlan3 | 1 | 0 | 9 | 1.00 | 0.10 | 0.18 | 0.36 |  |
|  |  | Long read | Metamaps | 8 | 0 | 2 | 1.00 | 0.80 | 0.89 | 0.95 |  |
|  |  |  | MMseqs2 | 1 | 0 | 9 | 1.00 | 0.10 | 0.18 | 0.36 |  |
|  |  |  | MEGAN-LR-Prot | 5 | 0 | 5 | 1.00 | 0.50 | 0.67 | 0.83 |  |
|  |  |  | MEGAN-LR-Nuc-HiFi | 6 | 0 | 4 | 1.00 | 0.60 | 0.75 | 0.88 |  |
|  |  |  | MEGAN-LR-Nuc-ONT | 6 | 0 | 4 | 1.00 | 0.60 | 0.75 | 0.88 |  |
|  |  |  | BugSeq-V2 | 9 | 0 | 1 | 1.00 | 0.90 | 0.95 | 0.98 |  |
| ONT Q20 Short | 0.001 percent | Short read | Kraken2 | 10 | 204 | 0 | 0.05 | 1.00 | 0.09 | 0.06 | 72.0 |
|  |  |  | Bracken | 10 | 225 | 0 | 0.04 | 1.00 | 0.08 | 0.05 | 67.0 |
|  |  |  | Centrifuge-h22 | 8 | 58 | 2 | 0.12 | 0.80 | 0.21 | 0.15 | 67.9 |
|  |  |  | Centrifuge-h500 | 8 | 6 | 2 | 0.57 | 0.80 | 0.67 | 0.61 | 55.4 |
|  |  |  | Metaphlan3 | 8 | 29 | 2 | 0.22 | 0.80 | 0.34 | 0.25 | 100.1 |
|  |  | Long read | Metamaps | 9 | 70 | 1 | 0.11 | 0.90 | 0.20 | 0.14 | 36.1 |
|  |  |  | MMseqs2 | 10 | 128 | 0 | 0.07 | 1.00 | 0.13 | 0.09 | 70.2 |
|  |  |  | MEGAN-LR-Prot | 10 | 6 | 0 | 0.63 | 1.00 | 0.77 | 0.68 | 100.0 |
|  |  |  | MEGAN-LR-Nuc-HiFi | 10 | 0 | 0 | 1.00 | 1.00 | 1.00 | 1.00 | 100.0 |
|  |  |  | MEGAN-LR-Nuc-ONT | 10 | 0 | 0 | 1.00 | 1.00 | 1.00 | 1.00 | 100.0 |
|  |  |  | BugSeq-V2 | 10 | 1 | 0 | 0.91 | 1.00 | 0.95 | 0.93 | 45.7 |
|  | 0.1 percent | Short read | Kraken2 | 10 | 30 | 0 | 0.25 | 1.00 | 0.40 | 0.29 |  |
|  |  |  | Bracken | 10 | 43 | 0 | 0.19 | 1.00 | 0.32 | 0.23 |  |
|  |  |  | Centrifuge-h22 | 8 | 11 | 2 | 0.42 | 0.80 | 0.55 | 0.47 |  |
|  |  |  | Centrifuge-h500 | 8 | 0 | 2 | 1.00 | 0.80 | 0.89 | 0.95 |  |
|  |  |  | Metaphlan3 | 8 | 18 | 2 | 0.31 | 0.80 | 0.44 | 0.35 |  |
|  |  | Long read | Metamaps | 9 | 12 | 1 | 0.43 | 0.90 | 0.58 | 0.48 |  |
|  |  |  | MMseqs2 | 10 | 13 | 0 | 0.44 | 1.00 | 0.61 | 0.49 |  |
|  |  |  | MEGAN-LR-Prot | 10 | 6 | 0 | 0.63 | 1.00 | 0.77 | 0.68 |  |
|  |  |  | MEGAN-LR-Nuc-HiFi | 10 | 0 | 0 | 1.00 | 1.00 | 1.00 | 1.00 |  |
|  |  |  | MEGAN-LR-Nuc-ONT | 10 | 0 | 0 | 1.00 | 1.00 | 1.00 | 1.00 |  |
|  |  |  | BugSeq-V2 | 10 | 0 | 0 | 1.00 | 1.00 | 1.00 | 1.00 |  |
|  | 1 percent | Short read | Kraken2 | 10 | 0 | 0 | 1.00 | 1.00 | 1.00 | 1.00 |  |
|  |  |  | Bracken | 10 | 1 | 0 | 0.91 | 1.00 | 0.95 | 0.93 |  |
|  |  |  | Centrifuge-h22 | 7 | 0 | 3 | 1.00 | 0.70 | 0.82 | 0.92 |  |
|  |  |  | Centrifuge-h500 | 6 | 0 | 4 | 1.00 | 0.60 | 0.75 | 0.88 |  |
|  |  |  | Metaphlan3 | 2 | 0 | 8 | 1.00 | 0.20 | 0.33 | 0.56 |  |
|  |  | Long read | Metamaps | 9 | 0 | 1 | 1.00 | 0.90 | 0.95 | 0.98 |  |
|  |  |  | MMseqs2 | 5 | 0 | 5 | 1.00 | 0.50 | 0.67 | 0.83 |  |
|  |  |  | MEGAN-LR-Prot | 4 | 0 | 6 | 1.00 | 0.40 | 0.57 | 0.77 |  |
|  |  |  | MEGAN-LR-Nuc-HiFi | 7 | 0 | 3 | 1.00 | 0.70 | 0.82 | 0.92 |  |
|  |  |  | MEGAN-LR-Nuc-ONT | 7 | 0 | 3 | 1.00 | 0.70 | 0.82 | 0.92 |  |
|  |  |  | BugSeq-V2 | 9 | 0 | 1 | 1.00 | 0.90 | 0.95 | 0.98 |  |

**Supplementary Table S20.** Genus-level detection results for the ONT Short datasets.

| **Dataset** | **Filtering Level** | **Method Type** | **Method** | **True Positives** | **False Positives** | **False Negatives** | **Precision** | **Recall** | **F_1_** | **F_0.5_** |
| --- | --- | --- | --- | --- | --- | --- | --- | --- | --- | --- |
| ONT R10 Short | 0.001 percent | Short read | Kraken2 | 10 | 38 | 0 | 0.21 | 1.00 | 0.34 | 0.25 |
|  |  |  | Bracken | 10 | 26 | 0 | 0.28 | 1.00 | 0.44 | 0.32 |
|  |  |  | Centrifuge-h22 | 8 | 9 | 2 | 0.47 | 0.80 | 0.59 | 0.51 |
|  |  |  | Centrifuge-h500 | 8 | 0 | 2 | 1.00 | 0.80 | 0.89 | 0.95 |
|  |  |  | Metaphlan3 | 10 | 11 | 0 | 0.48 | 1.00 | 0.64 | 0.53 |
|  |  | Long read | Metamaps | 9 | 17 | 1 | 0.35 | 0.90 | 0.50 | 0.39 |
|  |  |  | MMseqs2 | 10 | 51 | 0 | 0.16 | 1.00 | 0.28 | 0.20 |
|  |  |  | MEGAN-LR-Prot | 10 | 7 | 0 | 0.59 | 1.00 | 0.74 | 0.64 |
|  |  |  | MEGAN-LR-Nuc-HiFi | 10 | 1 | 0 | 0.91 | 1.00 | 0.95 | 0.93 |
|  |  |  | MEGAN-LR-Nuc-ONT | 10 | 1 | 0 | 0.91 | 1.00 | 0.95 | 0.93 |
|  |  |  | BugSeq-V2 | 10 | 0 | 0 | 1.00 | 1.00 | 1.00 | 1.00 |
|  | 0.1 percent | Short read | Kraken2 | 10 | 11 | 0 | 0.48 | 1.00 | 0.64 | 0.53 |
|  |  |  | Bracken | 10 | 9 | 0 | 0.53 | 1.00 | 0.69 | 0.58 |
|  |  |  | Centrifuge-h22 | 8 | 2 | 2 | 0.80 | 0.80 | 0.80 | 0.80 |
|  |  |  | Centrifuge-h500 | 7 | 0 | 3 | 1.00 | 0.70 | 0.82 | 0.92 |
|  |  |  | Metaphlan3 | 9 | 3 | 1 | 0.75 | 0.90 | 0.82 | 0.78 |
|  |  | Long read | Metamaps | 9 | 2 | 1 | 0.82 | 0.90 | 0.86 | 0.83 |
|  |  |  | MMseqs2 | 10 | 8 | 0 | 0.56 | 1.00 | 0.71 | 0.61 |
|  |  |  | MEGAN-LR-Prot | 10 | 7 | 0 | 0.59 | 1.00 | 0.74 | 0.64 |
|  |  |  | MEGAN-LR-Nuc-HiFi | 10 | 1 | 0 | 0.91 | 1.00 | 0.95 | 0.93 |
|  |  |  | MEGAN-LR-Nuc-ONT | 10 | 1 | 0 | 0.91 | 1.00 | 0.95 | 0.93 |
|  |  |  | BugSeq-V2 | 10 | 0 | 0 | 1.00 | 1.00 | 1.00 | 1.00 |
|  | 1 percent | Short read | Kraken2 | 9 | 0 | 1 | 1.00 | 0.90 | 0.95 | 0.98 |
|  |  |  | Bracken | 9 | 0 | 1 | 1.00 | 0.90 | 0.95 | 0.98 |
|  |  |  | Centrifuge-h22 | 8 | 0 | 2 | 1.00 | 0.80 | 0.89 | 0.95 |
|  |  |  | Centrifuge-h500 | 0 | 0 | 10 | 0.00 | 0.00 | 0.00 | 0.00 |
|  |  |  | Metaphlan3 | 2 | 0 | 8 | 1.00 | 0.20 | 0.33 | 0.56 |
|  |  | Long read | Metamaps | 8 | 0 | 2 | 1.00 | 0.80 | 0.89 | 0.95 |
|  |  |  | MMseqs2 | 3 | 0 | 7 | 1.00 | 0.30 | 0.46 | 0.68 |
|  |  |  | MEGAN-LR-Prot | 5 | 0 | 5 | 1.00 | 0.50 | 0.67 | 0.83 |
|  |  |  | MEGAN-LR-Nuc-HiFi | 6 | 0 | 4 | 1.00 | 0.60 | 0.75 | 0.88 |
|  |  |  | MEGAN-LR-Nuc-ONT | 6 | 0 | 4 | 1.00 | 0.60 | 0.75 | 0.88 |
|  |  |  | BugSeq-V2 | 9 | 0 | 1 | 1.00 | 0.90 | 0.95 | 0.98 |
| ONT Q20 Short | 0.001 percent | Short read | Kraken2 | 10 | 46 | 0 | 0.18 | 1.00 | 0.30 | 0.21 |
|  |  |  | Bracken | 10 | 35 | 0 | 0.22 | 1.00 | 0.36 | 0.26 |
|  |  |  | Centrifuge-h22 | 8 | 31 | 2 | 0.21 | 0.80 | 0.33 | 0.24 |
|  |  |  | Centrifuge-h500 | 8 | 3 | 2 | 0.73 | 0.80 | 0.76 | 0.74 |
|  |  |  | Metaphlan3 | 10 | 18 | 0 | 0.36 | 1.00 | 0.53 | 0.41 |
|  |  | Long read | Metamaps | 9 | 61 | 1 | 0.13 | 0.90 | 0.23 | 0.16 |
|  |  |  | MMseqs2 | 10 | 65 | 0 | 0.13 | 1.00 | 0.23 | 0.16 |
|  |  |  | MEGAN-LR-Prot | 10 | 5 | 0 | 0.67 | 1.00 | 0.80 | 0.71 |
|  |  |  | MEGAN-LR-Nuc-HiFi | 10 | 0 | 0 | 1.00 | 1.00 | 1.00 | 1.00 |
|  |  |  | MEGAN-LR-Nuc-ONT | 10 | 0 | 0 | 1.00 | 1.00 | 1.00 | 1.00 |
|  |  |  | BugSeq-V2 | 10 | 0 | 0 | 1.00 | 1.00 | 1.00 | 1.00 |
|  | 0.1 percent | Short read | Kraken2 | 10 | 12 | 0 | 0.46 | 1.00 | 0.63 | 0.51 |
|  |  |  | Bracken | 10 | 11 | 0 | 0.48 | 1.00 | 0.64 | 0.53 |
|  |  |  | Centrifuge-h22 | 8 | 4 | 2 | 0.67 | 0.80 | 0.73 | 0.69 |
|  |  |  | Centrifuge-h500 | 8 | 0 | 2 | 1.00 | 0.80 | 0.89 | 0.95 |
|  |  |  | Metaphlan3 | 9 | 10 | 1 | 0.47 | 0.90 | 0.62 | 0.52 |
|  |  | Long read | Metamaps | 9 | 8 | 1 | 0.53 | 0.90 | 0.67 | 0.58 |
|  |  |  | MMseqs2 | 10 | 13 | 0 | 0.44 | 1.00 | 0.61 | 0.49 |
|  |  |  | MEGAN-LR-Prot | 10 | 5 | 0 | 0.67 | 1.00 | 0.80 | 0.71 |
|  |  |  | MEGAN-LR-Nuc-HiFi | 10 | 0 | 0 | 1.00 | 1.00 | 1.00 | 1.00 |
|  |  |  | MEGAN-LR-Nuc-ONT | 10 | 0 | 0 | 1.00 | 1.00 | 1.00 | 1.00 |
|  |  |  | BugSeq-V2 | 10 | 0 | 0 | 1.00 | 1.00 | 1.00 | 1.00 |
|  | 1 percent | Short read | Kraken2 | 10 | 0 | 0 | 1.00 | 1.00 | 1.00 | 1.00 |
|  |  |  | Bracken | 10 | 0 | 0 | 1.00 | 1.00 | 1.00 | 1.00 |
|  |  |  | Centrifuge-h22 | 8 | 0 | 2 | 1.00 | 0.80 | 0.89 | 0.95 |
|  |  |  | Centrifuge-h500 | 7 | 0 | 3 | 1.00 | 0.70 | 0.82 | 0.92 |
|  |  |  | Metaphlan3 | 2 | 0 | 8 | 1.00 | 0.20 | 0.33 | 0.56 |
|  |  | Long read | Metamaps | 9 | 0 | 1 | 1.00 | 0.90 | 0.95 | 0.98 |
|  |  |  | MMseqs2 | 9 | 0 | 1 | 1.00 | 0.90 | 0.95 | 0.98 |
|  |  |  | MEGAN-LR-Prot | 8 | 0 | 2 | 1.00 | 0.80 | 0.89 | 0.95 |
|  |  |  | MEGAN-LR-Nuc-HiFi | 7 | 0 | 3 | 1.00 | 0.70 | 0.82 | 0.92 |
|  |  |  | MEGAN-LR-Nuc-ONT | 7 | 0 | 3 | 1.00 | 0.70 | 0.82 | 0.92 |
|  |  |  | BugSeq-V2 | 10 | 0 | 0 | 1.00 | 1.00 | 1.00 | 1.00 |

**Supplementary Table S21.** Chi-Squared test results for the species-level relative abundances for the ONT Short datasets.

|  | **ONT R10 Short** | | | | **ONT Q20 Short** | | | | | |
| --- | --- | --- | --- | --- | --- | --- | --- | --- | --- | --- |
| **Method** | **df** | **Chi-Squared Statistic** | **Critical Value** | **P-value** | **df** | **Chi-Squared Statistic** | | **Critical Value** | | **P-value** |
| Kraken2 | 10 | 692.05 | 18.31 | **<0.001** | 10 | 1097.90 | 18.31 | | **<0.001** | |
| Bracken | 10 | 775.88 | 18.31 | **<0.001** | 10 | 2447.28 | 18.31 | | **<0.001** | |
| Centrifuge-h22 | 10 | 116.53 | 18.31 | **<0.001** | 10 | 148.39 | 18.31 | | **<0.001** | |
| Centrifuge-h500 | 10 | 145.87 | 18.31 | **<0.001** | 10 | 56.27 | 18.31 | | **<0.001** | |
| Metaphlan3 | 10 | 7382.36 | 18.31 | **<0.001** | 10 | 18498.56 | 18.31 | | **<0.001** | |
| Metamaps | 10 | 116.01 | 18.31 | **<0.001** | 10 | 223.98 | 18.31 | | **<0.001** | |
| MMseqs2 | 10 | 6000.48 | 18.31 | **<0.001** | 10 | 2703.71 | 18.31 | | **<0.001** | |
| MEGAN-LR-Prot | 10 | 684.82 | 18.31 | **<0.001** | 10 | 411.40 | 18.31 | | **<0.001** | |
| MEGAN-LR-Nuc-HiFi | 10 | 458.92 | 18.31 | **<0.001** | 10 | 230.69 | 18.31 | | **<0.001** | |
| MEGAN-LR-Nuc-ONT | 10 | 463.59 | 18.31 | **<0.001** | 10 | 231.33 | 18.31 | | **<0.001** | |
| BugSeq-V2 | 10 | 4.65 | 18.31 | 0.913 | 10 | 50.10 | 18.31 | | **<0.001** | |

The Bonferroni correction for 11 tests sets the alpha to 0.0045, and tests that are significant after the correction are shown in bold. P-values in bold indicate rejection of the null hypothesis that there is no difference between the distributions (signifying the abundance estimates are significantly different).

**Supplementary Table S22.** Chi-Squared test results for the genus-level relative abundances for the ONT Short datasets.

|  | **ONT R10 Short** | | | | **ONT Q20 Short** | | | |
| --- | --- | --- | --- | --- | --- | --- | --- | --- |
| **Method** | **df** | **Chi-Squared Statistic** | **Critical Value** | **P-value** | **df** | **Chi-Squared Statistic** | **Critical Value** | **P-value** |
| Kraken2 | 10 | 333.25 | 18.31 | **<0.001** | 10 | 197.52 | 18.31 | **<0.001** |
| Bracken | 10 | 320.60 | 18.31 | **<0.001** | 10 | 184.75 | 18.31 | **<0.001** |
| Centrifuge-h22 | 10 | 18.11 | 18.31 | 0.053 | 10 | 100.94 | 18.31 | **<0.001** |
| Centrifuge-h500 | 10 | 56.50 | 18.31 | **<0.001** | 10 | 37.73 | 18.31 | **<0.001** |
| Metaphlan3 | 10 | 456.45 | 18.31 | **<0.001** | 10 | 644.52 | 18.31 | **<0.001** |
| Metamaps | 10 | 74.83 | 18.31 | **<0.001** | 10 | 122.05 | 18.31 | **<0.001** |
| MMseqs2 | 10 | 1077.72 | 18.31 | **<0.001** | 10 | 442.06 | 18.31 | **<0.001** |
| MEGAN-LR-Prot | 10 | 451.30 | 18.31 | **<0.001** | 10 | 243.85 | 18.31 | **<0.001** |
| MEGAN-LR-Nuc-HiFi | 10 | 459.80 | 18.31 | **<0.001** | 10 | 228.80 | 18.31 | **<0.001** |
| MEGAN-LR-Nuc-ONT | 10 | 464.12 | 18.31 | **<0.001** | 10 | 228.92 | 18.31 | **<0.001** |
| BugSeq-V2 | 10 | 4.61 | 18.31 | 0.916 | 10 | 29.19 | 18.31 | **0.001** |

The Bonferroni correction for 11 tests sets the alpha to 0.0045, and tests that are significant after the correction are shown in bold. P-values in bold indicate rejection of the null hypothesis that there is no difference between the distributions (signifying the abundance estimates are significantly different).

**Supplementary Table S23.** Species-level detection results for the short-read ATCC datasets.

| **Dataset** | **Filtering Level** | **Method** | **True Positives** | **False Positives** | **False Negatives** | **Precision** | **Recall** | **F1** | **F0.5** | **F1** |
| --- | --- | --- | --- | --- | --- | --- | --- | --- | --- | --- |
| Illumina ATCC MSA1003 | 0.001 percent | Kraken2 | 20 | 77 | 0 | 0.21 | 1.00 | 0.34 | 0.24 | 44.8 |
|  |  | Bracken | 20 | 113 | 0 | 0.15 | 1.00 | 0.26 | 0.18 | 36.4 |
|  |  | Centrifuge-h22 | 20 | 57 | 0 | 0.26 | 1.00 | 0.41 | 0.31 | 54.2 |
|  |  | Metaphlan3 | 12 | 1 | 8 | 0.92 | 0.60 | 0.73 | 0.83 | 12.7 |
|  |  | mOTUs | 4 | 1 | 16 | 0.80 | 0.20 | 0.32 | 0.50 | 51.6 |
|  |  | Sourmash-k31 | 20 | 7 | 0 | 0.74 | 1.00 | 0.85 | 0.78 | 57.2 |
|  |  | Sourmash-k51 | 20 | 5 | 0 | 0.80 | 1.00 | 0.89 | 0.83 | 55.4 |
|  | 0.1 percent | Kraken2 | 11 | 1 | 9 | 0.92 | 0.55 | 0.69 | 0.81 |  |
|  |  | Bracken | 13 | 8 | 7 | 0.62 | 0.65 | 0.63 | 0.62 |  |
|  |  | Centrifuge-h22 | 11 | 1 | 9 | 0.92 | 0.55 | 0.69 | 0.81 |  |
|  |  | Metaphlan3 | 5 | 0 | 15 | 1.00 | 0.25 | 0.40 | 0.63 |  |
|  |  | mOTUs | 0 | 0 | 20 | 0.00 | 0.00 | 0.00 | 0.00 |  |
|  |  | Sourmash-k31 | 20 | 7 | 0 | 0.74 | 1.00 | 0.85 | 0.78 |  |
|  |  | Sourmash-k51 | 20 | 5 | 0 | 0.80 | 1.00 | 0.89 | 0.83 |  |
|  | 1 percent | Kraken2 | 8 | 0 | 12 | 1.00 | 0.40 | 0.57 | 0.77 |  |
|  |  | Bracken | 9 | 1 | 11 | 0.90 | 0.45 | 0.60 | 0.75 |  |
|  |  | Centrifuge-h22 | 7 | 0 | 13 | 1.00 | 0.35 | 0.52 | 0.73 |  |
|  |  | Metaphlan3 | 0 | 0 | 20 | 0 | 0 | 0 | 0 |  |
|  |  | mOTUs | 0 | 0 | 20 | 0 | 0 | 0 | 0 |  |
|  |  | Sourmash-k31 | 20 | 7 | 0 | 0.74 | 1.00 | 0.85 | 0.78 |  |
|  |  | Sourmash-k51 | 20 | 5 | 0 | 0.80 | 1.00 | 0.89 | 0.83 |  |
| SR-Sim ATCC MSA1003 | 0.001 percent | Kraken2 | 20 | 68 | 0 | 0.23 | 1.00 | 0.37 | 0.27 | 67.4 |
|  |  | Bracken | 20 | 95 | 0 | 0.17 | 1.00 | 0.30 | 0.21 | 53.4 |
|  |  | Centrifuge-h22 | 20 | 33 | 0 | 0.38 | 1.00 | 0.55 | 0.43 | 75.6 |
|  |  | Metaphlan3 | 12 | 1 | 8 | 0.92 | 0.60 | 0.73 | 0.83 | 24.4 |
|  |  | mOTUs* | 12 | 1 | 8 | 0.92 | 0.60 | 0.73 | 0.83 |  |
|  |  | Sourmash-k31 | 20 | 3 | 0 | 0.87 | 1.00 | 0.93 | 0.89 | 66.7 |
|  |  | Sourmash-k51 | 20 | 2 | 0 | 0.91 | 1.00 | 0.95 | 0.93 | 65.3 |
|  | 0.1 percent | Kraken2 | 12 | 0 | 8 | 1.00 | 0.60 | 0.75 | 0.88 |  |
|  |  | Bracken | 13 | 7 | 7 | 0.65 | 0.65 | 0.65 | 0.65 |  |
|  |  | Centrifuge-h22 | 11 | 1 | 9 | 0.92 | 0.55 | 0.69 | 0.81 |  |
|  |  | Metaphlan3 | 5 | 0 | 15 | 1.00 | 0.25 | 0.40 | 0.63 |  |
|  |  | mOTUs* | 4 | 0 | 16 | 1.00 | 0.20 | 0.33 | 0.56 |  |
|  |  | Sourmash-k31 | 20 | 3 | 0 | 0.87 | 1.00 | 0.93 | 0.89 |  |
|  |  | Sourmash-k51 | 20 | 2 | 0 | 0.91 | 1.00 | 0.95 | 0.93 |  |
|  | 1 percent | Kraken2 | 6 | 0 | 14 | 1.00 | 0.30 | 0.46 | 0.68 |  |
|  |  | Bracken | 8 | 1 | 12 | 0.89 | 0.40 | 0.55 | 0.71 |  |
|  |  | Centrifuge-h22 | 7 | 0 | 13 | 1.00 | 0.35 | 0.52 | 0.73 |  |
|  |  | Metaphlan3 | 0 | 0 | 20 | 0 | 0 | 0 | 0 |  |
|  |  | mOTUs* | 0 | 0 | 20 | 0 | 0 | 0 | 0 |  |
|  |  | Sourmash-k31 | 17 | 2 | 3 | 0.90 | 0.85 | 0.87 | 0.89 |  |
|  |  | Sourmash-k51 | 17 | 1 | 3 | 0.94 | 0.85 | 0.89 | 0.92 |  |

*Results for mOTUs are from the main analysis, as the mOTUS long read mode breaks long reads into suitably sized short reads, as does our short-read simulation analysis.

**Supplementary Table S24.** Genus-level detection results for the short-read ATCC datasets.

| **Dataset** | **Filtering Level** | **Method** | **True Positives** | **False Positives** | **False Negatives** | **Precision** | **Recall** | **F1** | **F0.5** |
| --- | --- | --- | --- | --- | --- | --- | --- | --- | --- |
| Illumina ATCC MSA1003 | 0.001 percent | Kraken2 | 18 | 31 | 0 | 0.37 | 1.00 | 0.54 | 0.42 |
|  |  | Bracken | 18 | 34 | 0 | 0.35 | 1.00 | 0.51 | 0.40 |
|  |  | Centrifuge-h22 | 18 | 41 | 0 | 0.31 | 1.00 | 0.47 | 0.35 |
|  |  | Metaphlan3 | 11 | 0 | 7 | 1.00 | 0.61 | 0.76 | 0.89 |
|  |  | mOTUs | 5 | 0 | 13 | 1.00 | 0.28 | 0.44 | 0.66 |
|  |  | Sourmash-k31 | 18 | 4 | 0 | 0.82 | 1.00 | 0.90 | 0.85 |
|  |  | Sourmash-k51 | 18 | 3 | 0 | 0.86 | 1.00 | 0.92 | 0.88 |
|  | 0.1 percent | Kraken2 | 11 | 1 | 7 | 0.92 | 0.61 | 0.73 | 0.83 |
|  |  | Bracken | 11 | 3 | 7 | 0.79 | 0.61 | 0.69 | 0.74 |
|  |  | Centrifuge-h22 | 11 | 0 | 7 | 1.00 | 0.61 | 0.76 | 0.89 |
|  |  | Metaphlan3 | 5 | 0 | 13 | 1.00 | 0.28 | 0.44 | 0.66 |
|  |  | mOTUs | 0 | 0 | 18 | 0 | 0 | 0 | 0 |
|  |  | Sourmash-k31 | 18 | 4 | 0 | 0.82 | 1.00 | 0.90 | 0.85 |
|  |  | Sourmash-k51 | 18 | 3 | 0 | 0.86 | 1.00 | 0.92 | 0.88 |
|  | 1 percent | Kraken2 | 7 | 0 | 11 | 1.00 | 0.39 | 0.56 | 0.76 |
|  |  | Bracken | 8 | 1 | 10 | 0.89 | 0.44 | 0.59 | 0.74 |
|  |  | Centrifuge-h22 | 7 | 0 | 11 | 1.00 | 0.39 | 0.56 | 0.76 |
|  |  | Metaphlan3 | 0 | 0 | 18 | 0 | 0 | 0 | 0 |
|  |  | mOTUs | 0 | 0 | 18 | 0 | 0 | 0 | 0 |
|  |  | Sourmash-k31 | 16 | 4 | 2 | 0.80 | 0.89 | 0.84 | 0.82 |
|  |  | Sourmash-k51 | 15 | 3 | 3 | 0.83 | 0.83 | 0.83 | 0.83 |
| SR-Sim ATCC MSA1003 | 0.001 percent | Kraken2 | 18 | 20 | 0 | 0.47 | 1.00 | 0.64 | 0.53 |
|  |  | Bracken | 18 | 23 | 0 | 0.44 | 1.00 | 0.61 | 0.49 |
|  |  | Centrifuge-h22 | 18 | 7 | 0 | 0.72 | 1.00 | 0.84 | 0.76 |
|  |  | Metaphlan3 | 11 | 0 | 7 | 1.00 | 0.61 | 0.76 | 0.89 |
|  |  | mOTUs* | 11 | 0 | 7 | 1.00 | 0.61 | 0.76 | 0.89 |
|  |  | Sourmash-k31 | 18 | 1 | 0 | 0.95 | 1.00 | 0.97 | 0.96 |
|  |  | Sourmash-k51 | 18 | 1 | 0 | 0.95 | 1.00 | 0.97 | 0.96 |
|  | 0.1 percent | Kraken2 | 11 | 1 | 7 | 0.92 | 0.61 | 0.73 | 0.83 |
|  |  | Bracken | 11 | 2 | 7 | 0.85 | 0.61 | 0.71 | 0.79 |
|  |  | Centrifuge-h22 | 11 | 0 | 7 | 1.00 | 0.61 | 0.76 | 0.89 |
|  |  | Metaphlan3 | 5 | 0 | 13 | 1.00 | 0.28 | 0.44 | 0.66 |
|  |  | mOTUs* | 4 | 0 | 14 | 1.00 | 0.22 | 0.36 | 0.59 |
|  |  | Sourmash-k31 | 18 | 1 | 0 | 0.95 | 1.00 | 0.97 | 0.96 |
|  |  | Sourmash-k51 | 18 | 1 | 0 | 0.95 | 1.00 | 0.97 | 0.96 |
|  | 1 percent | Kraken2 | 8 | 0 | 10 | 1.00 | 0.44 | 0.61 | 0.80 |
|  |  | Bracken | 8 | 1 | 10 | 0.89 | 0.44 | 0.59 | 0.74 |
|  |  | Centrifuge-h22 | 8 | 0 | 10 | 1.00 | 0.44 | 0.61 | 0.80 |
|  |  | Metaphlan3 | 1 | 0 | 17 | 1.00 | 0.06 | 0.11 | 0.23 |
|  |  | mOTUs* | 0 | 0 | 18 | 0 | 0 | 0 | 0 |
|  |  | Sourmash-k31 | 15 | 0 | 3 | 1.00 | 0.83 | 0.91 | 0.96 |
|  |  | Sourmash-k51 | 15 | 0 | 3 | 1.00 | 0.83 | 0.91 | 0.96 |

*Results for mOTUs are from the main analysis, as the mOTUS long read mode breaks long reads into suitably sized short reads, as does our short-read simulation analysis.

**Supplementary Table S25.** Chi-Squared test results for the species-level relative abundances for the short-read ATCC datasets.

|  | **Illumina ATCC MSA1003** | | | | **SR-Sim ATCC MSA1003** | | | |
| --- | --- | --- | --- | --- | --- | --- | --- | --- |
| **Method** | **df** | **Chi-Squared Statistic** | **Critical Value** | **P-value** | **df** | **Chi-Squared Statistic** | **Critical Value** | **P-value** |
| Kraken2 | 20 | 177.09 | 31.41 | **<0.001** | 20 | 115.16 | 31.41 | **<0.001** |
| Bracken | 20 | 1372.10 | 31.41 | **<0.001** | 20 | 1228.57 | 31.41 | **<0.001** |
| Centrifuge-h22 | 20 | 126.61 | 31.41 | **<0.001** | 20 | 133.65 | 31.41 | **<0.001** |
| Metaphlan3 | 20 | 213.41 | 31.41 | **<0.001** | 20 | 33.8 | 31.41 | 0.028 |
| mOTUs* | 20 | 1779.90 | 31.41 | **<0.001** | 20 | 821.56 | 31.41 | **<0.001** |
| Sourmash-k31 | 20 | 17962.47 | 31.41 | **<0.001** | 20 | 8632.23 | 31.41 | **<0.001** |
| Sourmash-k51 | 20 | 15650.98 | 31.41 | **<0.001** | 20 | 7821.87 | 31.41 | **<0.001** |

The Bonferroni correction for 7 tests sets the alpha to 0.007, and tests that are significant after the correction are shown in bold. P-values in bold indicate rejection of the null hypothesis that there is no difference between the distributions (signifying the abundance estimates are significantly different). *Results for mOTUs for SR-Sim are from the main analysis, as the mOTUS long read mode breaks long reads into suitably sized short reads, as does our short-read simulation analysis.

**Supplementary Table S26.** Chi-Squared test results for the genus-level relative abundances for the short-read ATCC datasets.

|  | **Illumina ATCC MSA1003** | | | | **SR-Sim ATCC MSA1003** | | | |
| --- | --- | --- | --- | --- | --- | --- | --- | --- |
| **Method** | **df** | **Chi-Squared Statistic** | **Critical Value** | **P-value** | **df** | **Chi-Squared Statistic** | **Critical Value** | **P-value** |
| Kraken2 | 18 | 96.02 | 28.87 | **<0.001** | 18 | 80.46 | 28.87 | **<0.001** |
| Bracken | 18 | 698.21 | 28.87 | **<0.001** | 18 | 766.53 | 28.87 | **<0.001** |
| Centrifuge-h22 | 18 | 55.63 | 28.87 | **<0.001** | 18 | 71.91 | 28.87 | **<0.001** |
| Metaphlan3 | 18 | 8.12 | 28.87 | 0.977 | 18 | 43.74 | 28.87 | **0.001** |
| mOTUs* | 18 | 30.31 | 28.87 | **<0.001** | 18 | 31.13 | 28.87 | 0.028 |
| Sourmash-k31 | 18 | 101.4 | 28.87 | **<0.001** | 18 | 101.4 | 28.87 | 0.054 |
| Sourmash-k51 | 18 | 26.16 | 28.87 | 0.096 | 18 | 26.16 | 28.87 | 0.075 |

The Bonferroni correction for 7 tests sets the alpha to 0.007, and tests that are significant after the correction are shown in bold. P-values in bold indicate rejection of the null hypothesis that there is no difference between the distributions (signifying the abundance estimates are significantly different). *Results for mOTUs for SR-Sim are from the main analysis, as the mOTUS long read mode breaks long reads into suitably sized short reads, as does our short-read simulation analysis.

**Supplementary Table S27.** Species-level detection results for the short-read Zymo D6300 datasets.

| **Dataset** | **Filtering Level** | **Method** | **True Positives** | **False Positives** | **False Negatives** | **Precision** | **Recall** | **F1** | **F0.5** | **F1** |
| --- | --- | --- | --- | --- | --- | --- | --- | --- | --- | --- |
| Illumina Zymo D6300 | 0.001 percent | Kraken2 | 10 | 62 | 0 | 0.14 | 1.00 | 0.24 | 0.17 | 59.7 |
|  |  | Bracken | 10 | 96 | 0 | 0.09 | 1.00 | 0.17 | 0.11 | 80.0 |
|  |  | Centrifuge-h22 | 8 | 27 | 2 | 0.23 | 0.80 | 0.36 | 0.27 | 81.7 |
|  |  | Metaphlan3 | 7 | 1 | 3 | 0.88 | 0.70 | 0.78 | 0.83 | 82.7 |
|  |  | mOTUs | 7 | 0 | 3 | 1.00 | 0.70 | 0.82 | 0.92 | 55.2 |
|  |  | Sourmash-k31 | 10 | 2 | 0 | 0.83 | 1.00 | 0.91 | 0.86 | 99.3 |
|  |  | Sourmash-k51 | 9 | 2 | 1 | 0.82 | 0.90 | 0.86 | 0.83 | 82.3 |
|  | 0.1 percent | Kraken2 | 9 | 0 | 1 | 1.00 | 0.90 | 0.95 | 0.98 |  |
|  |  | Bracken | 10 | 3 | 0 | 0.77 | 1.00 | 0.87 | 0.81 |  |
|  |  | Centrifuge-h22 | 7 | 0 | 3 | 1.00 | 0.70 | 0.82 | 0.92 |  |
|  |  | Metaphlan3 | 5 | 0 | 5 | 1.00 | 0.50 | 0.67 | 0.83 |  |
|  |  | mOTUs | 0 | 0 | 10 | 0 | 0 | 0 | 0 |  |
|  |  | Sourmash-k31 | 10 | 2 | 0 | 0.83 | 1.00 | 0.91 | 0.86 |  |
|  |  | Sourmash-k51 | 9 | 2 | 1 | 0.82 | 0.90 | 0.86 | 0.83 |  |
|  | 1 percent | Kraken2 | 7 | 0 | 3 | 1.00 | 0.70 | 0.82 | 0.92 |  |
|  |  | Bracken | 8 | 1 | 2 | 0.89 | 0.80 | 0.84 | 0.87 |  |
|  |  | Centrifuge-h22 | 7 | 0 | 3 | 1.00 | 0.70 | 0.82 | 0.92 |  |
|  |  | Metaphlan3 | 0 | 0 | 10 | 0 | 0 | 0 | 0 |  |
|  |  | mOTUs | 0 | 0 | 10 | 0 | 0 | 0 | 0 |  |
|  |  | Sourmash-k31 | 10 | 1 | 0 | 0.91 | 1.00 | 0.95 | 0.93 |  |
|  |  | Sourmash-k51 | 9 | 1 | 1 | 0.90 | 0.90 | 0.90 | 0.90 |  |
| SR-Sim ZymoD6300 | 0.001 percent | Kraken2 | 10 | 146 | 0 | 0.06 | 1.00 | 0.12 | 0.08 | 59.0 |
|  |  | Bracken | 10 | 190 | 0 | 0.05 | 1.00 | 0.10 | 0.06 | 27.5 |
|  |  | Centrifuge-h22 | 8 | 63 | 2 | 0.11 | 0.80 | 0.20 | 0.14 | 51.7 |
|  |  | Metaphlan3 | 7 | 1 | 3 | 0.88 | 0.70 | 0.78 | 0.83 | 82.7 |
|  |  | mOTUs* | 9 | 0 | 6 | 1 | 0.6 | 0.75 | 0.88 |  |
|  |  | Sourmash-k31 | 10 | 6 | 0 | 0.63 | 1.00 | 0.77 | 0.68 | 61.7 |
|  |  | Sourmash-k51 | 10 | 1 | 0 | 0.91 | 1.00 | 0.95 | 0.93 | 22.5 |
|  | 0.1 percent | Kraken2 | 10 | 2 | 0 | 0.83 | 1.00 | 0.91 | 0.86 |  |
|  |  | Bracken | 10 | 16 | 0 | 0.39 | 1.00 | 0.56 | 0.44 |  |
|  |  | Centrifuge-h22 | 8 | 0 | 2 | 1.00 | 0.80 | 0.89 | 0.95 |  |
|  |  | Metaphlan3 | 5 | 0 | 5 | 1.00 | 0.50 | 0.67 | 0.83 |  |
|  |  | mOTUs* | 0 | 0 | 10 | 0 | 0 | 0 | 0 |  |
|  |  | Sourmash-k31 | 10 | 6 | 0 | 0.63 | 1.00 | 0.77 | 0.68 |  |
|  |  | Sourmash-k51 | 10 | 1 | 0 | 0.91 | 1.00 | 0.95 | 0.93 |  |
|  | 1 percent | Kraken2 | 9 | 0 | 1 | 1.00 | 0.90 | 0.95 | 0.98 |  |
|  |  | Bracken | 10 | 1 | 0 | 0.91 | 1.00 | 0.95 | 0.93 |  |
|  |  | Centrifuge-h22 | 7 | 0 | 3 | 1.00 | 0.70 | 0.82 | 0.92 |  |
|  |  | Metaphlan3 | 0 | 0 | 10 | 0 | 0 | 0 | 0 |  |
|  |  | mOTUs* | 0 | 0 | 10 | 0 | 0 | 0 | 0 |  |
|  |  | Sourmash-k31 | 9 | 4 | 1 | 0.69 | 0.90 | 0.78 | 0.73 |  |
|  |  | Sourmash-k51 | 10 | 0 | 0 | 1.00 | 1.00 | 1.00 | 1.00 |  |

*Results for mOTUs are from the main analysis, as the mOTUS long read mode breaks long reads into suitably sized short reads, as does our short-read simulation analysis.

**Supplementary Table S28.** Genus-level detection results for the short-read Zymo D6300 datasets.

| **Dataset** | **Filtering Level** | **Method** | **True Positives** | **False Positives** | **False Negatives** | **Precision** | **Recall** | **F1** | **F0.5** |
| --- | --- | --- | --- | --- | --- | --- | --- | --- | --- |
| Illumina Zymo D6300 | 0.001 percent | Kraken2 | 10 | 19 | 0 | 0.35 | 1.00 | 0.51 | 0.40 |
|  |  | Bracken | 10 | 20 | 0 | 0.33 | 1.00 | 0.50 | 0.38 |
|  |  | Centrifuge-h22 | 8 | 11 | 2 | 0.42 | 0.80 | 0.55 | 0.47 |
|  |  | Metaphlan3 | 9 | 0 | 1 | 1.00 | 0.90 | 0.95 | 0.98 |
|  |  | mOTUs | 7 | 0 | 3 | 1.00 | 0.70 | 0.82 | 0.92 |
|  |  | Sourmash-k31 | 10 | 0 | 0 | 1.00 | 1.00 | 1.00 | 1.00 |
|  |  | Sourmash-k51 | 10 | 0 | 0 | 1.00 | 1.00 | 1.00 | 1.00 |
|  | 0.1 percent | Kraken2 | 10 | 0 | 0 | 1.00 | 1.00 | 1.00 | 1.00 |
|  |  | Bracken | 10 | 1 | 0 | 0.91 | 1.00 | 0.95 | 0.93 |
|  |  | Centrifuge-h22 | 8 | 0 | 2 | 1.00 | 0.80 | 0.89 | 0.95 |
|  |  | Metaphlan3 | 6 | 0 | 4 | 1.00 | 0.60 | 0.75 | 0.88 |
|  |  | mOTUs | 0 | 0 | 10 | 0 | 0 | 0 | 0 |
|  |  | Sourmash-k31 | 10 | 0 | 0 | 1.00 | 1.00 | 1.00 | 1.00 |
|  |  | Sourmash-k51 | 10 | 0 | 0 | 1.00 | 1.00 | 1.00 | 1.00 |
|  | 1 percent | Kraken2 | 8 | 0 | 2 | 1.00 | 0.80 | 0.89 | 0.95 |
|  |  | Bracken | 8 | 0 | 2 | 1.00 | 0.80 | 0.89 | 0.95 |
|  |  | Centrifuge-h22 | 8 | 0 | 2 | 1.00 | 0.80 | 0.89 | 0.95 |
|  |  | Metaphlan3 | 0 | 0 | 10 | 0 | 0 | 0 | 0 |
|  |  | mOTUs | 0 | 0 | 10 | 0 | 0 | 0 | 0 |
|  |  | Sourmash-k31 | 10 | 0 | 0 | 1.00 | 1.00 | 1.00 | 1.00 |
|  |  | Sourmash-k51 | 10 | 0 | 0 | 1.00 | 1.00 | 1.00 | 1.00 |
| SR-Sim ZymoD6300 | 0.001 percent | Kraken2 | 10 | 44 | 0 | 0.19 | 1.00 | 0.31 | 0.22 |
|  |  | Bracken | 10 | 45 | 0 | 0.18 | 1.00 | 0.31 | 0.22 |
|  |  | Centrifuge-h22 | 8 | 49 | 2 | 0.14 | 0.80 | 0.24 | 0.17 |
|  |  | Metaphlan3 | 9 | 0 | 1 | 1.00 | 0.90 | 0.95 | 0.98 |
|  |  | mOTUs* | 8 | 0 | 2 | 1 | 0.8 | 0.89 | 0.95 |
|  |  | Sourmash-k31 | 10 | 1 | 0 | 0.91 | 1.00 | 0.95 | 0.93 |
|  |  | Sourmash-k51 | 10 | 0 | 0 | 1.00 | 1.00 | 1.00 | 1.00 |
|  | 0.1 percent | Kraken2 | 10 | 2 | 0 | 0.83 | 1.00 | 0.91 | 0.86 |
|  |  | Bracken | 10 | 2 | 0 | 0.83 | 1.00 | 0.91 | 0.86 |
|  |  | Centrifuge-h22 | 8 | 0 | 2 | 1.00 | 0.80 | 0.89 | 0.95 |
|  |  | Metaphlan3 | 6 | 0 | 4 | 1.00 | 0.60 | 0.75 | 0.88 |
|  |  | mOTUs* | 0 | 0 | 10 | 0 | 0 | 0 | 0 |
|  |  | Sourmash-k31 | 10 | 1 | 0 | 0.91 | 1.00 | 0.95 | 0.93 |
|  |  | Sourmash-k51 | 10 | 0 | 0 | 1.00 | 1.00 | 1.00 | 1.00 |
|  | 1 percent | Kraken2 | 10 | 0 | 0 | 1.00 | 1.00 | 1.00 | 1.00 |
|  |  | Bracken | 10 | 0 | 0 | 1.00 | 1.00 | 1.00 | 1.00 |
|  |  | Centrifuge-h22 | 8 | 0 | 2 | 1.00 | 0.80 | 0.89 | 0.95 |
|  |  | Metaphlan3 | 0 | 0 | 10 | 0 | 0 | 0 | 0 |
|  |  | mOTUs* | 0 | 0 | 10 | 0 | 0 | 0 | 0 |
|  |  | Sourmash-k31 | 10 | 1 | 0 | 0.91 | 1.00 | 0.95 | 0.93 |
|  |  | Sourmash-k51 | 10 | 0 | 0 | 1.00 | 1.00 | 1.00 | 1.00 |

*Results for mOTUs are from the main analysis, as the mOTUS long read mode breaks long reads into suitably sized short reads, as does our short-read simulation analysis.

**Supplementary Table S29.** Chi-Squared test results for the species abundances for the short-read Zymo D6300 datasets.

|  | **Illumina Zymo D6300** | | | | **SR-Sim ZymoD6300** | | | |
| --- | --- | --- | --- | --- | --- | --- | --- | --- |
| **Method** | **df** | **Chi-Squared Statistic** | **Critical Value** | **P-value** | **df** | **Chi-Squared Statistic** | **Critical Value** | **P-value** |
| Kraken2 | 10 | 150.06 | 18.31 | **<0.001** | 10 | 489.76 | 18.31 | **<0.001** |
| Bracken | 10 | 940.18 | 18.31 | **<0.001** | 10 | 7609.48 | 18.31 | **<0.001** |
| Centrifuge-h22 | 10 | 105.08 | 18.31 | **<0.001** | 10 | 183.08 | 18.31 | **<0.001** |
| Metaphlan3 | 10 | 1121.18 | 18.31 | **<0.001** | 10 | 1121.18 | 18.31 | **<0.001** |
| mOTUs* | 10 | 42.01 | 18.31 | **<0.001** | 10 | 93.11 | 18.31 | **<0.001** |
| Sourmash-k31 | 10 | 161.09 | 18.31 | **<0.001** | 10 | 18003.55 | 18.31 | **<0.001** |
| Sourmash-k51 | 10 | 899.49 | 18.31 | **<0.001** | 10 | 8.88 | 18.31 | 0.544 |

The Bonferroni correction for 7 tests sets the alpha to 0.007, and tests that are significant after the correction are shown in bold. P-values in bold indicate rejection of the null hypothesis that there is no difference between the distributions (signifying the abundance estimates are significantly different). *Results for mOTUs for SR-Sim are from the main analysis, as the mOTUS long read mode breaks long reads into suitably sized short reads, as does our short-read simulation analysis.

**Supplementary Table S30.** Chi-Squared test results for the genus abundances for the short-read Zymo D6300 datasets.

|  | **Illumina Zymo D6300** | | | | **SR-Sim ZymoD6300** | | | |
| --- | --- | --- | --- | --- | --- | --- | --- | --- |
| **Method** | **df** | **Chi-Squared Statistic** | **Critical Value** | **P-value** | **df** | **Chi-Squared Statistic** | **Critical Value** | **P-value** |
| Kraken2 | 10 | 99.42 | 18.31 | **<0.001** | 10 | 57.16 | 18.31 | **<0.001** |
| Bracken | 10 | 206.34 | 18.31 | **<0.001** | 10 | 156.43 | 18.31 | **<0.001** |
| Centrifuge-h22 | 10 | 90.32 | 18.31 | **<0.001** | 10 | 37.40 | 18.31 | **<0.001** |
| Metaphlan3 | 10 | 53.68 | 18.31 | **<0.001** | 10 | 53.68 | 18.31 | **<0.001** |
| mOTUs* | 10 | 41.88 | 18.31 | **<0.001** | 10 | 92.95 | 18.31 | **<0.001** |
| Sourmash-k31 | 10 | 160.19 | 18.31 | **<0.001** | 10 | 32.99 | 18.31 | **<0.001** |
| Sourmash-k51 | 10 | 66.66 | 18.31 | **<0.001** | 10 | 8.9 | 18.31 | 0.542 |

The Bonferroni correction for 7 tests sets the alpha to 0.007, and tests that are significant after the correction are shown in bold. P-values in bold indicate rejection of the null hypothesis that there is no difference between the distributions (signifying the abundance estimates are significantly different). *Results for mOTUs for SR-Sim are from the main analysis, as the mOTUS long read mode breaks long reads into suitably sized short reads, as does our short-read simulation analysis.
